# TAF2 condensation in nuclear speckles links basal transcription factor TFIID to RNA splicing

**DOI:** 10.1101/2024.02.05.578926

**Authors:** Tanja Bhuiyan, Paulina K. Mendoza Sanchez, Niccolò Arecco, Juhyeong Kim, Sheikh Nizamuddin, Andrea Prunotto, Mehmet Tekman, Martin L. Biniossek, Stefanie Koidl, Thorsten Hugel, Sebastian J. Arnold

## Abstract

TFIID is an essential basal transcription factor, crucial for RNA polymerase II (pol II) promoter recognition and transcription initiation. The TFIID complex consists of the TATA-binding protein (TBP) and 13 TBP-associated factors (TAFs) that contain intrinsically disordered regions (IDRs) with currently unknown functions. Here, we show that a conserved IDR drives TAF2 condensation in nuclear speckles, independently of other TFIID subunits. Quantitative mass spectrometry analyses reveal that the TAF2 IDR specifically interacts with the nuclear speckle and spliceosome-associated protein SRRM2. Consequently, TAF2 recruits SRRM2 to TFIID to form non-canonical TFIID-SRRM2 complexes. Reduced SRRM2 recruitment elicits alternative splicing events in RNAs coding for proteins involved in transcription and transmembrane transport. Further, genome-wide binding analyses suggest TAF2 shuttling between nuclear speckles and pol II promoters. This study identifies an IDR of the basal transcription machinery as a molecular guide for protein partitioning into nuclear compartments, controlling protein complex composition and pre-mRNA splicing.

## Introduction

In eukaryotes, the transcription of the vast majority of genes is performed by RNA polymerase II (pol II), which requires accessory factors for different steps of transcription (reviewed in^1^). TFIID is an essential basal transcription factor involved in the rate-limiting first step of transcription initiation at pol II promoters (reviewed in^2,3^). TFIID recognizes and binds pol II core promoters and facilitates the formation of pre-initiation complexes (PICs), which include TFIIA, TFIIB, TFIIE, TFIIF, TFIIH, and pol II^4–6^. Active transcription further depends on the Mediator complex, which interacts with the PIC and imparts enhancer-promoter gene looping (reviewed in^7^). Human TFIID is a large multiprotein complex consisting of the TATA-binding protein (TBP) and 13 TBP-associated factors called TAFs (reviewed in^8^). Cryogenic electron microscopy (cryo-EM) data^9–11^ revealed that nuclear TFIID exists as a three-lobed stable complex termed holo-TFIID (Fig. 1A). Individual TAFs regulate different aspects of TFIID function, such as TFIID complex assembly (TAF1^12^), TFIID complex integrity (TAF5/6/9^13^), and promoter recognition (TAF1/2/4/7^10,11,14^). Expectedly, TAFs are essential genes as gene deletions in mice are embryonically lethal^15–17^.

**Figure 1:**
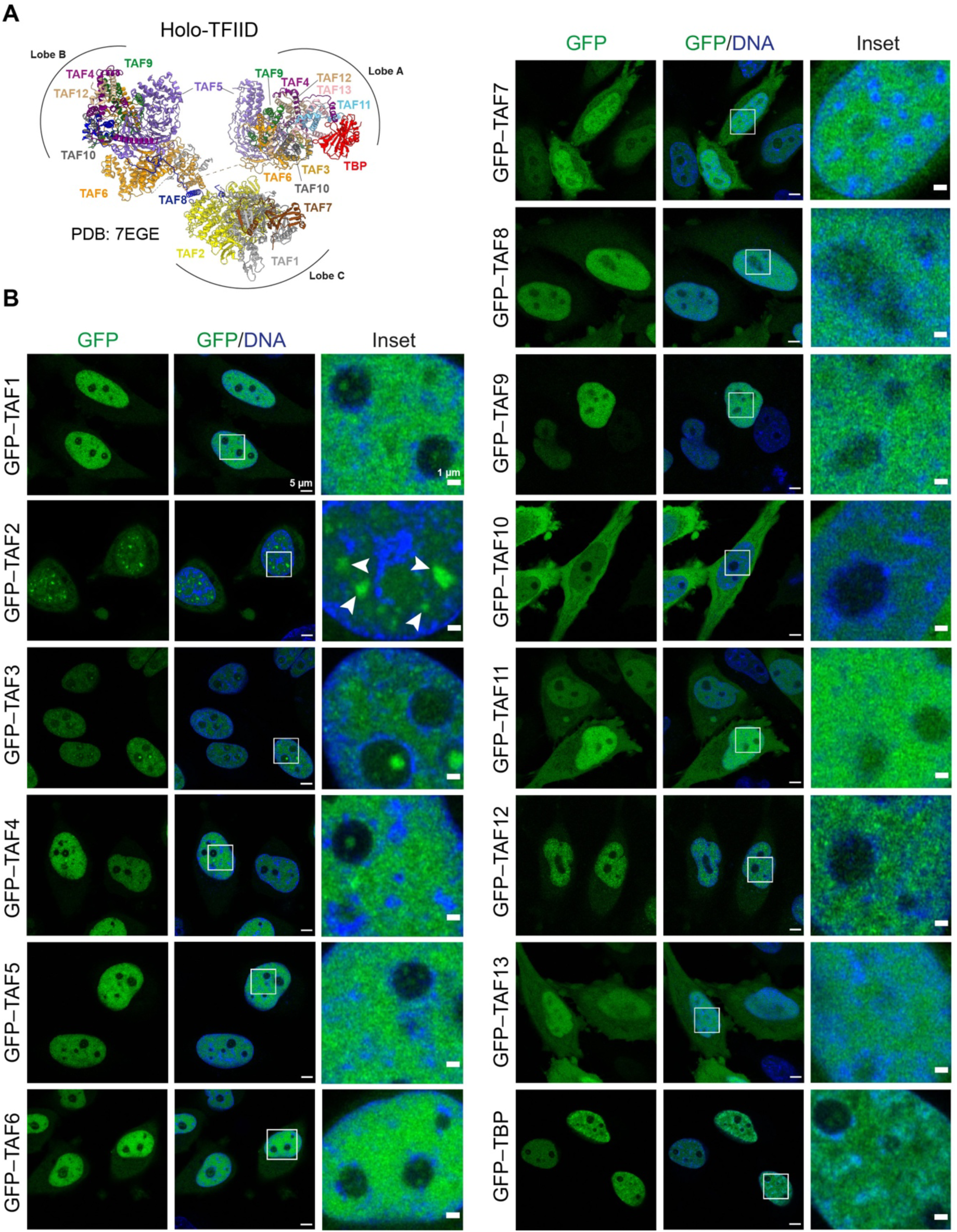
Cellular distribution of individual TFIID subunits. **(A)** Cryo-EM structure of human holo-TFIID, protein data bank (PDB) structure code 7EGE^11^. **(B)** Live-cell confocal microscopy (AiryScan) images of HeLa cells expressing individual TFIID subunits as N-terminal GFP fusions under the control of a doxycycline-inducible promoter. Single confocal planes are shown. DNA was stained with Hoechst 33342. White arrow heads point to nuclear speckles. Scale bars, 5 µm and 1 µm (inset). See also Figs. S1A and B.

TAF2 (formerly TAFII150 or CIF) is an integral part of TFIID and participates in the recognition of core promoter DNA^18–20^ through its structured aminopeptidase M1-like domain^11,14,21^, which lacks peptidase activity^22^. The TAF2 aminopeptidase-like domain interacts with TAF1 to form a DNA-binding subcomplex with TAF7^11^ for promoter recognition. TAF2 also interacts with TAF8, which provides stability to the TFIID complex^10,11,23^. Human mutations in the aminopeptidase-like domain are associated with intellectual disability syndromes^24–26^ and, moreover, *TAF2* is co-amplified with *MYC* in breast carcinomas^27^. Several lines of evidence suggest that TAF2 might be involved in cellular processes independent of TFIID. Notably, TAF2 is only substoichiometrically associated with the TFIID complex as shown by size-exclusion chromatography^19,28^ and quantitative mass spectrometry^12,13^. Further, TAF2 contains a C-terminal intrinsically disordered region (IDR), which interacts non-specifically with promoter DNA^29^, suggesting that the IDR participates in TAF2-DNA interactions. Further, a recent cryo-EM study showed that TAF2 dissociation from promoter DNA and subsequent binding to the TFIIH core is necessary to allow for later steps during transcription initiation, such as promoter melting^11^. Together, these data raise the intriguing possibility that TAF2 might be partially sequestered in other protein complexes or nuclear compartments.

IDRs are common features of transcriptional and chromatin regulators^30^ and are predicted to make up *ca*. 30% of the human proteome^31^. These regions typically exhibit low evolutionary constraints on their primary amino acid sequence, but specific molecular features, such as their physicochemical properties are often found to be evolutionarily conserved^32^. IDRs can drive phase transitions and thus influence the localization of proteins in different membraneless compartments, such as nuclear bodies or other types of biomolecular condensates including transcriptional condensates (reviewed in^33,34^). Phase separation mechanisms driven by IDRs have been implicated in almost all steps along the transcription cycle: coactivator activity of the Mediator complex at super enhancers^35^, recruitment of Mediator and RNA polymerase II^36^, promoter-proximal pausing^37^, or transcription elongation^38,39^. In addition to TAF2, several other TFIID subunits contain IDRs^40^, but the relevance of these domains for transcription or nuclear compartmentalization is unclear.

Here, we demonstrate that the TAF2 IDR partitions a TAF2 protein pool into nuclear speckles, independently of other TFIID subunits. Combining high-resolution imaging with quantitative mass spectrometry, we find that TAF2 interacts with the splicing factor SRRM2. We show that the TAF2 IDR recruits SRRM2 to the TFIID complex and thereby influences pre-mRNA splicing. This study thus suggests a direct link between basal transcription factors and RNA splicing.

## Results

### TAF2 accumulates in nuclear speckles in human cells

Members of the pre-initiation complex (PIC) can form biomolecular condensates *in vitro* during reconstituted mammalian transcription^41,42^. To understand if TFIID can form biomolecular condensates in cells, we explored the nuclear distribution patterns of all individual TFIID subunits. To this end, we inducibly expressed each TFIID subunit fused to GFP under the control of a tet-inducible promoter in HeLa cells (Flp-In T-Rex system^12,13^). Live-cell confocal high-resolution imaging revealed a range of different cellular distribution patterns of TBP and the TAFs (Figs. 1B and S1A). Unexpectedly, TAF2 was the only TFIID subunit, which displayed micrometer-sized, irregularly shaped nuclear foci in the DNA-free nucleoplasm (Fig. 1B, inset, white arrowheads). GFP–TAF2 protein levels in whole cell extracts were similar to those of other TFIID subunits, such as GFP–TAF5 (Fig. S1B), suggesting that protein expression levels did not primarily account for this phenomenon. The GFP–TAF2 nuclear distribution patterns appeared reminiscent of those of nuclear speckles, which are nuclear bodies, previously also referred to as interchromatin clusters (reviewed in^43–46^). These membraneless compartments concentrate mRNA splicing, processing and export factors in addition to proteins involved in transcriptional regulation^47^. Moreover, several studies suggest that nuclear speckles form via phase separation mechanisms^48–51^ and they are therefore widely considered to be biomolecular condensates^34^. To test if GFP–TAF2 nuclear foci are localized in nuclear speckles, we co-stained GFP–TAF2 cells with the nuclear speckle marker sc35, which recognizes phosphorylated forms of the nuclear speckle protein SRRM2^52^. GFP– TAF2 foci and nuclear speckles colocalized specifically (Figs. 2A and 2B, top panel and Fig. S2A), confirming that GFP–TAF2 foci indeed localize to nuclear speckles.

**Figure 2:**
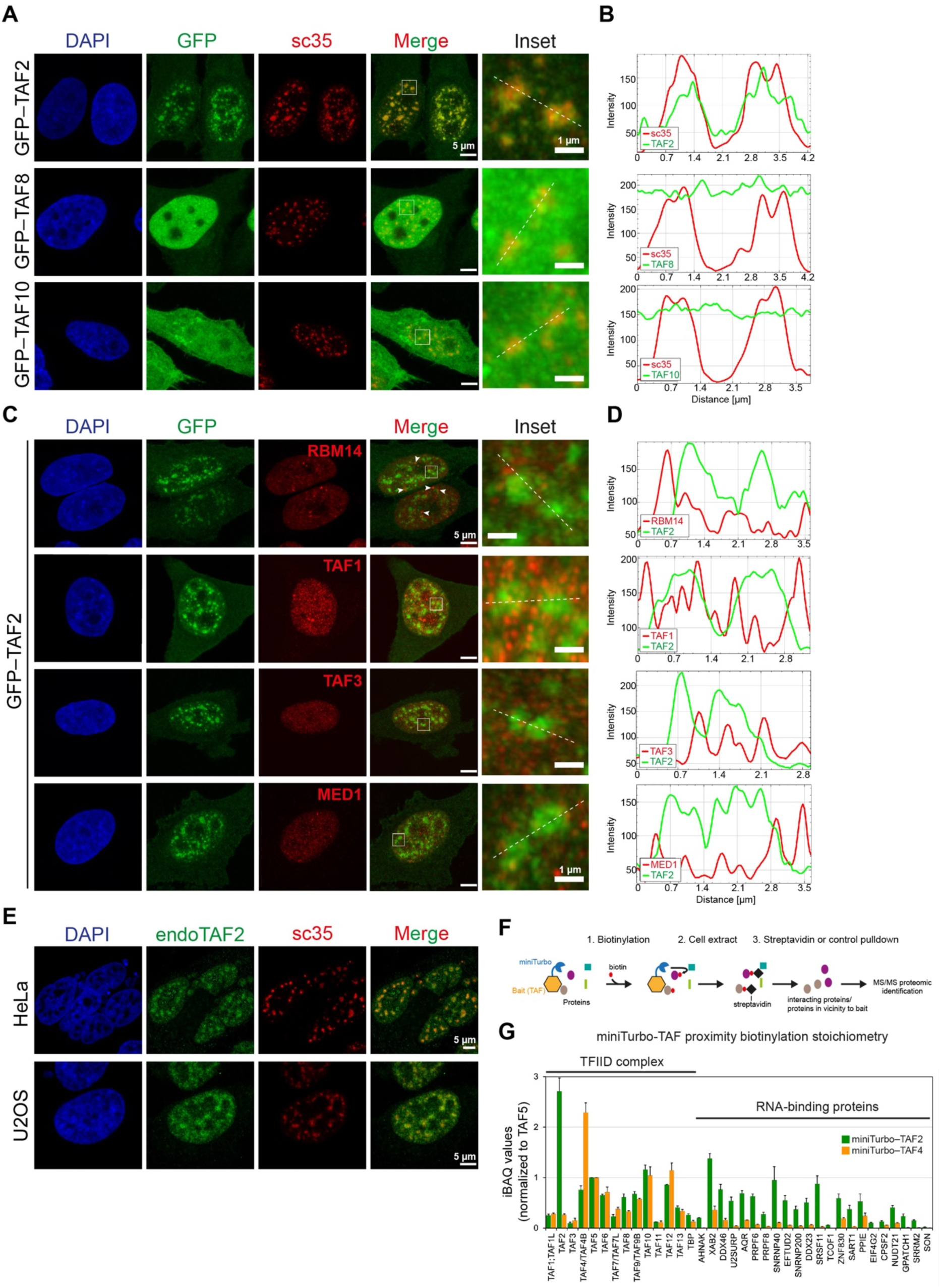
TAF2 is the only TFIID subunit that accumulates in nuclear speckles. **(A)** Fixed-cell confocal microscopy (AiryScan) images of HeLa cells expressing the individual TFIID subunits of the 3TAF complex (TAF2, TAF8, and TAF10) as GFP-fusions. Cells are stained with the nuclear speckle marker sc35. Maximum intensity projections from Z-stacks are shown. See also Fig. S2A. **(B)** Fluorescence intensity profiles along the dashed lines in the insets in **(A)**. **(C)** Fixed-cell confocal microscopy (AiryScan) images of the HeLa GFP–TAF2 cell line stained with antibodies directed against endogenous RBM14 (paraspeckle marker), TAF1, TAF3, and MED1 (Mediator subunit 1). White arrow heads point at paraspeckles lying adjacent to nuclear speckles. Maximum intensity projections from Z-stacks are shown. **(D)** Fluorescence intensity profiles along the dashed lines in the insets in **(C)**. **(E)** Fixed-cell confocal microscopy (AiryScan) images of human HeLa and U2OS cells. Single confocal planes are shown. **(F)** Schematic of the experimental strategy for proximity biotinylation using miniTurbo-tagged proteins. **(G)** Relative abundances of TFIID subunits and RNA-binding proteins from proximity biotinylation experiments. HeLa nuclear extracts expressing miniTurbo-tagged TAF2 and TAF4 were subjected to streptavidin or control pulldowns and analyzed by label-free quantitative mass spectrometry (MS). TFIID subunits and the top 20 miniTurbo–TAF2 RNA-binding protein interactors (according to *t*-test difference, Supplemental table S1), in addition to nuclear speckle proteins SRRM2 and SON are shown. Normalized iBAQ values are plotted as mean ± s.d. of technical triplicates.

### TAF2 is the only TFIID subunit, which accumulates in nuclear speckles

TAF2 forms a cytoplasmic subcomplex with TAF8 and TAF10, which is imported as so-called 3TAF complex to the nucleus^12,53^. To examine if the entire 3TAF complex could accumulate in nuclear speckles, we co-stained the GFP–TAF8 and GFP–TAF10 cell lines with sc35 and found that these TAFs did not accumulate in nuclear speckles (Figs. 2A and 2B). GFP–TAF2 neither accumulated in paraspeckles, as revealed by a co-staining with the paraspeckle marker RBM14, nor did TAF2 foci display any prominent overlaps with endogenous TAF1, TAF3 or Mediator complex subunit MED1 (Figs. 2C and D). To address if also endogenous TAF2 accumulates in nuclear speckles, we stained HeLa and human osteosarcoma cells (U2OS) with an anti-TAF2 antibody^53^. Here, also endogenous TAF2 formed nuclear foci, which co-localized with the nuclear speckle marker sc35 (Fig. 2E), suggesting that the inducible HeLa GFP–TAF2 protein expression system realistically recapitulates nuclear TAF2 protein distribution.

TAF2 has previously been implicated in transcription initiation processes merely in the context of TFIID. To validate that TAF2 is the sole TFIID subunit that accumulates in nuclear speckles, we employed an orthogonal experimental strategy using proximity labeling via the biotin ligase miniTurbo^54^. Similar to the GFP–TAF2 expression construct, we N-terminally tagged TAF2 to generate an inducible miniTurbo–TAF2 cell line. We also generated a miniTurbo–TAF4 cell line to control for TFIID-specific protein-protein interactions. We performed proximity biotinylation of TAF2 or TAF4 interacting proteins, followed by streptavidin pulldown and quantitative mass spectrometry (MS; Fig. 2F). Both miniTurbo– TAF2 and miniTurbo–TAF4 interacted with the other TFIID complex members at the expected stoichiometries^13^ (Fig. 2G, Supplemental table S1). The top TAF2 interactors (according to *t*-test difference, Supplemental table S1) included RNA-binding proteins, many of which are part of the U2 or U5 small nuclear ribonucleoprotein (U2 or U5 snRNP) splicing complexes, such as *XAB2*, *U2SURP*, *SNRNP40* or *EFTUD2* (Fig. 2G). In contrast, RNA-binding proteins were less abundant miniTurbo-TAF4 interaction partners (Fig. 2G), indicating that TAF4, as part of the TFIID complex, does not accumulate in nuclear speckles. In agreement, the gene ontology (GO) analysis of the specific interactors in both experiments revealed high enrichment scores for GO categories, such as ‘U2-type spliceosomal complex’ and ‘RNA splicing’ for TAF2 and lower scores for TAF4 (Figs. S2B and S2C). Supporting these findings, also a proteome-wide screen for RNA dependence^55^ (https://r-deep.dkfz.de) showed that TAF2 was the only TFIID subunit in the screen to display an RNA-dependent shift, indicating TAF2 interactions with RNA-binding proteins and/or RNA (Fig. S2D). In conclusion, TAF2 is the only TFIID subunit that accumulates in nuclear speckles and interacts with proteins involved in pre-mRNA splicing, such as components of the U2 snRNP complex.

### The TAF2 IDR contains several conserved molecular features including a stretch of histidines and lysines (H/K-stretch)

TAF2 contains a conserved intrinsically disordered region (IDR) at its C-terminus^29^. To understand if the TAF2 IDR contains molecular features that could drive TAF2 accumulation in nuclear speckles, we analyzed the amino-acid (aa) composition, charge distribution, and domain architecture of TAF2 (Fig. 3A). In addition to the N-terminal aminopeptidase M1-like domain, TAF2 contains a region with structural motifs identified as HEAT2-like repeats^22^ at its C-terminus. To precisely determine the boundaries of the IDR with respect to the HEAT2-like repeats, we employed two different prediction algorithms for intrinsic disorder. The PONDR-VSL2 algorithm^56^ predicted an IDR spanning aa 1009–1199 (Figs. 3A and S3A), whereas Disopred3^57^ split this IDR into two separate regions spanning aa 1005–1147 and aa 1168– 1199, respectively (Figs. 3A and S3B). The histidine/lysine stretch (H/K-stretch, aa 1142– 1171) and a region enriched in serines within Disopred IDR2 (S-stretch, aa 1168-1199) were determined using the amino-acid composition heatmap. We also searched the TAF2 sequence for nuclear localization signals (NLSs) and found a putative NLS overlapping with the positively charged H/K-stretch. Histidine repeats have previously been proposed to drive proteins to nuclear speckles^58,59^ and an evolutionary analysis revealed that this stretch is highly conserved in vertebrates and partly in insects and plants, but not in yeast (Figs. 3B and S3C), where nuclear speckles are not present^44^.

**Figure 3:**
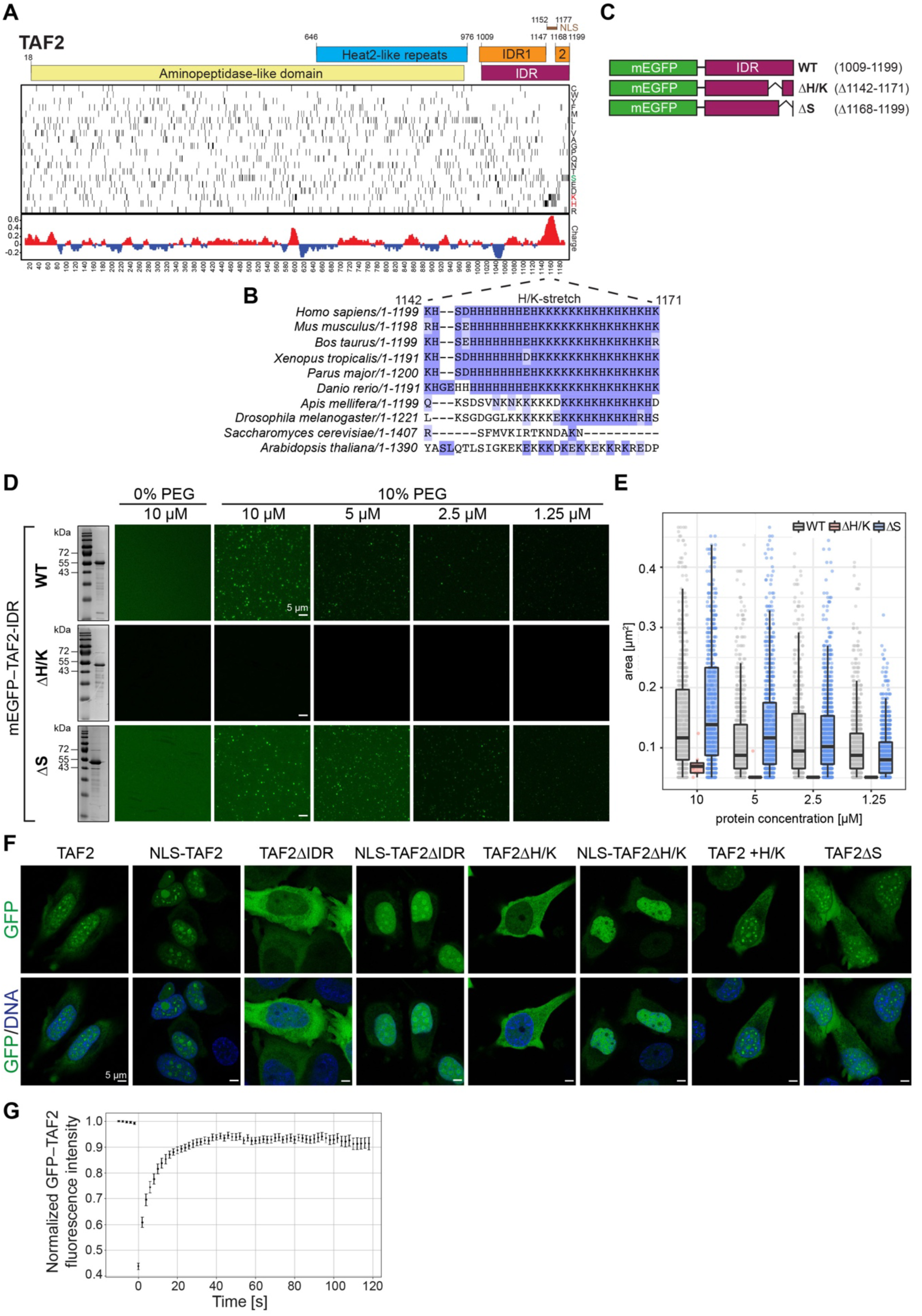
An intrinsically disordered domain (IDR) drives TAF2 condensation in nuclear speckles via a conserved histidine/lysine stretch. **(A)** Schematic of human TAF2 protein domain organization. IDRs were predicted using PONDR-VSL2^56^ and DISOPRED3^57^ (see Figs. S3A and B). The heatmap illustrates the positions of TAF2 residues (rows, amino acids; columns, position) together with the related average electrostatic charge (red, positive charge; blue, negative charge) calculated with a sliding window of 20 amino acids. **(B)** Multiple protein sequence alignment of full-length TAF2 protein sequences from 10 species across Eukarya, cropped to the H/K-stretch (human TAF2 residues 1142–1171). Residues were colored according to conservation following the Blosum62 score. See also Fig. S3C. **(C)** Schematic of TAF2-IDR-fusion constructs with monomeric EGFP (mEGFP) for bacterial protein expression and purification. **(D)** *In vitro* droplet assay in the presence or absence of polyethylene glycol 8000 (PEG-8000) with the mEGFP-TAF2 IDR fusion constructs described in **(C). (E)** Quantification of droplets in **(D)**. Four independent fields for each construct were analyzed and droplet area and number were plotted. **(F)** Live-cell confocal microscopy (AiryScan) images of HeLa cells expressing GFP–TAF2 full-length and deletion constructs under the control of a doxycycline-inducible promoter. Single confocal planes are shown. DNA was stained with Hoechst 33342. Scale bar, 5 µm. **(G)** Fluorescence recovery after photobleaching (FRAP) of GFP–TAF2 in nuclear speckles. 13 nuclear speckles from different cells were analyzed and their fluorescence normalized to background. Data are representative of two independent experiments. See also Fig. S3I. NLS, nuclear localization signal.

### The H/K-stretch conveys TAF2 IDR’s phase separation capacity and drives TAF2 to nuclear speckles

It was suggested that IDRs with charged blocks of amino acids can drive phase separation^60^. Therefore, we asked if the TAF2 IDR could form phase separated droplets *in vitro*. Bacterially expressed and purified TAF2 IDR fused to monomeric EGFP (mEGFP) via a linker sequence (Fig. 3C) formed droplets *in vitro* in a concentration-dependent manner from 1.25 µM–10 µM in the presence of 10% polyethyleneglycol 8000 (PEG-8000; Fig. 3D) and the droplets fused during time-lapse imaging (Fig. S3D). Deletion of the H/K-stretch from the TAF2 IDR diminished droplet formation, whereas deletion of the S-stretch did not decrease the droplet formation efficiency (Figs. 3D and E), suggesting that the H/K-stretch is crucial for TAF2-IDR *in vitro* phase separation. As TAF2 is not the only TFIID subunit which contains an IDR, the capacity of the other TAF IDRs to undergo *in vitro* phase separation was tested in a screen with batch-purified proteins, summarized in Supplemental table S2. Other candidates for phase separation were the IDRs of TAF1, TAF3 and TAF6, which were further tested in an *in vitro* assay. We only found the IDRs of TAF1 and TAF3 to form droplets at concentrations of 40 µM in the presence of 10% PEG (Fig. S3E), which range above the physiological concentrations typically observed for TFIID subunits in HeLa cells (estimated 2–300 nM^61^).

To test if the H/K-stretch could target TAF2 to nuclear speckles in cells, we designed GFP–TAF2 deletion constructs either lacking the entire IDR (ΔIDR), the H/K-stretch (ΔH/K) or containing only the H/K-stretch within its IDR (+H/K) (Fig. S3F). We also added a construct lacking the S-stretch (ΔS) to our set and verified similar protein expression levels in HeLa cells (Fig. S3G). Live-cell imaging of these cells showed that deleting the entire IDR or the H/K-stretch caused a strong reduction in nuclear fluorescence compared to wild-type TAF2 (Fig. 3F). As this was likely due to compromised nuclear import caused by the removal of the predicted putative NLS in the H/K-stretch, we back-added an NLS to these constructs and found that TAF2ΔIDR and ΔH/K were targeted efficiently to the nucleus (Fig. 3F). Deletion of the IDR or the H/K-stretch, but not the S-stretch, caused a diffuse nuclear TAF2 distribution pattern lacking any accumulation in nuclear speckles (Fig. 3F). Indeed, immunostainings using the nuclear speckle marker RBM25 showed that GFP–TAF2 deletion proteins colocalized with nuclear speckles only when they harbored the H/K-stretch (Fig. S3H). To understand if TAF2 is present as a liquid-like condensate in nuclear speckles, we performed fluorescence recovery after photobleaching (FRAP) experiments. Bleaching of GFP–TAF2 in nuclear speckles was followed by a time-dependent fluorescence recovery that plateaued within 20 s through influx of TAF2 from the surrounding nucleoplasm (Fig. 3G). A standard fit of the normalized curve assuming free diffusion^62^ resulted in a half maximal recovery time (*t-half*) of 4.9 s and the mobile fraction was estimated to 87% (Fig. S3I). These results show that TAF2 molecules in nuclear speckles exchange rapidly with the nucleoplasm.

In summary, these data suggest that the H/K-stretch within the TAF2 IDR represents a putative nuclear speckle targeting signal, driving TAF2 condensation in nuclear speckles. TAF2 forms a liquid-like phase separated compartment, with a molecules exchange on the timescale of seconds with the surrounding nucleoplasm. Further, the TAF2 IDR was the only tested IDR in the TFIID complex capable of *in vitro* droplet formation at concentrations in the lower micromolar range, highlighting robust TAF2-IDR phase separation properties.

### The TAF2 IDR mediates interactions with the spliceosome-associated protein SRRM2 and is not required for TFIID complex assembly

To examine the functional relevance of the TAF2 IDR within the TFIID complex, we performed GFP pulldowns from nuclear extracts of HeLa cells expressing GFP–TAF2, GFP-TAF2ΔIDR, or GFP-NLS-TAF2ΔIDR (Fig. S4A) and subjected them to label-free quantitative MS (Fig. 4A; Supplemental table S3). Wild-type TAF2 as well as both IDR-deletion proteins associated with the other TFIID complex members at similar stoichiometries (Fig. 4B, Supplemental table S3), suggesting that the TAF2 IDR is not required for TAF2 incorporation into the TFIID complex and general TFIID assembly. Upon IDR deletion, a total of 27 out of 39 significant interactors were retained, including the RNA-binding protein RBM14 and the TFIID subunits (Figs. 4A–B, S4B, Supplemental table S4). IDR deletion resulted in an increase of significant interactors, including many RNA-binding proteins in both TAF2ΔIDR (increase by 79) and NLS-TAF2ΔIDR (increase by 31) experiments (Supplemental table S3), indicating that these interactions are favored by the lack of the TAF2 IDR. As both TAFΔIDR constructs do not accumulate in nuclear speckles (Fig. 3F), this phenomenon may be due to increased interaction frequencies between the TAF2ΔIDR constructs and freely diffusing RNA-binding proteins in the nucleoplasm. Importantly, we identified only three proteins whose interactions with TAF2 were lost upon IDR deletion, namely the ribosomal subunit proteins RPS23 and RPS3A, and the nuclear speckle and spliceosome-associated protein serine/arginine repetitive matrix protein 2 (SRRM2, Figs. 4A–B, S4B, Supplemental table S4). An SRRM2 stoichiometry of approximately 0.2 in the wild-type GFP–TAF2 pulldown suggested that substoichiometric SRRM2 protein levels are associated with TFIID (Fig. 4B). As SRRM2 was the only nuclear speckle protein affected, we verified the SRRM2-TAF2 association in co-immunoprecipitations (co-IPs) suitable for the isolation and detection of nuclear speckle proteins^52^. In concordance with the MS results, GFP–TAF2 interacted with SRRM2, whereas the GFP-NLS-TAF2ΔIDR and GFP–TAF5 constructs did not display any detectable SRRM2 interactions (Fig. 4C). Since TAF2 and SRRM2 both accumulate in nuclear speckles and the co-IPs detected specific interactions between these proteins, we concluded that TAF2 and SRRM2 likely interact with each other in nuclear speckles.

**Figure 4:**
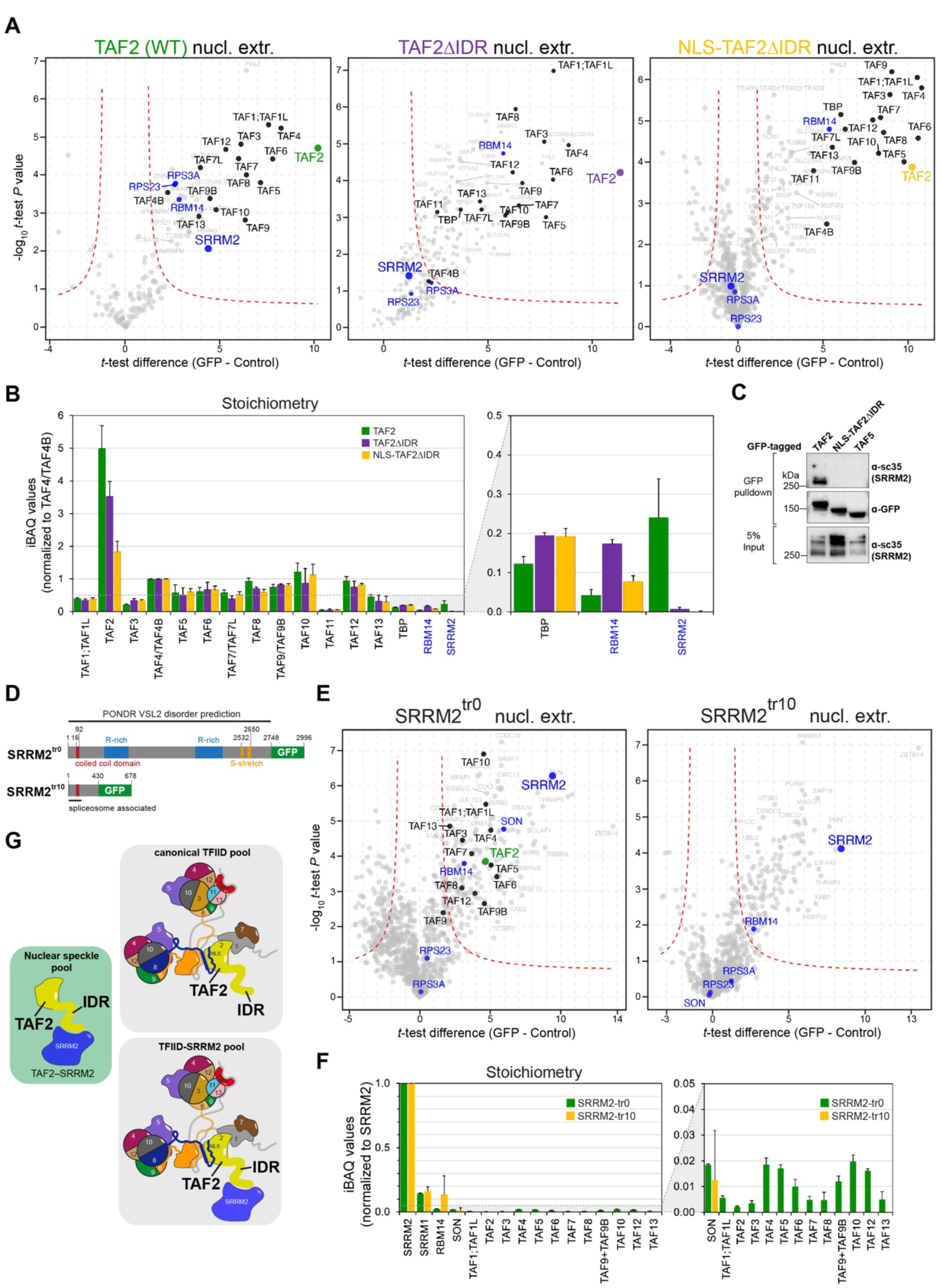
The TAF2 IDR recruits SRRM2 to the TFIID complex. **(A)** Volcano plots displaying label-free quantitative mass spectrometry (MS) data. GFP or control pulldowns were performed as technical triplicates from HeLa cell nuclear extracts expressing GFP-tagged wild-type TAF2 (WT, green), TAF2ΔIDR (purple) and NLS-TAF2ΔIDR (yellow). TFIID subunits are colored black, the nuclear speckle proteins SRRM2 and the RNA-binding proteins RBM14, RPS23, and RPS3A are colored blue. Dashed red lines indicate the threshold between non-significant and significant interactors (two-sample *t*-test; FDR=1%; *S*0=1). Not all TFIID subunits are displayed due to MS data filtering criteria (see methods section). See also Fig. S4A. **(B)** Relative abundances (stoichiometries) of TFIID subunits in addition to RBM14 and SRRM2 from the quantitative MS experiments in **(A)**. Normalized iBAQ values are plotted as mean ± s.d. of technical triplicates. **(C)** Co-immunoprecipitation (co-IP) assays from HeLa cell extracts were run on a 4–12% bis-tris polyacrylamide gel followed by immunoblot. The interactions with SRRM2 were probed using the sc35 antibody, which recognizes phosphorylated SRRM2^52^. Uncropped images with several exposure times are shown in the source data related to Fig. 4C. Data are representative of n=2 experiments. **(D)** Schematic displaying SRRM2 proteins expressed in two engineered HAP1 cell lines^52^. Endogenous SRRM2–GFP fusion proteins are expressed as full-length (SRRM2^tr0^) or IDR-deletion proteins (SRRM2^tr10^). Illustrated are the positions of the coiled coil domain, arginine-rich (R-rich), serine-rich (S-rich) domains and the IDR predicted by the PONDR-VSL2^56^ algorithm. The conserved first 135 aa of the yeast homolog Cwc21 are associated with the spliceosome^65^. **(E)** Volcano plots displaying label-free quantitative MS data. GFP or control pulldowns were performed as technical triplicates from HAP1 nuclear extracts expressing endogenous SRRM2^tr0^ or SRRM2^tr10^. Threshold between non-significant and significant interactors: two-sample *t*-test; FDR=0.1%; *S*0=1 for SRRM2^tr0^ and FDR=1%; *S*0=1 for SRRM2^tr10^. The SRRM2^tr0^ GFP-pulldown and MS data are representative of two biological replicates. **(F)** Stoichiometries of TFIID subunits and RNA-binding proteins SRRM1, RBM14, and SON from quantitative MS experiments in **(E)**. Normalized iBAQ values are plotted as mean ± s.d. of technical triplicates. **(G)** Schematic illustrating different nuclear TAF2 compartments. Half circles in TFIID represent histone-fold domain (HFD) partners; the lines represent protein domains not resolved by cryo-EM. SRRM2 is depicted as globular structure for visual clarity.

Together, these findings demonstrate that the TAF2 IDR is neither required for TAF2 incorporation into TFIID, nor for TFIID complex assembly. Instead, the TAF2 IDR is required for the interaction with the spliceosome-associated protein SRRM2.

### The splicing factor SRRM2 interacts with the TFIID complex via IDR-IDR interactions

Human SRRM2 (also known as SRm300) exhibits multiple stretches of serines and arginines (Fig. S4C), localizes to nuclear speckles^63,64^, and upholds the nuclear speckle structure together with the RNA-binding protein SON^52^. SRRM2 is a splicing factor involved in pre-mRNA splicing as part of the spliceosome B^act^ stage^65^, but it is not required for constitutive pre-mRNA splicing^66^. SRRM2 specifically interacts with the U2 snRNP complex^67^, whose components were also significantly enriched in miniTurbo–TAF2 proximity labeling MS data (Fig. 2D). To further dissect the physical interaction between SRRM2 and TAF2, we used engineered human HAP1 cell lines^52^, which endogenously express the SRRM2 full-length protein (SRRM2-tr0) or a truncated form of SRRM2 (SRRM2-tr10) as GFP fusions (Fig. 4D). SRRM2 is predicted to be intrinsically disordered along the entire protein length^52^, however, SRRM2-tr10 retains the domain, which associates with the spliceosome and can thus be considered an SRRM2-IDR deletion protein. In both cell lines nuclear speckle integrity remained intact (Fig. S4D) as described previously^52^. As both cell lines express SRRM2 as C-terminal GFP fusions, we employed the quantitative MS assay with GFP pulldowns and conditions optimized to detect TFIID interactions (Supplemental table S5). We found that full-length SRRM2 (SRRM2-tr0) interacted with the expected partners, such as SON and, remarkably, also with almost all TFIID subunits including TAF2 (Figs. 4E and F). Upon SRRM2-IDR deletion (SRRM2-tr10), the interactions with SON and the TFIID complex were compromised, in fact TFIID peptides were not detected in this experiment (Supplemental table S5). Importantly, TAF2 and TAF1 proteins could still be detected in the nuclear fractions of these cells (Fig. S4E), underlining that SRRM2 interactions with these TFIID subunits were indeed compromised and not due to reduced nuclear TAF1 or TAF2 protein levels. These experiments validate the MS data of GFP–TAF2 pulldowns (Figs. 4A and B) and show that SRRM2 and TFIID can form a complex via interactions between the TAF2 and SRRM2 IDRs. Given that the SRRM2-tr0-GFP pulldown revealed the entire TFIID complex, we concluded that we isolated a non-canonical TFIID-SRRM2 pool in the nucleoplasm and not the TAF2– SRRM2 pool residing in nuclear speckles.

These data, in combination with previous biochemical^28,52^, cryo-EM^10,11^, and imaging data presented here, indicate that nuclear TAF2 can exist in three protein pools: in the canonical holo-TFIID complex, in a non-canonical TFIID complex associated with SRRM2, and in a complex with SRRM2 in nuclear speckles (Fig. 4G).

### The TAF2 IDR is important for appropriate TAF2 promoter association

To study the importance of the TAF2 IDR in the context of TFIID, we analyzed previously described TAF2 genomic functions. TAF2 binds non-sequence specific promoter DNA as shown by electromobility shift assays^18^, DNA footprinting assays^29^, and cryo-EM studies^11,14^. Interestingly, the DNA footprinting study has shown that a tin-based oxochloride compound could inhibit the IDR-DNA interaction and thereby enhance TFIID promoter binding^29^. To understand if the TAF2 IDR plays a role in promoter binding on the genome-wide scale and in living cells, we used a modified protocol for cleavage under targets and release using nuclease (CUT&RUN) called greenCUT&RUN^68,69^, which specifically localizes GFP-tagged proteins on chromatin. We found that GFP-tagged TAF2 and NLS-TAF2 localized to transcription start sites (TSSs), as expected (Fig. S5A). As a subset of the TAF2 protein pool localizes to nuclear speckles, which are non-genomic sites, we expected an increase in TAF2 promoter occupancy upon disruption of the TAF2-nuclear speckle association. Indeed, the TAF2 constructs deficient for nuclear speckle association (GFP-NLS-TAF2ΔIDR and GFP-NLS-TAF2ΔH/K) exhibited a strong increase in coverage over TSSs compared to their wild-type counter parts (Figs. S5A and B). This effect was more pronounced when coverage was plotted over TAF1 peak centers (Fig. 5A) or when it was visualized in snapshots of genomic tracks over promoters typically bound by TFIID (Fig. 5B).

**Figure 5:**
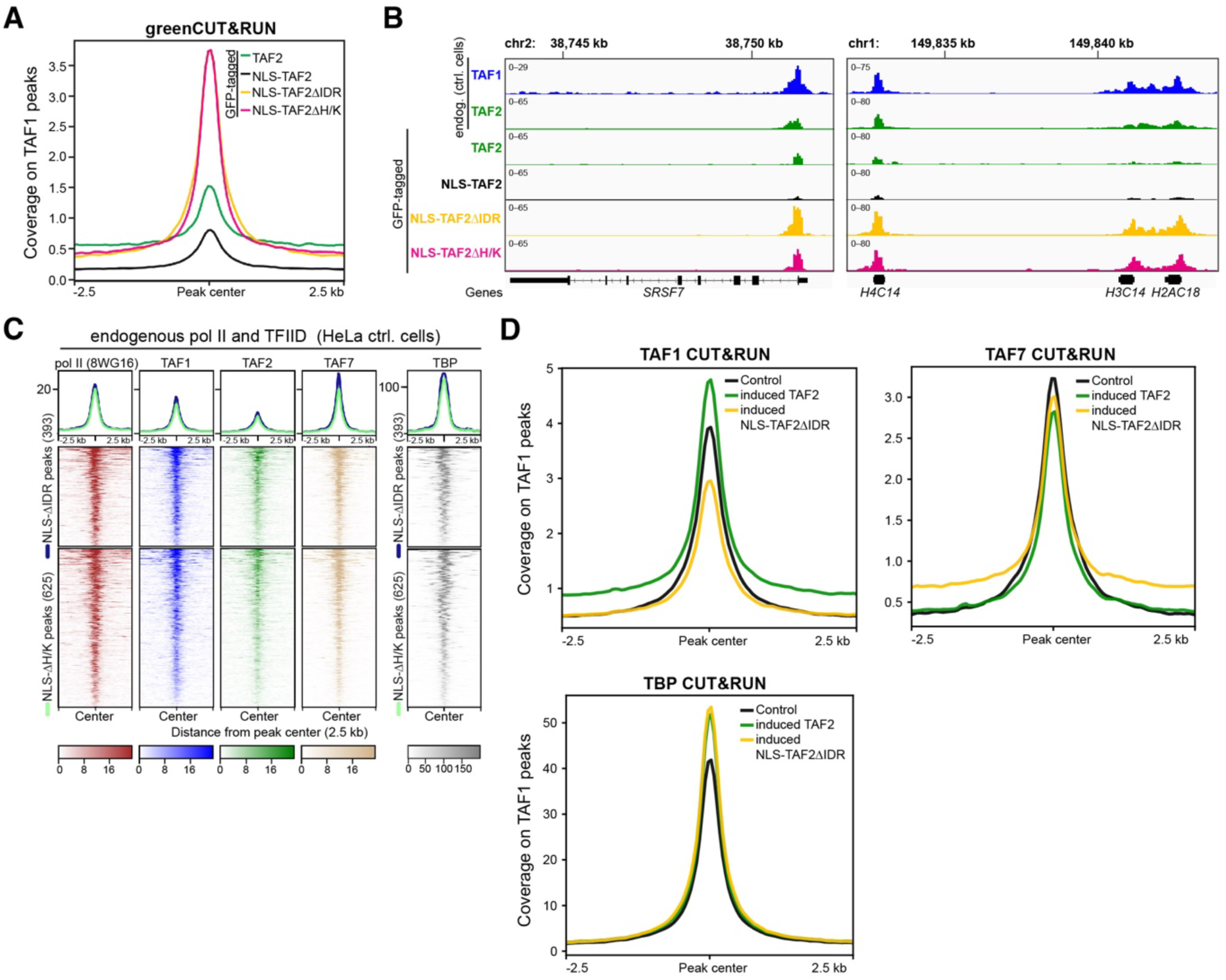
The TAF2 IDR is important for appropriate TAF2 promoter association. **(A)** Profile plot displaying greenCUT&RUN coverage of GFP-tagged full-length TAF2 or deletion constructs expressed in HeLa cells. Spike-in normalized coverage (1x normalization, RPGC) is centered on TAF1 peak regions (9574) and the coverage 2.5 kilobases (kb) up- and downstream is shown. See also Figs. S5A and B. **(B)** IGV tracks illustrating coverage of greenCUT&RUN experiments shown in **(A)**, in addition to endogenous TAF1 and TAF2 in HeLa control (ctrl.) cells. Peaks on the promoters of the genes *SRSF7* and the histone cluster genes *H4C14*, *H3C14* and *H2AC18* are displayed. **(C)** Heatmaps displaying CUT&RUN coverage of RNA polymerase II (pol II) and endogenous TFIID subunits in control (ctrl.) HeLa cells. The spike-in normalized coverage is centered over GFP-NLS-TAF2ΔIDR (393) and GFP-NLS-TAF2ΔH/K (625) peaks and coverage 2.5 kb up- and downstream is shown. **(D)** Profile plots of CUT&RUN coverage of endogenous TAF1, TAF7 and TBP in HeLa control and GFP–TAF2 or GFP-NLS-TAF2ΔIDR expressing cells. Spike-in normalized coverage is centered on TAF1 peak regions and the coverage 2.5 kilobases (kb) up- and downstream is shown. Data are representative of two biological replicates. NLS, nuclear localization signal.

To confirm that the diffuse nuclear TAF2 constructs localize predominantly to pol II promoters, we identified 625 and 393 common peaks from two independent greenCUT&RUN experiments for the GFP-NLS-TAF2ΔH/K and GFP-NLS-TAF2ΔIDR constructs, respectively. Most of these peaks occurred in promoter regions as did TAF1 or TAF2 peaks in HeLa control cells (Fig. S5C). Further, spike-in normalized coverages of RNA pol II, TAF1, TAF2, TAF7, and TBP from CUT&RUN in control HeLa cells enriched sharply over the GFP-NLS-TAF2ΔIDR and GFP-NLS-TAF2ΔH/K peak centers (Fig. 5C), demonstrating that diffuse nuclear TAF2 occupies *bona fide* TFIID binding sites on pol II promoters. We therefore asked if promoter occupancy of other TFIID subunits was changed in the presence of enhanced GFP-NLS-TAF2ΔIDR promoter occupancy. CUT&RUN for endogenous TAF1, TAF7, and TBP in the cell lines expressing GFP–TAF2 or GFP-NLS-TAF2ΔIDR showed that enhanced GFP-NLS-TAF2ΔIDR promoter enrichment did not cause enhanced TAF1 or TAF7 promoter association (Fig. 5D). Whereas TAF7 promoter enrichment remained unchanged, TAF1 enrichment was slightly reduced in the GFP-NLS-TAF2ΔIDR expressing cells. In contrast, TBP promoter association was increased in cells expressing either GFP–TAF2 or GFP-NLS-TAF2ΔIDR (Fig. 5D), in line with previous results from DNA footprinting assays^29^. These data suggest that increased TAF2 enrichment on promoters may interfere with TAF1-promoter binding. It is possible that the increased GFP-NLS-TAF2ΔIDR enrichment is caused by TAF2 molecules not properly evicted from promoters during transcription initiation, potentially leading to sterical interference with TAF1 which is positioned close to TAF2 in lobe C^11,70^.

In conclusion, the TAF2 IDR is important for appropriate TAF2-promoter DNA association, which appears to involve TAF2 promoter release for subsequent steps during transcription initiation. Further, TAF2 molecules lacking their IDR-encoded nuclear speckle targeting signal relocate to RNA pol II promoters, indicating shuttling between nuclear speckles and TAF2-bound promoters.

### The TAF2 IDR has a minor impact on global gene expression

To analyze the impact of the TAF2-IDR deletion construct on gene expression, we performed RNA sequencing (RNA-seq) of poly(A)-containing mRNAs and compared global transcript profiles of cells expressing full-length GFP–TAF2 or GFP-NLS-TAF2ΔIDR. The MS data (Fig. 4) showed that the expression of these two TAF2 constructs induced the formation of SRRM2-containing and SRRM2-lacking TFIID complexes (Fig. 6A). Differential expression (DE) analysis between TAF2 expressing and control cell lines revealed 1147 and 1517 (TAF2) and 1075 and 1593 (NLS-TAF2ΔIDR) significantly (*P*-valadj<0.05) up- and downregulated genes, respectively (Figs. S6A–B and Supplemental table S6). Common significantly up-regulated genes included the serine-peptidase inhibitor *SPINK6*, the proto-oncogene *MYC*, and the transmembrane receptor *MUC13* (Fig. 6B). Commonly downregulated genes were related to extracellular matrix organization, such as the collagen chains *COL1A1 or COL1A2* (Fig. 6B). We only found a small set of uniquely up- or downregulated genes in TAF2 or NLS-TAF2ΔIDR expressing cells, such as the transcription factor *IRX4*, which was significantly downregulated only in the NLS-TAF2ΔIDR condition and the phosphatases *DUSP1* and *ALPI* that were significantly upregulated only in the wild-type TAF2 condition (Fig. 6B). Further, the increased promoter enrichment of GFP-NLS-TAF2ΔIDR did not coincide with significant changes in gene expression, except for the up-regulated genes *MYC* and *IER5L* with log2-fold changes above 0.4 (Fig. S6C). Thus, the gene expression changes in both TAF2 and NLS-TAF2ΔIDR expressing cells compared to controls were highly similar and GO analyses did not reveal any specific changes in categories enriched in the NLS-TAF2ΔIDR condition (Fig. S6D).

**Figure 6:**
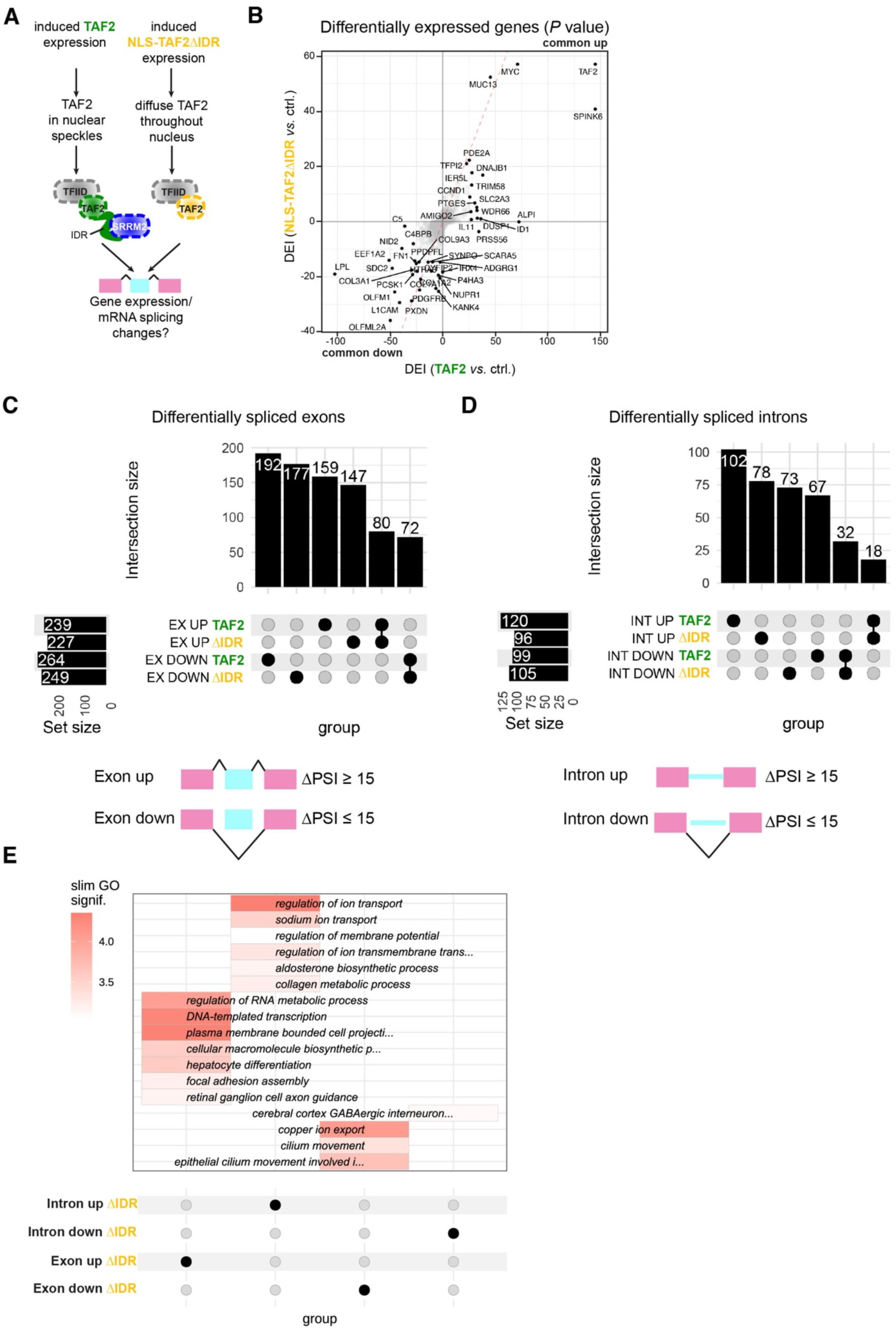
SRRM2-TFIID complexes have minor effects on global gene expression but elicit unique alternative splicing events. **(A)** Schematic illustrating canonical (left) and non-canonical (right) TFIID complexes upon the induced expression of GFP–TAF2 or GFP–NLS-TAF2ΔIDR constructs. **(B)** RNA differential expression (DE) analysis. The plot shows the Differential Expression Index (DEI) of the comparisons TAF2 *vs.* control (ctrl.) and NLS-TAF2ΔIDR *vs.* control (ctrl.) on the y and x-axis, respectively. DEI is defined as the product of the sign of the log2 fold change (FC) and the -log10 of its adjusted *P* value: DEI*=*sign[log(FC)]⋅[-log10(*P*-valadj)]^111^. Common up- and downregulated genes can be found in the top right and bottom left corners of the plot, respectively. The top 30 genes with the lowest adjusted *P* values of the DE analysis were highlighted for each comparison. Genes lying on the diagonal (dashed red line) display equal significant DEI in both comparisons. Each comparison was performed on technical triplicates. The Volcano plots for the individual comparisons can be found in Figs. S6A and B. **(C)** Upset plot showing the intersections of common and unique alternative splicing events of exons **(C)** and introns **(D)**. The threshold for an event to be considered was percentage of sequence inclusion |ΔPSI| ≥15. EX, exons; INT, introns. **(E)** Gene ontology (GO) analysis of unique alternative splicing events in GFP-NLS-TAF2ΔIDR expressing cells, performed using GOfuncR^115^ and slimGO. SlimGO categories are clustered by mean (slim) GO significance in each gene group. See also Fig. S6E.

In conclusion, the expression of both TAF2 proteins, GFP–TAF2 and GFP-NLS-TAF2ΔIDR, elicit similar global gene expression changes at the level of mature mRNAs. Therefore, neither SRRM2 recruitment to TFIID nor increased TAF2 promoter occupancy appear to have a major impact on global gene expression.

### SRRM2-TFIID complexes elicit a set of unique alternative splicing events

As the TAF2 IDR drives TAF2 to nuclear speckles and recruits the splicing factor SRRM2 to the TFIID complex, we tested if the induced expression of GFP-NLS-TAF2ΔIDR affects pre-mRNA splicing (Fig. 6A). We analyzed alternative splicing (AS) using the poly(A)-containing mRNA data set and evaluated annotated splice junctions based on reads spanning exons (referred to as ‘alternative splicing events’). Interestingly, we detected differential AS of exons and introns in GFP–TAF2 or GFP-NLS-TAF2ΔIDR expressing cell lines relative to controls, with an absolute difference in percentage of sequence inclusion (ΔPSI) greater than or equal to 15 (Supplemental table S7). In total, we identified in GFP–TAF2 and GFP-NLS-TAF2ΔIDR expressing cells 80 and 72 common cassette exon inclusion and skipping events, respectively (Fig. 6C). Moreover, 32 common intron splicing and 18 common intron retention events were identified between the two conditions (Fig. 6D). However, we identified a larger set of events that were unique to either GFP–TAF2 or GFP-NLS-TAF2ΔIDR expression (Figs. 6C and D). Accordingly, we found a total of 351 uniquely differentially spliced exons in 306 genes in the wild-type TAF2 condition and 324 uniquely differentially spliced exons in 291 genes in the NLS-TAF2ΔIDR condition (Fig. 6C, Supplemental table S7). Regarding alternatively spliced introns, we detected 169 unique events in 146 genes in the wild-type condition and 151 unique events in 138 genes in the NLS-TAF2ΔIDR condition (Fig. 6D, Supplemental table S7). Both, the induced expression of exogenous TAF2 and the IDR-deletion mutant caused alternative splicing events, suggesting that modulation of TAF2 protein levels *per se* influences RNA splicing. To further explore the possible biological consequences of the TFIID-SRRM2 interaction, we examined the enrichment of GO terms in the genes from each group of uniquely changed events in the wild-type TAF2 and ΔIDR conditions (Fig. S6E). In the ΔIDR condition, overrepresented pathways in genes from exon inclusion events (‘exon up’) included ‘transcription’ and ‘RNA metabolic process’, whereas the GO category ‘copper ion transport’ was enriched in genes from exon skipping events (‘exon down’, Figs. 6E and S6E). Changed retained introns in the ΔIDR condition occurred in pre-mRNAs of genes related to transmembrane transport (‘introns up’) and increasingly spliced out introns occurred in pre-mRNAs of genes related to neurodevelopmental processes (‘introns down’, Figs. 6E and S6E). These changes suggest that reduced SRRM2 recruitment to TFIID influences specific pre-mRNA splicing processes.

In conclusion, our data suggest that the presence or absence of SRRM2 in TFIID complexes, mediated by the TAF2 IDR, elicits a set of unique alternative splicing events. The formation of SRRM2-free TFIID complexes might lead to the production of different protein isoforms that contribute to the regulation of biological processes including transcription and transmembrane transport.

## Discussion

Here we report that the TFIID subunit TAF2 is sequestered in nuclear speckles and recruits the splicing factor SRRM2 to TFIID. Our findings can be summarized in a model for TAF2 nuclear compartmentalization, in which TAF2 is partitioned into several protein pools (Fig. 7). Here, nuclear TAF2 could dissociate from canonical holo-TFIID and translocate to nuclear speckles, a process driven by the TAF2 IDR (Fig. 3). Our findings are consistent with previous data, which demonstrated that nuclear TAF2 associates substoichiometrically with the TFIID complex in human^12,13,19,28^ and yeast^71^. IDRs can act as molecular switches to partition proteins into different types of condensates (reviewed in^30^). For example, serine-2 phosphorylation of the RNA pol II C-terminal domain, a well-known IDR^72,73^, drives the exchange from RNA pol II initiation to elongation condensates^74^. The TAF2 IDR is also extensively phosphorylated^75^ and modulation of its phosphorylation status could provide a signal to switch between nuclear compartments. We propose that TAF2 localizes to at least two compartments, nucleoplasmic holo-TFIID and nuclear speckle condensates. After dissociation from holo-TFIID, TAF2 could interact with SRRM2 in nuclear speckles through IDR-IDR contacts, leave nuclear speckles as TAF2–SRRM2 complex, and re-associate with the TFIID complex, which is supported by our MS data (Fig. 4). The resulting non-canonical TFIID-SRRM2 complex could bind, like canonical TFIID, to pol II promoters and form pre-initiation complexes (PICs) to affect alternative splicing of pre-mRNAs.

**Figure 7:**
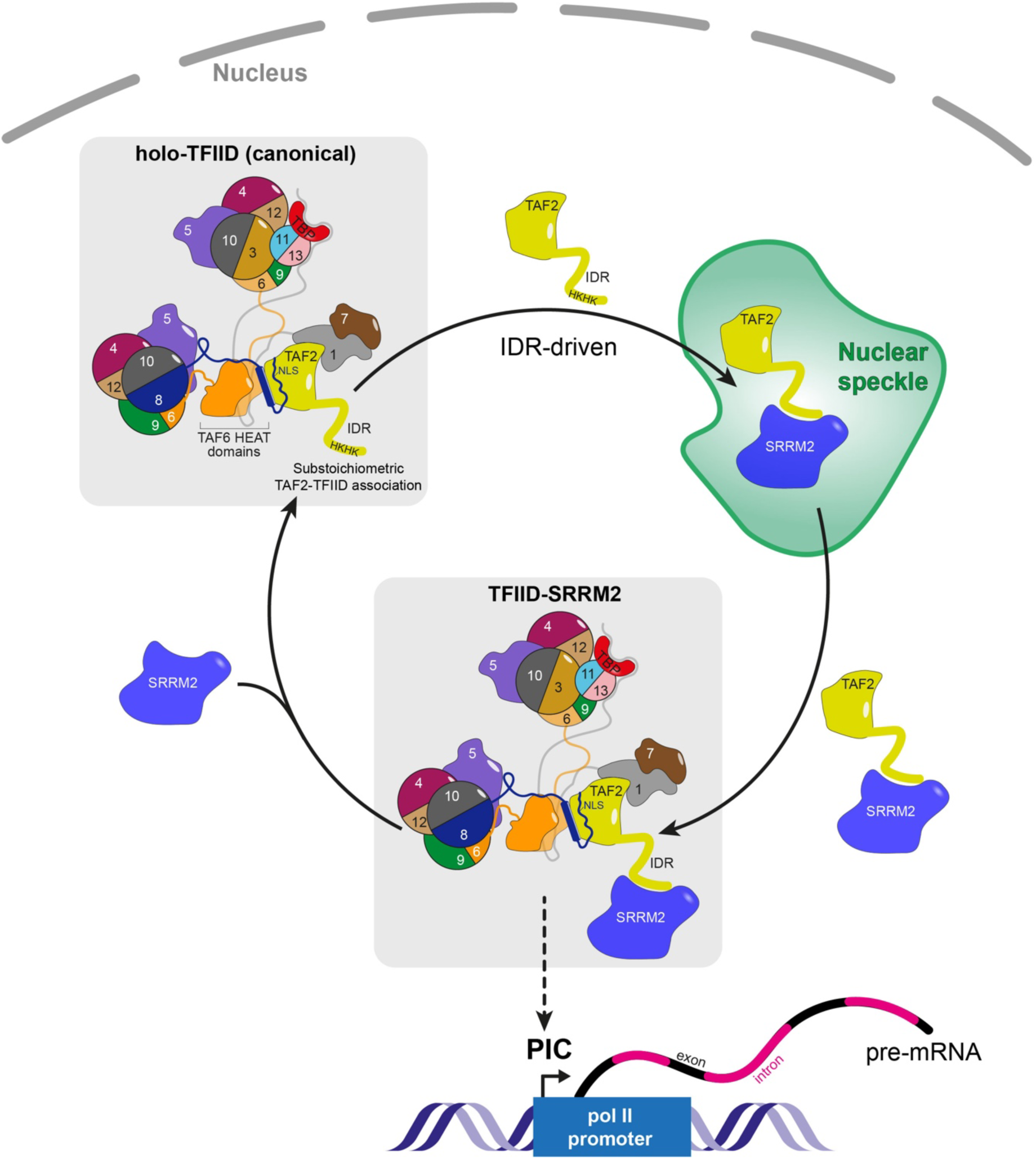
Model for nuclear TAF2 compartmentalization driven by the TAF2 IDR. Scheme of the proposed TAF2 nuclear compartmentalization cycle. TAF2 is compartmentalized into a canonical holo-TFIID pool and into nuclear speckles. TAF2 is substoichiometrically associated with holo-TFIID and targeted to nuclear speckles via the histidine/lysine (H/K)-stretch within the TAF2 IDR. In nuclear speckles, TAF2 associates with SRRM2 via IDR-IDR interactions and then reassociates with TFIID to form a non-canonical TFIID-SRRM2 complex. It is possible that the TFIID-SRRM2 complex, like canonical holo-TFIID, is recruited to promoters to form pre-initiation complexes (PICs) where they could influence splicing of nascent pre-mRNA. IDR, intrinsically disordered region; NLS, nuclear localization signal; pol II, RNA polymerase II. For further details, see the discussion section.

We have demonstrated that the TAF2 IDR contains a putative nuclear speckle targeting signal, consisting of a conserved stretch of histidines and lysines (H/K-stretch, Fig. 3F). Although histidine stretches have been shown to target proteins to nuclear speckles^58,59^, it is unclear if these sequences bind IDR-specific ‘driver’ proteins that help them find their designated nuclear compartment. The H/K-stretch differs from a recently identified 15-amino acid putative nuclear speckle targeting motif (STM) characterized by periodicity of proline residues^76^. This motif occurs in speckle-targeting transcription factors, such as p53, required for chromatin-nuclear speckle association. p53 was shown to associate with nuclear speckles for amplified RNA expression^77^, but the protein does not accumulate in nuclear speckles like TAF2. Thus, it is plausible that speckle-targeting and -accumulating proteins might utilize different targeting motifs, respectively. Proteins with arginine-enriched mixed charge domains have been shown to undergo condensation and nuclear speckle incorporation^78^. The TAF2 IDR does not contain such molecular features, but a serine-rich stretch at the C-terminus, which, when phosphorylated, could provide a negative charge domain. It is also possible that the TAF2 IDR associates directly with nuclear speckle RNAs, such as the long non-coding RNA (lncRNA) MALAT1, but a recent screen for RNA-interacting transcription factors has not identified TAF2 as a hit^79^. Instead, we showed that the TAF2 IDR binds specifically to SRRM2 (Figs. 4A–C), which could delegate TAF2 to nuclear speckles. The H/K-stretch could be a general characteristic of nuclear speckle proteins involved in transcriptional regulation. Another example of a histidine-repeat containing nuclear speckle protein is the kinase cyclinT1 (*CCNT1*), which is part of Positive transcription elongation factor b (P-TEFb)^58,80^ and forms liquid-like droplets in a histidine-repeat dependent manner^81^. In our assays, deletion of the H/K-stretch also compromised TAF2 nuclear import, suggesting alternative TAF2 import mechanisms in addition to the reported TAF2 import via the TAF8 NLS^53^. Indeed, it has been shown that nuclear TAF2 can be detected by mass spectrometry in the absence of TAF8^82^.

The TAF2 mass spectrometry data and reciprocal pulldown experiments using SRRM2 as a bait revealed the existence of non-canonical TFIID complexes, which are associated with substoichiometric levels of SRRM2 (Fig. 4). Non-canonical TFIID complexes have been reported in human cells, such as TFIID complexes lacking TAF10^83^ or TAF7^84^. TFIID has also been shown to be recruited specifically to promoter-proximal 5’ splice sites^85^ and to associate with the cleavage–polyadenylation specificity factor (CPSF)^86^. Together, these studies support our model, in which TAF2 dissociates from TFIID to promote the formation of non-canonical TFIID complexes with functions in RNA processing. Indeed, our proximity labeling experiments with TAF2 and TAF4 (Fig. 2D) revealed several subunits of the CPSF complex as significant interactors in both TAF2 and TAF4 experiments, underlining that TFIID is associated with factors of the RNA processing machinery. TAF2 also interacts with many proteins of the U2 snRNP complex and these interactions might be mediated not only by the IDR, but also by the structured TAF2 protein domains. The TAF2 IDR, however, is specifically responsible for the interaction with SRRM2. We did not address if SRRM2 can be recruited to the PIC via TFIID, but it is an intriguing possibility that TAF2-SRRM2 complexes serve to bring SRRM2 out of nuclear speckles to nascent pre-mRNAs where it can join the active spliceosome.

The alternative splicing (AS) analysis revealed hundreds of unique alternative splicing events in cells forming TFIID complexes associating with or lacking SRRM2. These results are in line with reports showing that SRRM2 is not required for constitutive splicing^66^ and that it regulates splicing of only a subset of RNAs coding for proteins involved in innate immunity, cell homeostasis^64,66^ or inflammatory metabolism^87^. Further, and expectedly, the observed changes in AS upon reduced TFIID-SRRM2 complex formation were less pronounced compared to a *SRRM2* gene knockdown^87^ since TFIID-associated SRRM2 only represents a small fraction of total nuclear SRRM2. However, observed AS changes were found in RNAs encoding proteins related to transmembrane transport and neurodevelopmental processes, also affected by SRRM2 dysfunction in humans. For instance, an *SRRM2* loss of function variant was shown to cause a rare neurodevelopmental disorder^88^ and SRRM2 is involved in the etiology of Alzheimer’s^89^ and Parkinson’s disease^90^. Our study thus provides evidence for the function of an IDR within the large multiprotein complex TFIID that controls nuclear compartmentalization and thus links distinct molecular processes, such as transcription initiation and RNA splicing.

### Limitations of the study

The lack of a reliable commercially available TAF2 antibody has restricted our study of the functions of the endogenous TAF2 protein and the limiting amounts of the polyclonal TAF2 antibody (pAb 3038^53^) were crucial to show that also endogenous TAF2 accumulates in nuclear speckles. Further, we have not created TAF2 knockout cell lines, as TAF2 is an essential gene (https://depmap.org/portal/gene/TAF2?tab=overview). Therefore, our study relies on inducible TAF2 or TAF2ΔIDR expression in the presence of endogenous TAF2, which leads to the formation of a mixture of TFIID complexes. Future studies will benefit from the creation of cell lines, in which endogenous TAF2 can be inducibly depleted.

## Materials and methods

### Generation of bacterial and mammalian expression plasmids

For mammalian protein expression, cDNA of human TAF2 was amplified by PCR with gene-specific primers fused to attB recombination sequences for Gateway® cloning (Thermo Fisher Scientific). The amplified sequence was recombined via BP reaction into the pDONR201 donor resulting in pENTR-TAF2 for N-terminal tagging. The TAF2 sequence was recombined via LR reaction into a pcDNA5/FRT/TO plasmid containing GFP^91^ or miniTurbo N-terminal of the recombination site. The miniTurbo cDNA was PCR amplified from the 3xHA-miniTurbo-NLS_pCDNA3 plasmid, a gift from Alice Ting (Addgene plasmid #107172; http://n2t.net/addgene:107172; RRID:Addgene_107172). For the GFP–TAF1 construct, a pcDNA5/FRT/TO plasmid containing the β-globin intron C-terminal of GFP was used to enhance GFP–TAF1 protein expression. Gateway cloning was performed according to the manufacturer’s protocol (Thermo Fisher Scientific). All other TAFs were cloned into expression vectors in the same manner, except for TAF4, whose cDNA was assembled from three fragments using NEBuilder® HiFi DNA Assembly Master Mix (NEB, E2621S). A TAF4 G/C-rich fragment was synthesized as gene block with reduced G/C content (IDT).

For bacterial protein expression, PCR-generated TAF cDNA encompassing the defined intrinsically disordered domains (IDRs) was inserted into pET-mEGFP expression plasmid^35^ (a kind gift by Richard Young, MIT, Boston, USA) via restriction digest. The vector contains a 6-fold His tag followed by mEGFP and a 14 amino acid linker sequence “GAPGSAGSAAGGSG”. The vector expressing mEGFP alone contains the linker sequence followed by a STOP codon.

TAF2 constructs with IDR deletions were generated using NEBuilder® to assemble different TAF2 fragments into the pENTR-TAF2 or pET-mEGFP plasmid. All expression constructs were validated by Sanger sequencing.

### Cell culture and generation of cell lines

Flp-In T-REx HeLa^92^ and U2OS cells were maintained in Dulbecco’s Modified Eagle medium (DMEM) containing 4.5 mg/L glucose and 1 mM sodium pyruvate (Thermo Fisher Scientific, 41966-029), supplemented with 10% v/v fetal bovine serum (FBS, Thermo Fisher Scientific, 10270106) and, for U2OS cells, 2 mM L-glutamine (Sigma, G7513). For selection of the Tet repressor and the Flp-In recombination target (FRT) site, cells were maintained in the presence of 5 µg/mL blasticidin (Invivogen, ant-bl) and 200 µg/mL zeocin (Thermo Fisher Scientific, 46-0509), respectively.

For the generation of stable doxycycline (Dox)-inducible HeLa cell lines, 200,000 cells were seeded in a 6-well plate format and pOG44 (Thermo Fisher Scientific, V600520) was co-transfected with a pcDNA5/FRT/TO plasmid containing the gene of interest in a 1:10 ratio using polyethylenimine “MAX” (Polysciences, 24765). After medium exchange 6 h post-transfection, cells were selected for the integration of the transgene 48 h post-transfection with 250 µg/mL hygromycin B (Thermo Fisher Scientific, 10687010). Selection medium was refreshed every 2–3 days and the hygromycin-resistant cell population expanded. The transgene expression was induced with 1 µg/mL doxycycline for 18–24 h and the cell lines were validated by Western blot, microscopy, and, if applicable, mass spectrometry.

HAP1 cells with C-terminal GFP knock-in in the endogenous SRRM2 locus (SRRM2-GFP truncations 0 and 10^52^) were a kind gift by Tugce Aktas (Max Planck Institute for Molecular Genetics, Berlin, Germany). We note that SRRM2-tr0 represents a C-terminal truncation of only four amino acids compared to the endogenous full-length protein (2752 aa) and we therefore refer to SRRM2-tr0 as the full-length protein. The cells were cultured in IMDM (Thermo Fisher Scientific, 12440–053) in the presence of 10% FBS (Thermo Fisher Scientific, 10270106) and 50 U/mL Penicillin-Streptomycin (Thermo Fisher Scientific, 15070-063). All cells were cultured at 37 °C in an incubator with 5% CO2 and humidity. All cell lines were routinely tested for mycoplasma contamination.

### Immunofluorescence of tissue culture cells and imaging

Approximately 50,000 cells were seeded into 8-well Ibidi μ-slides (Ibidi, 80826) and ectopic protein expression in Flp-In T-REx HeLa cells was induced with 1 μg/mL doxycycline (MP Biomedicals, 195044, Dox) minimum three hours post-seeding. 17–20 h after Dox induction, cells were either fixed or used for live-cell imaging.

For immunofluorescence staining, cells were fixed in 2% paraformaldehyde for 20 min at room temperature (RT), followed by three PBS washes and permeabilization with PBS containing 0.1% Triton X-100 for 3 min at RT. After two PBS washes, cells were incubated in 100 mM glycine/PBS for 20 min at RT. Cell were washed once with PBS and unspecific binding of the secondary antibodies blocked by incubation with 10% normal goat serum in PBS for 30 min at RT. Cells were incubated with primary antibodies diluted in 10% normal goat serum at 4 °C overnight, except for the MED1 antibody, which was incubated overnight at room temperature. Antibodies used were mouse monoclonal anti-sc35/SRRM2 (Sigma, S4045, 1:2,000), rabbit polyclonal anti-RBM25 (Sigma, HPA070713, 1:200), rabbit polyclonal anti-RBM14 (Sigma, HPA006628, 1:50), rabbit monoclonal anti-TAF1 (in-house/Abcam, cTAF1 177-4^93^)1:100), TAF3 (Abcam, ab188332, 1:500), MED1 (Abcam, ab64965, 1:500) and rabbit polyclonal anti-TAF2 (pAb 3038^53^, 1:100). Cells were washed three times in PBS and incubated for 1 h at RT with the appropriate secondary antibodies conjugated with fluorescent dyes (Invitrogen, Molecular probes, Alexa Fluor 488 or Alexa Fluor 568), diluted 1:500 in 5% normal goat serum/PBS. GFP-tagged proteins were imaged without additional antibody labeling. After three washes in PBS, cells were incubated in 2 μg/mL 4′,6-diamidino-2-phenylindole (DAPI, GeneCopoeia, C002) in PBS for 5 min at RT. After three PBS washes, cells were mounted using Ibidi mount (Ibidi, 50001). Images were acquired with a Zeiss LSM 880 confocal microscope with AiryScan detector, using the AiryScan LSM and super resolution modes with the 63x/1.4 NA oil Plan-Apochromat objective (Carl Zeiss) at a zoom range 2–4x. Excitation of the fluorophores was performed on separate tracks for each frame with a multiline argon 405-nm and 488-nm laser, and a 561-nm laser. For Z-stacks, images through the nuclei were taken at constant intervals and with optimal spacing. Images were processed using AiryScan Processing (ZEN Black, Carl Zeiss) with default settings and maximum intensity projections were created using FIJI (Image J, versions 1.51n or 1.53f51). In some images, the brightness was adjusted for visual clarity using FIJI; these settings were kept constant between comparative experimental designs, such as Dox-induction tests. Intensity profiles across nuclear speckles were obtained with FIJI.

For live-cell imaging, nuclei were stained with Hoechst 33342 (Thermo Fisher Scientific, H3570) and imaged with the LSM 880 with AiryScan microscope in a temperature-, CO2-, and humidity-controlled chamber with a heated stage. Images were processed as described for fixed-cell imaging.

### Fluorescence recovery after photobleaching (FRAP)

HeLa cells were seeded in glass bottom Ibidi dishes (Ibidi, 81158), and GFP–TAF2 expression was induced with 1 µg/mL doxycycline 20 h prior to photobleaching. FRAP was performed with a Zeiss LSM 800 microscope equipped with a 63x/1.4 NA oil objective lens using a 488 nm diode laser set to 70% bleach (iteration: 2, scan speed: 2). Regions of interest (ROIs) were defined as follows: bleached GFP–TAF2 foci region in the speckle (ROI1), the entire nucleus to correct for acquisition bleaching in the diffuse GFP–TAF2 pool (ROI2), and a background region outside the cell for correction (ROI3). 10 images were obtained pre-bleaching and 60 images were obtained every 2 s post-bleaching. Intensities of FRAP regions were extracted using ImageJ. FRAP data were analyzed with a customized Python script, where normalization was performed according to the double normalization method^94^. Normalized FRAP curve was fit using a single exponential function:

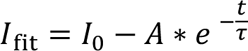

Where *I*_0_ represents the normalized intensity upon full recovery (plateau), *A* the difference *I*_0_ − *I_α_*, *I_α_* the normalized intensity at the first post-bleach time point *t*, and *τ* the characteristic time constant determined by the fit. The following formulas were used to compute the mobile fraction and the half-maximal recovery time^95^:

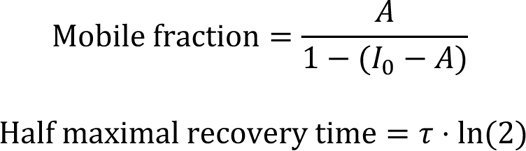

We calculated the mobile fraction to be 0.87 and the half-maximal recovery time 4.9 s. Quantification was carried out from FRAP curves derived from 13 nuclei; these experiments were repeated two times in total with similar results. The code for the FRAP analysis can be found at the Github link in the data and code availability section.

### Bacterial protein expression and purification

pET-mEGFP-TAF2 expression constructs with a 6-fold his-tag were transformed into BL21(DE3) and a 5 mL starter culture with 50 µg/mL kanamycin grown at 37 °C shaking overnight. 100 mL Luria Bertani (LB) medium supplemented with 50 µg/mL kanamycin were grown at 37 °C for *ca*. 3 h to reach OD 0.4. Cultures were transferred to 18 °C to reach OD 0.6 and protein expression was induced using 0.4 mM IPTG. The bacteria were pelleted at 4500 rpm for 20 min at 4°C and lysed in 30 mL lysis buffer (20 mM NaH2PO4, 300 mM NaCl, 20 mM imidazole, 0.1% Triton-X 100 at pH 8) supplemented with EDTA-free complete protease inhibitors (Roche, 56619600). Lysozyme was added to 1 mg/mL and lysate incubated for 20 min on ice. After three freeze/thaw cycles, the lysate was sonicated (12 cycles 10 sec pulse/10 sec pause with 30% amplitude) at 4 °C. Crude lysates were cleared by ultracentrifugation in Beckman ultracentrifuge (SW28 rotor) for 90 min at 25,000 rpm at 4°C.

Protein purification was performed on an ÄKTA pure 25 liquid chromatography system. The supernatant was adjusted to the conditions of the binding buffer (20 mM NaH2PO4, 500 mM NaCl, 20 mM imidazole at pH 8) and loaded on a nickel affinity chromatography column (1 mL HisTrap, GE Healthcare, 17-5247-01). Bound proteins were eluted in elution buffer (20 mM NaH2PO4, 500 mM NaCl, 500 mM imidazole, pH 8) and fractions analyzed via 12 % SDS-PAGE followed by Bio-Safe Coomassie Stain (Biorad,1610787). The peak fractions were combined and dialyzed in dialysis buffer (50 mM Tris pH 7.6, 500 mM NaCl, 10% glycerol, 1 mM DTT) at 4 °C overnight. The protein concentration was measured with Protein Assay Dye Reagent Concentrate (Biorad, 5000006) using a gamma-globulin protein standard.

### *In vitro* droplet assay and droplet image analysis

Dialyzed proteins were spun in a tabletop centrifuge at maximum speed at 4 °C for 5 min to remove precipitates. Proteins were desalted and concentrated to 100 uM in phase separation buffer according to^35^ (50 mM Tris pH 7.6, 125 mM NaCl, 10 % glycerol, 1 mM DTT) using Amicon Ultra-15 3 K or 30 K centrifugation units (Merck Millipore). The phase separation assay was performed in the presence or absence of 10% polyethylene glycol 8000 (PEG-8000, Promega, V3011) and the reaction mix loaded in a home-made imaging chamber consisting of a glass slide attached to a cover slip by two parallel strips of double-sided tape. Imaging of the droplets was performed immediately after mixing with a Zeiss confocal microscope LSM 800 with 63x/1.4 NA oil Plan-Apochromat objective (Carl Zeiss) and a zoom of 2.6x. Droplets that have settled on the coverslip were imaged and the same focal plane was used for the controls where no or few droplets were observed.

To analyze *in vitro* droplet assays, a custom-made Python script^74^, modified by Jonathan Henninger for image files from a Zeiss microscope, was used. The code for this analysis is available at the Github link in the data and code availability section. The droplets were segmented from the images on the following criteria: (1) an intensity threshold that was three s.d. above the mean of the image; (2) size thresholds (7 pixels minimum droplet size); and (3) a minimum circularity (circularity = 4π · (area)/(perimeter^2^)) of 0.5 (1 being a perfect circle). After segmentation, mean intensity for each droplet was calculated while excluding pixels near the phase interface. Hundreds of droplets identified in four independent fields of view were quantified.

### Cell fractionation for immunoblotting and GFP pulldown

HeLa cells expressing GFP–TAF2 and GFP(-NLS)-TAF2ΔIDR were seeded on 15 15-cm dishes, grown to 80-90% confluence and induced with 1µg/mL doxycycline for 24 h. Cells were harvested by trypsinization (Trypsin-EDTA, Sigma-Aldrich), washed three times in PBS and resuspended in 10% glycerol PBS before snap freezing in liquid nitrogen and storage at -80 °C. The cell pellet was thawed on ice and washed once with 10 mL PBS to remove glycerol. Nuclear and cytoplasmic extracts were obtained following a modified version of the Dignam and Roeder procedure^96^. The pellet was resuspended in five volumes of cold buffer A (10 mM Hepes-KOH pH 7.9, 1.5 mM MgCl2, 10 mM KCl) and incubated on ice for 10 minutes. After this initial lysis, the cells were centrifuged at 400 x *g* for 5 minutes at 4 °C, resuspended in buffer A complete (buffer A with 0.15% NP40, 0.5 mM DTT, 1 x Complete protease inhibitors, Roche), and dounced 40 times. The suspension was centrifuged at 2,840 x *g* for 15 minutes and the supernatant containing the cytoplasmic extract was cleared by centrifugation at maximum speed for 1 hour at 4 °C. For storage at -80 °C, 10% glycerol was added to the cytoplasmic extract. The nuclear fraction contained in the pellet was washed with 10 volumes of buffer A complete, resuspended in buffer C (420 mM NaCl, 20 mM Hepes-KOH pH 7.9, 20% v/v glycerol, 2 mM MgCl2, 0.2 mM EDTA, 0.5 mM DTT, 0.15% NP40, 1 x Complete protease inhibitors, Roche) and rotated at 4 °C for 2 hours. After rotation, the nuclei extracts were cleared by centrifugation at maximum speed for 1 hour at 4 °C. The extracts were snap frozen in liquid nitrogen and kept at -80 °C until the GFP pulldowns were set or until immunoblot.

### GFP pulldown and sample preparation for mass spectrometry

The GFP co-immunoprecipitations were essentially performed as previously described^97^. In brief, the total protein concentration in the nuclear and cytoplasmic extracts was measured using Protein Assay Dye Reagent Concentrate (Biorad, 5000006). 1 mg nuclear or 3 mg cytoplasmic extract per pulldown was incubated in triplicates with 15 μl of a 50% slurry of GFP-trap beads (ChromoTek, gta-20) or control agarose beads (ChromoTek, bab-20) in buffer C (300 mM NaCl, 20 mM Hepes-KOH pH 7.9, 20% v/v glycerol, 2 mM MgCl2, 0.2 mM EDTA, 1x Complete protease inhibitors (Roche), 0.5 mM DTT) containing 0.1% NP40 and 0.4% ethidium bromide and rotated overnight at 4 °C. The beads were washed twice with buffer C containing 0.5% NP40, twice with PBS containing 0.5% NP40, 0.5 mM DTT, and 1x Complete protease inhibitors (Roche), and twice with PBS followed by elution with 100 µl elution buffer (100 mM Tris-HCl pH 7.5, 2 M urea, 10 mM DTT) for 20 min, 1200 rpm, 22 °C. 50 mM iodoacetamide (IAA) was added to propitiate irreversible reduction of disulfide bonds. The samples underwent overnight on-bead digestion with 3.5 µl of trypsin (0.1 mg/mL, Promega) and were dissolved in 50 mM acetic acid in elution buffer. The samples were desalted through a C18 matrix (HyperSep SpinTip C-18, Thermo Fisher Scientific). 0.1% Trifluoroacetic acid (TFA; Thermo Fisher Scientific) in ddH2O was used as binding and washing solution and the tryptic peptides were then eluted from the tips with a releasing solution of 65% acetonitrile (LC-MS grade, Pierce) in ddH2O. The eluted tryptic peptides were lyophilized in a vacuum centrifuge (Christ, RVC 2-25 CD plus) with the following settings: chamber temperature 45 °C, rotation 600 min-1, vacuum pressure 15 mbar, safety pressure 20 mbar).

### MiniTurbo proximity labeling assay and sample preparation for mass spectrometry

For proximity biotinylation using miniTurbo-fusion proteins^54^, HeLa cells were seeded in five 15 cm dishes to approximately 50% confluency. miniTurbo–TAF expression was induced using 0.1 µg/mL doxycycline 16–20 h before cell harvest. Prior to extract preparation, the cells were incubated with 50 µM biotin (1 hour for miniTurbo–TAF2 and 3 hours for miniTurbo– TAF4). For extract preparation, the plates were placed on ice and the cells washed in ice-cold PBS. The cells were incubated with 10 mL of ice-cold buffer A (10 mM Hepes-KOH pH 7.9, 1.5 mM MgCl2,10 mM KCl, 1 mM DTT, 1x Complete Protease inhibitors, Roche) for 1–2 min, while swirling. The buffer was removed from the plates and 2 mL of buffer A containing 0.2% NP40 added to the cells while swirling. After 3–5 min, the cells were scraped from the plates and the lysate transferred to a pre-chilled 15 mL falcon tube. The cytoplasmic extract was separated from the nuclei by centrifugation at 2,800 rpm at 4°C for 6 min. For nuclear extract preparation, the nuclear pellet was resuspended in 3.6 mL of buffer C containing 0.2% NP40 (420 mM NaCl, 20 mM Hepes-KOH pH 7.9, 20% v/v glycerol, 2 mM MgCl2, 0.5 mM EDTA, 0.2% NP40, 1 mM DTT, 1x Complete Protease inhibitors, Roche) and the suspension tumbled at 4 °C for 45–75 min. The extract was centrifuged in an ultracentrifuge (Beckman, SW60 rotor) at 4 °C for 30 min and the supernatant transferred to a fresh tube. Aliquots of cytoplasmic and nuclear extracts were analyzed via immunoblot for biotinylation efficiency using streptavidin-HRP (Abcam, ab7403) and for miniTurbo–TAF protein expression using a rat anti-HA-HRP antibody (3F10, Merck, 12013819001). The protein concentration was determined with Protein Assay Dye Reagent Concentrate (Biorad, 5000006) using a gamma-globulin protein standard.

For the streptavidin and control pulldowns, 10 µL streptavidin bead slurry (GE Healthcare, 17-5113-01) per pulldown were either mock treated or blocked with 1 mM biotin by incubation in Buffer C (420 mM NaCl, 20 mM Hepes-KOH pH 7.9, 20% v/v glycerol, 2 mM MgCl2, 0.2 mM EDTA, 0.2% NP40) on a rotating wheel at 4 °C for 30–60 min. The beads were washed three times in Buffer C complete including 1x complete protease inhibitors (Roche) and 0.5 mM DTT. To prevent nucleic acid-mediated interactions, 2 µL ethidium bromide were added to each bead preparation. The beads were added to 1 mg nuclear extract and incubated on a rotating wheel at 4 °C for 60–90 min. The beads were spun at 2000 rpm, at 4 °C for 2 min and washed three times in buffer C, followed by three washes in PBS containing 1x Roche Complete protease inhibitors and 0.5 mM DTT, and three washes in PBS, followed by elution with 100 µl elution buffer (100 mM Tris-HCl pH 7.5, 2 M urea, 10 mM DTT). Samples were prepared for mass spectrometry analysis as described for GFP pulldowns.

### Nanoflow-HPLC tandem mass spectrometry (MS/MS)

Nanoflow-LC-MS/MS for sample analysis was performed with an Orbitrap Fusion Lumos mass spectrometer coupled to an Easy nano-LC 1200 HPLC (Thermo Fisher Scientific). For peptide separation by HPLC a gradient of increasing organic proportion (buffer A: 0.1 % formic acid, buffer B: 0.1 % formic acid in 80 % acetonitrile) was used with an C18 separation column of 25 cm length. The flow rate was 300nl/min. The mass spectrometer was operated in the data dependent mode with a TOP-10 method. Each MS scan was followed by a maximum of 10 MS/MS scans.

### Max Quant and Perseus analyses

The raw data files were analyzed using MaxQuant^98^ (version 1.6.5.0) with Andromeda peptide search against a reviewed human protein database without isoforms from Uniprot (released in June 2017; 20,188 entries) and the GFP protein sequence in case of the GFP-fusion protein pulldowns. We selected label-free quantification (LFQ) and match between run options. The intensity based absolute quantification (iBAQ) algorithm was activated for subsequent relative protein abundance estimation^99^. Peptide spectrum matching and relative protein quantitation were performed using a false discovery rate (FDR) of 1%. The resulting proteinGroups files are available on ProteomeXchange via the PRIDE^100^ server as outlined in the data and code availability section. The GFP–TAF2, GFP-TAF2ΔIDR and GFP-NLS-TAF2ΔIDR proteinGroups.txt file also contains data from GFP or agarose pulldowns from cytoplasmic HeLa cell fractions, which were processed as nuclear fractions, shown in Figs. 4A–B. The GFP pulldowns from the cytoplasmic fractions did not reveal GFP–TAF2 interactions with SRRM2 (data not shown).

The proteinGroups files with the specific (GFP or streptavidin pulldown) along with the respective control datasets (agarose or control pulldown) were loaded into Perseus^101^ (version v1.6.15.0). Potential contaminants and reverse hits were filtered out and LFQ intensity values log2 transformed to obtain a Gaussian normal distribution of the data. Groups based on specific, or control pulldowns were defined and identified proteins were accepted when measured three times within at least one of the two groups. Missing LFQ values were imputed based on the distribution of the LFQ values in the entire dataset (mode: entire matrix; settings: width=0.3; shift=1.8). Enriched proteins were identified by comparing the LFQ values of the specific and control triplicates using a two-tailed Student’s t-test. For the cut-off, the FDR was set to 1% and the threshold for significance (S0) was set to 1 for the miniTurbo–TAF analyses. The FDR was set to 0.1 or 1% and S0 to 1 in the GFP-pulldown analyses.

Relative protein abundances for the stoichiometry plots were obtained by subtraction of the average of non-specific iBAQ values from each of the iBAQ values of the specific pulldowns for the corresponding protein. The resulting values were divided by the values of a selected co-purifying protein for normalization and representation purposes.

### Co-immunoprecipitation assay for nuclear speckle protein SRRM2

Co-immunoprecipitation of GFP–TAF2 and SRRM2 was performed as described^52^. Briefly, 6 Mio cells were induced with 1 µg/mL doxycycline for 20 hours. The cells were washed twice in ice-cold PBS and resuspended in 600 uL NLB (1xPBS, 300 mM NaCl, 1% TritonX-100, 0.1% Tween-20) freshly supplemented with 1x Complete protease inhibitors (Roche), and 1x PhosSTOP (4906845001, Roche) and incubated on ice for 15 min. The lysate was centrifuged at 20,000 x *g* for 10 min at 4 °C. For the GFP pulldown, 25 µL of slurry of the GFP trap (Chromotek, #gta) were added to the lysate and rotated overnight at 4 °C. The beads were washed three times in NLB and bound proteins were eluted in 30 uL 1x NuPAGE LDS (Thermo Fisher Scientific, NP0007), supplemented with reducing agent, by boiling for 10 min at 80 °C.

### Whole-cell lysates

Cells were seeded in six-well dishes (800,000 cells/well) 4–6 hours prior to the induction of GFP–TAF protein expression with 1 ug/mL doxycycline for 18–24 hours before collection. Cell lysates were prepared in 2-fold sample buffer (320 mM Tris-HCl pH 6.8, 8% SDS, 40% glycerol, 0.10% bromophenol blue) and boiled at 95 °C for 5 min.

### Immunoblotting

Equal volumes of whole-cell lysate or 20–30 µg of cell extracts were separated via SDS-PAGE with the appropriate acrylamide concentration (8–12%) and transferred onto a 0.45 µm nitrocellulose (Amersham Protron, 10600002) or 0.2 µm or 0.45 µm PVDF membranes (Amersham Protron, 10600021 or Immobilon-P, Millipore, IPVH00010) for 1 h at 100 V in tris-glycine buffer with 10 % methanol. Membranes were blocked in blocking buffer (5% skimmed milk in TBST [tris-buffered saline, 0.1 % Tween-20] or PBST [phosphate-buffered saline, 0.1% Tween-20]) and incubated with the respective primary antibody in blocking buffer overnight at 4 °C. The appropriate HRP-conjugated secondary antibodies (Biorad) were diluted 1:10,000 in blocking buffer and the membranes incubated in this solution for 1 h at room temperature. Membranes were developed using ECL (Biorad Clarity ECL, 170-5061 or ECL Prime, Amersham, RPN2232) and analyzed on a ChemiDoc imaging system (BioRad) with subsequent processing with the Image Lab software (Biorad).

For immunoblotting SRRM2, lysates were prepared in NuPAGE LDS sample buffer (Thermo Fisher Scientific, NP0007) and reducing agent, boiled at 80 °C for 10 min and proteins separated using precast NuPAGE 4–12 % Bis-Tris gels (Thermo Fisher Scientific, NP0322PK2) at 80 V for *ca*. 3 hours. The proteins were transferred on a 0.45 µm PVDF membrane at 20 V for 16–18 hours at 4 °C in 10 mM CAPS pH 11 and 10 % methanol.

Antibodies used for immunoblotting in this study include rabbit polyclonal anti-GFP (ab290, Abcam, 1:2,500 or Sigma, G1544, 1:1000), mouse monoclonal anti-GFP (JL8, Clontech, 632381, 1:1000) mouse monoclonal anti-vinculin (SantaCruz, sc-73614, 1:1,000), mouse monoclonal anti-TBP (20C7, in house supernatant, 1:2 dilution in PBS), rabbit polyclonal anti-TAF2 (pAb 3038^53^, 1:500 in 0.3 % skimmed milk in PBS), mouse monoclonal anti-α-tubulin (Merck Millipore, DM1A, CP06), mouse monoclonal anti-sc35/SRRM2 (Sigma-Aldrich, S4045-.2ML, 1:1000), rabbit monoclonal anti-TAF1 (in-house/Abcam, cTAF1 177-4^93^), mouse monoclonal anti-lamin A/C (4C11, Cell Signaling, 4777).

Membranes were stripped to remove primary antibodies by incubation in 50 mM Tris-HCl pH 7.0, 2% SDS, 50 mM DTT for 15 min, washed in H2O and blocked again in blocking buffer for 1 hour at room temperature. HRP was inactivated with 1 mM sodium azide in blocking buffer prior to a second incubation with a primary antibody for Fig. S1B.

### Genome localization by (green)CUT&RUN

Genome localization analysis of GFP-tagged TAF2 full-length and deletion constructs was performed by greenCUT&RUN with the combination of enhancer-MNase and Lag16-MNase as described^68,69^. Briefly, cells were seeded into 10 cm dishes and GFP–TAF2 protein expression induced 18–20 h prior to cell harvest in Wash buffer (20 mM Hepes-KOH pH 7.5, 150 mM NaCl, 0.5 mM spermidine, 1x EDTA-free Complete Protease Inhibitor, Roche). After incubation with activated BioMag®Plus Concanavalin A magnetic beads (Polysciences, 86057-10) for 10 min at room temperature, cells were permeabilized in permeabilization buffer (Wash buffer containing 0.05% digitonin for HeLa cells) for 4 min at room temperature. 2 µg/mL enhancer-MNase and 2 µg/mL Lag16-MNase were added to the cells and incubated for 30 min at 4 °C while flicking. MNase was activated by incubation with 2 mM CaCl2 on wet ice for 30 min and fragments released during incubation at 37 °C for 30 min. The reaction was stopped using 2x Stop buffer (340 mM NaCl, 20 mM EDTA, 4 mM EGTA, 0.02% digitonin, 100 µg/mL RNAse A, 50 µg/mL glycogen) and incubated with 0.1% SDS and proteinase K. The DNA was isolated using phenol/chloroform extraction and precipitated with 100% ethanol at -20 °C overnight. The next day, DNA was washed in 100% ethanol and eluted in 1x TE buffer (10 mM Tris-HCl pH 8.0, 0.1 mM EDTA).

To localize endogenous TAF1, TAF2, TAF7, TBP, and RNA polymerase II, standard CUT&RUN^102^ was performed using the following antibodies (1 µg per CUT&RUN sample if not stated otherwise): rabbit monoclonal anti-TAF1 antibody (in-house/Abcam, cTAF1 177-4^93^), rabbit polyclonal anti-TAF2 antibody (pAb 3028^53^), rabbit polyclonal anti-TAF7 antibody (pAb 3475^103^), rabbit polyclonal anti-TBP (Abcam, ab28175), mouse monoclonal anti-RNA polymerase II CTD repeat (8WG16, Santa Cruz, sc-56767). For control CUT&RUN experiments, normal rabbit IgG (Cell Signaling, 2729, 1 µg per sample) was used.

For standard CUT&RUN and greenCUT&RUN, 10 pg of mononucleosomal *Drosophila* DNA was used as spike-in DNA for normalization purposes and sequencing libraries were prepared as described below.

### RNA isolation

Cells were seeded in triplicate in six-well dishes (500,000 cells/well) 4–6 hours prior to GFP-TAF protein induction with 1 µg/mL doxycycline (Dox) for 17.5 hours before collection. Control HeLa Flp-In T-Rex cells were also subjected to 1 µg/mL Dox treatment. Cells were harvested and total RNA isolated using an RNA isolation kit (RNeasy Mini kit, Qiagen, 74106). 8 µg of total RNA were treated with DNaseI and purified according to the manufacturer’s protocol (TURBO DNA-free kit, Thermo Fisher Scientific, AM1907).

### Library preparation and deep sequencing

For CUT&RUN^102^ and greenCUT&RUN^68,69^, purified DNA fragments were subjected to library preparation with NEB Next Ultra II (NewEngland Biolabs, E7645L) and NEB MultiplexOligo sets I/II/III/IV (NewEngland Biolabs, E7500L) without size selection according to the manufacturer’s protocol with modifications: the end prep was 60 min at 50 °C, and the PCR amplification step was performed with 10 s annealing/extension time and 15 cycles according to^102^. The DNA concentration was determined using using a Qubit instrument (Invitrogen, USA) and the size distribution of the fragments was analyzed on an Agilent Bioanalyzer (DNA 100 assay). CUT&RUN libraries (10–20 Mio reads per sample and 1/3 of reads for the controls) were sequenced on a NovaSeq 6000 (Illumina) paired end 100 bp at the Deep Sequencing Facility (Max Planck Institute for Immunobiology and Epigenetics, Freiburg, Germany) or at Novogene (Cambridge, United Kingdom). Some of the greenCUT&RUN libraries were sequenced on the MiniSeq (Illumina, 6–8 Mio reads per sample and 1/3 of reads for the controls).

For RNA sequencing, libraries were prepared using the Illumina Stranded mRNA Prep kit (Illumina, 20040534) with a starting amount of 100 ng high quality total RNA. Within the procedure Oligo(dT) magnetic beads purify and capture polyadenylated RNAs. The purified PolyA+ RNA molecules are fragmented and copied into first strand complementary DNA (cDNA) using reverse transcriptase and random primers. In a second strand cDNA synthesis step, dUTP replaces dTTP to achieve strand specificity. The final steps add adenine (A) and thymine (T) bases to fragment ends and ligate adapters (Y-shaped, universal anchors). The resulting products are purified and selectively amplified (unique dual index primers) for multiplexed sequencing on an Illumina system. Libraries were sequenced to 50 million reads per sample on a NovaSeq 6000 (Illumina) paired end 100 bp and reads processed using bcl2fastq version: 2.20.0.422.

### Bioinformatic analysis of proteomic and genomic data

#### Protein sequence analysis

For the amino acid composition and charge plots (Figs. 3A and S4C), a custom-made R script was used to create a binary heatmap together with the related average electrostatic charge. The TAF2 and SRRM2 protein sequences and the charge info were obtained from Uniprot and through the Emboss algorithm (http://www.bioinformatics.nl/cgi-bin/emboss; protein properties “charge”), respectively. The algorithm assigns to residues ’D’ and ’E’ a charge of - 1, to residues ’K’ and ’R’ a charge of +1, and to the residue ’H’ a charge of +0.5. The mean charge was calculated across a sliding window of 20 residues. The code to reproduce this plot can be found under the respective Github link in the data and code availability section.

The predictions of the TAF2 intrinsically disordered regions were performed using the online tools Predictor of Natural Disordered Regions (PONDR, http://www.pondr.com) with VSL2 algorithm^56^ and DISOPRED3^57^ (http://bioinf.cs.ucl.ac.uk/psipred/). NLStradamus^104^ was used to identify nuclear localization signals (NLSs) in the TAF2 protein sequence.

#### greenCUT&RUN and CUT&RUN analysis

Initially, reads underwent quality control filtering using Trim-galore (v0.6.4) with default parameters (http://www.bioinformatics.babraham.ac.uk/projects/trim_galore/). Subsequently, the alignment was carried out using bowtie2 (v2.3.5.1) (https://bowtie-bio.sourceforge.net/bowtie2/index.shtml) with the following options: -dovetail -local -very-sensitive-local -no-unal -no-mixed -no-discordant -I 10 -X 700^68^. The alignment for human was performed against the hg38/GRCh38.p13 version of the genome (https://www.gencodegenes.org/human/), while the spike-in reads were aligned to the BDGP5 version of the Drosophila genome sequence (EnsEMBL v.75, www.ensembl.org). The total number of aligned reads, representing both human and spike-in, was calculated using the Flagstat program of samtools (version 1.10). Peaks were identified using HOMER^105^ with default parameters, except for the inclusion of –C 0. Normalization of reads during peak calling was conducted using spike-in, as previously described^68^. Peaks in the experiment were called against control samples. Peak files from HOMER were uploaded to the Galaxy platform^106^ and the public server usegalaxy.eu was used for data analysis. To merge peaks, bedtools intersect intervals (v 2.30.0+galaxy1) was used with 1 bp required overlap. Peaks were annotated using the HOMER annotatePeaks function.

The generated bam files from the experiment were also uploaded for further analysis to the Galaxy platform. Spike-in normalized genome coverage (BigWig) files were created with DeepTools^107^ bamCoverage (v3.5.1.0.0) using a blacklist region file for hg38, bin size 50 bases and 1x normalization (RPCG). The scaling factor was calculated as follows: 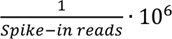 for CUT&RUN and 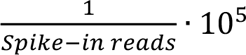 for greenCUT&RUN. Spike-in normalized coverage tracks were visualized using IGV (v2.12.3). Heatmaps were generated in Galaxy using DeepTools computeMatrix (v3.5.2+galaxy0) and plotHeatmap (v3.5.1.0.1) and profile plots were generated using DeepTools computeMatrix and plotProfile (3.5.2+galaxy0).

#### RNA-seq analysis

RNA-seq pre-processing was performed on the FRT (control), GFP-NLS-TAF2ΔIDR, and GFP–TAF2 FASTQ samples to produce a joined count matrix. Downstream differential gene analysis then contrasted each sample condition against the other to produce differentially expressed (DE) genes. The pre-processing utilized the Rsubread v2.12.0^108^. Raw FASTQ reads were aligned to the Homo Sapiens (hg38) reference genome, with the resulting BAM files then being quantified at the exon feature level to produce raw feature tables which were then joined into a single count matrix. The downstream analysis was run via the R package RNA-seq Helper^109^ based on DESeq2 v1.40.2^110^. Using the “run_deseq” function, quality control was run on the matrix; filtering out any genes with less than 30 total counts across all samples, as well as any that did not appear in more than 4 samples. DESeq2 normalization, along with a variance-stabilizing transform (VST) to correct for unwanted variation on the data was also performed via the “pca_and_matrices” function, resulting in principal component analysis (PCA) plots showing good clustering of all samples by condition. The plot showing the Differential Expression Index (DEI, Fig. 6B) was generated using a custom-made R script. DEI is defined as the product of the sign of the log2 fold change (FC) and the log of its *P* value: DEI*=*sign[log(FC)]⋅[−log10(*P* value) according to^111^.

#### Alternative splicing (AS) analysis

AS analyses on the RNA-seq data sets were performed using vast-tools v2.5.1^112^. Briefly, the software quantifies inclusion/exclusion levels of AS events (exons, introns, alternative splice sites), by calculating the percentage of sequence inclusion (PSI) as: inclusion/(inclusion+exclusion) x 100. To increase sequencing coverage and avoid biases towards highly expressed genes the three independent replicates of each condition were merged prior to the analysis to create a super sample. Differentially spliced events were identified using three different criteria: (i) |ΔPSI| >= 15% between the super sample and control; (ii) *P*-value (*t*-test) <= 0.01 and an average |ΔPSI| >= 10 between three individual replicates (iii) Bayesian inference using a Minimum Value (MV) of |ΔPSI| at 95% confidence >= 15%. The code to reproduce the analysis and figures can be found at the Github link in the data and code availability section.

#### Gene ontology analysis of mass spectrometry, RNA-seq data, and AS analysis

The Gene Ontology (GO) studies for mass spectrometry data in Figs. S2B and C and RNA-seq data in Fig. S6C have been performed as described^111,113^. Briefly, several runs of clusterProfiler^114^ were performed, and each run was defined by an increasingly stringent selection of genes based on their *P* values resulting from the MS analysis with Perseus^101^ or RNA-seq differential expression analysis. This resulted in several GO *P* values for each category. The reported final *P* values were obtained by averaging the negative log10 of the obtained GO *P* values. The aim of this procedure was to reduce the effect of arbitrariness in the definition of significantly enriched proteins or DE genes in the MS pulldown and RNA-seq data set, respectively.

GO analysis on the AS data was performed with GOfuncR^115^ using the whole human genome assembly hg38 (v88) as background. GOfuncR was chosen since its output can be directly processed by slimGO, a routine of the R package DMRrichR^116–118^, which generates a summary of selected GO terms. The multiple threshold method described above was not applied to the AS data set due to the limited number of differentially spliced events in each group.

#### Multiple protein sequence alignment

Alignment of 10 full-length protein sequences of TAF2 from different species (Figs. 3B and S3C): Human: Q6P1X5, mouse: B9EJX5, cow: A0A3Q1MUH2, western clawed frog: ENSXETP00000027252, bird: ENSPMJP00000014835, zebrafish: Q32PW3, honeybee: A0A088AQA5, fruit fly: Q24325, budding yeast: P23255, arabidopsis: Q8LPF0. The alignment was performed using the Jalview 2 bioinformatics tool^119^ and Clustal Omega^120^. Residues were colored according to conservation following the Blosum62 score.

## List of supplemental tables

Supplemental table S1: MS data and iBAQ values from proximity labeling experiments in HeLa cells expressing miniTurbo–TAF2 and miniTurbo–TAF4 (related to Fig. 2G)

Supplemental table S2: Results of *in vitro* phase separation screen of IDRs in TFIID subunits

Supplemental table S3: MS data underlying Volcano plots and iBAQ values from GFP–TAF2, GFP-TAF2ΔIDR, and GFP-NLS-TAF2ΔIDR GFP pulldowns from HeLa cell nuclear extracts (related to Figs. 4A and B)

Supplemental table S4: Gene lists from Upset plot in Fig. S4B

Supplemental table S5: MS data underlying Volcano plots and iBAQ values from SRRM2-tr0 and SRRM2-t10 GFP pulldowns from HAP1 nuclear extracts (related to Figs. 4E and F)

Supplemental table S6: RNA-seq data differential expression analysis of HeLa cells expressing GFP–TAF2 *versus* control or GFP-NLS-TAF2ΔIDR *versus* control (related to Figs. 6B and S6A–B)

Supplemental table S7: Differentially spliced exons and introns from AS analysis of GFP– TAF2 expressing HeLa cells *versus* control (related to Fig. 6C) and GFP-NLS-TAF2ΔIDR expressing HeLa cells *versus* control (related to Fig. 6D)

## Acknowledgments

We would like to acknowledge the Lighthouse Core Facility (Medical Center–University of Freiburg, Germany) for their assistance with confocal microscopy and the Deep Sequencing Facility (Max Planck Institute for Immunobiology and Epigenetics, Freiburg, Germany), for sequencing CUT&RUN samples, and RNA-seq library preparation and sequencing. We acknowledge the support of the Freiburg Galaxy Team, funded by the Collaborative Research Centre 992 Medical Epigenetics (DFG grant SFB 992/1 2012) and the German Federal Ministry of Education and Research BMBF grant 031 A538A de.NBI-RBC. We thank Carsten Schwan (Medical Center–University of Freiburg), Svenja Ulferts (Medical Center–University of Freiburg), and Sophie Weyrauch (University of Freiburg) for their help with FRAP imaging. We thank Jonathan Henninger (Carnegie University, Pittsburgh, USA) for modifying their script for the *in vitro* droplet assay analysis and Richard Young (Massachusetts Institute of Technology, Cambridge, USA) for sending us the pET-mEGFP plasmid. We thank Sven Diederichs (Medical Center–University of Freiburg, German Cancer Research Center, Heidelberg, Germany) for helpful discussions and introduction to the R-DeeP dataset. We are grateful to Tugce Aktas (Max Planck Institute for Molecular Genetics, Berlin, Germany) for providing us with the HAP1 cell lines and Laszlo Tora (IGBMC, Strasbourg, France) for the anti-TAF2 (pAb 3038) and the anti-TAF7 (pAb 3475) polyclonal antibodies. We thank Raphael Reuten, Jonathan Henninger, Laszlo Tora, and Andrea Bernardini for critical comments and reading of the manuscript and acknowledge all members of the Arnold laboratory for helpful discussions.

## Author contributions

T.B. discovered the TAF2-nuclear speckle association, conceptualized the project, and designed the study. T.B. performed most experiments and data analyses in Figs. 1–6. P.K.M.S. made the GFP–TAF4 cell line, performed immunoblotting in Figs. S1B and S4A and prepared samples for mass spectrometry in Fig. 4. N.A. performed the alternative splicing analysis in Fig. 6. J.K. performed the FRAP experiment and analysis in Figs. 3G and S3I. S.N. provided the analysis pipeline for CUT&RUN data mapping and peak calling, helped analyze RNA-seq data in Fig. 6, and performed CUT&RUN experiments: GFP-NLS-TAF2-d1142-1171_rep1 and GFP-TAF2_rep1 in Fig. 5. A.P. helped analyze RNA-seq data, performed all gene ontology analyses and visualized data in Figs. S2B–C, 3A, 3E, S3A, 4A, 4E, S4B, S5C, 6B–E, and S6. M.T. performed RNA-seq differential expression analysis. M.L.B. performed quantitative mass spectrometry in Fig. 4. S.K. performed four CUT&RUN experiments in Figs. 6 and S6. T.H. supervised J.K., gave advice and helped interpret the data. S.J.A. gave advice on experimental design and data interpretation. T.B. wrote the manuscript and all authors discussed the results and commented on the manuscript text.

## Resource availability Lead contact

Further information and requests for resources and reagents should be directed to and will be fulfilled by the lead contact, Sebastian J. Arnold.

## Materials availability

Plasmids and cell lines generated during this study are available from the lead contact upon request.

## Data and code availability

The CUT&RUN and RNA-seq data in this study have been deposited in the Gene Expression Omnibus (GEO) database under accession code GSE254081. The mass spectrometry proteomics data have been deposited to the ProteomeXchange Consortium (http://proteomecentral.proteomexchange.org) via the PRIDE partner repository^100^ with the dataset identifier PXD049117. The imaging data supporting this study have not been deposited in a public repository due to file sizes but are available from the corresponding authors upon request. The code used for alternative splicing analysis is available at https://github.com/Ni-Ar/TAF2_OE_AS (reviewer link will be provided upon request). The R code used to produce the amino acid composition and charge plot is available at https://github.com/andreaprunotto/amino_acid_composition_plot. The customized Python script for FRAP data analysis can be found at https://gitlab.physchem.uni-freiburg.de/ak.hugel/frap-analysis. The customized Python script for *in vitro* droplet assay analysis can be found at https://github.com/tanbhuiyan/In-vitro-droplet-analysis. Any additional information required to reanalyze the data reported in this paper is available from the lead contact upon request.

## Figures

**Figure S1:**
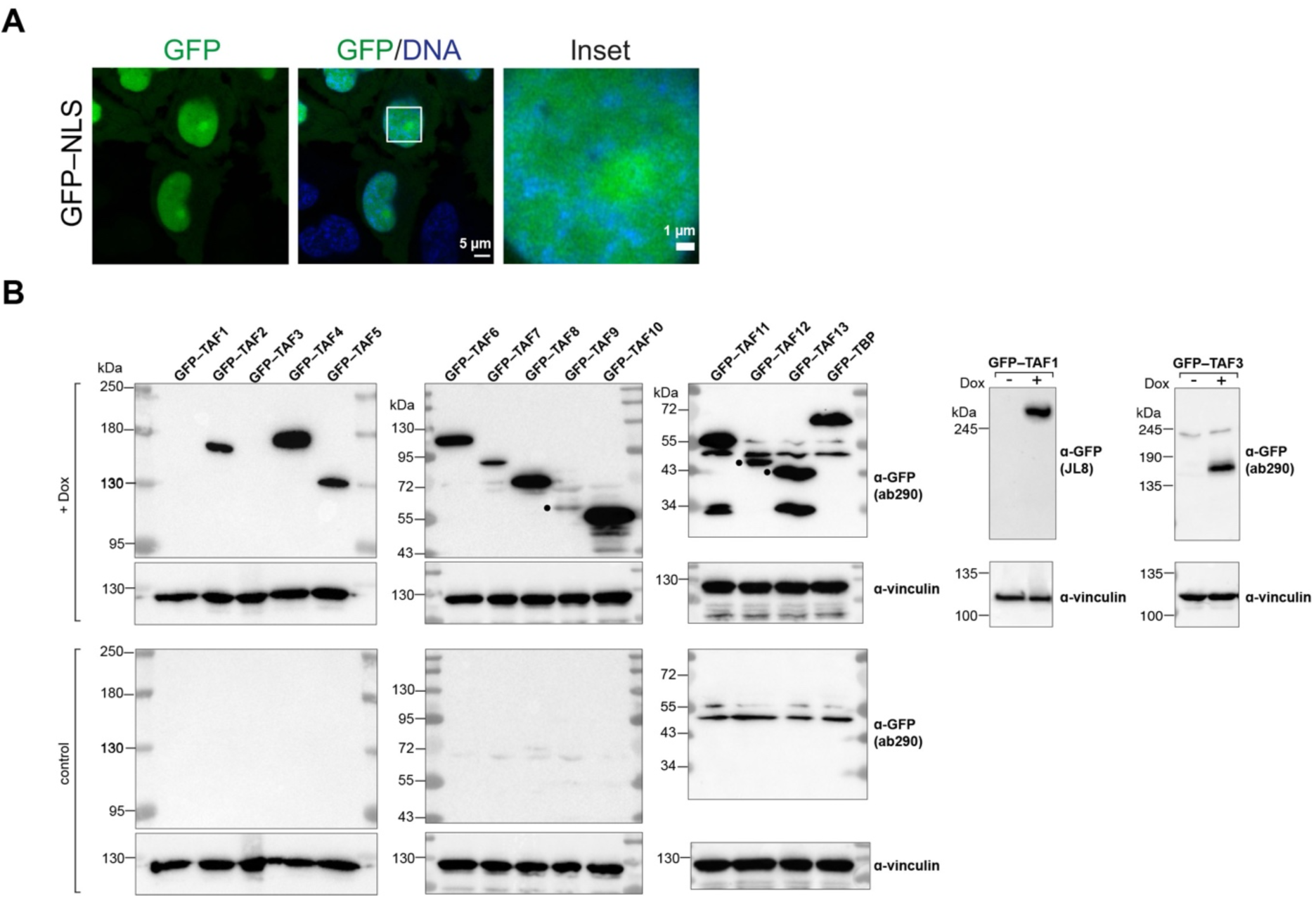
Live-cell imaging and protein expression of GFP-tagged TFIID subunits. **(A)** Live-cell confocal microscopy (AiryScan) images of HeLa cells expressing a GFP-tagged simian virus 40 (SV40) nuclear localization signal (NLS). Single confocal planes are shown. DNA was stained with Hoechst 33342. Scale bars, 5 µm and 1 µm (inset). **(B)** Immunoblot of whole-cell lysates from HeLa cells imaged in Fig. 1B. Protein expression was induced for 20 h with doxycycline (Dox). GFP–TAF1 and GFP–TAF3 expression was not detected and thus immunoblotted separately. Filled circles indicate bands corresponding to the expected molecular weight of the GFP-fusion proteins. Uncropped images can be found in the source data related to Fig. S1B.

**Figure S2:**
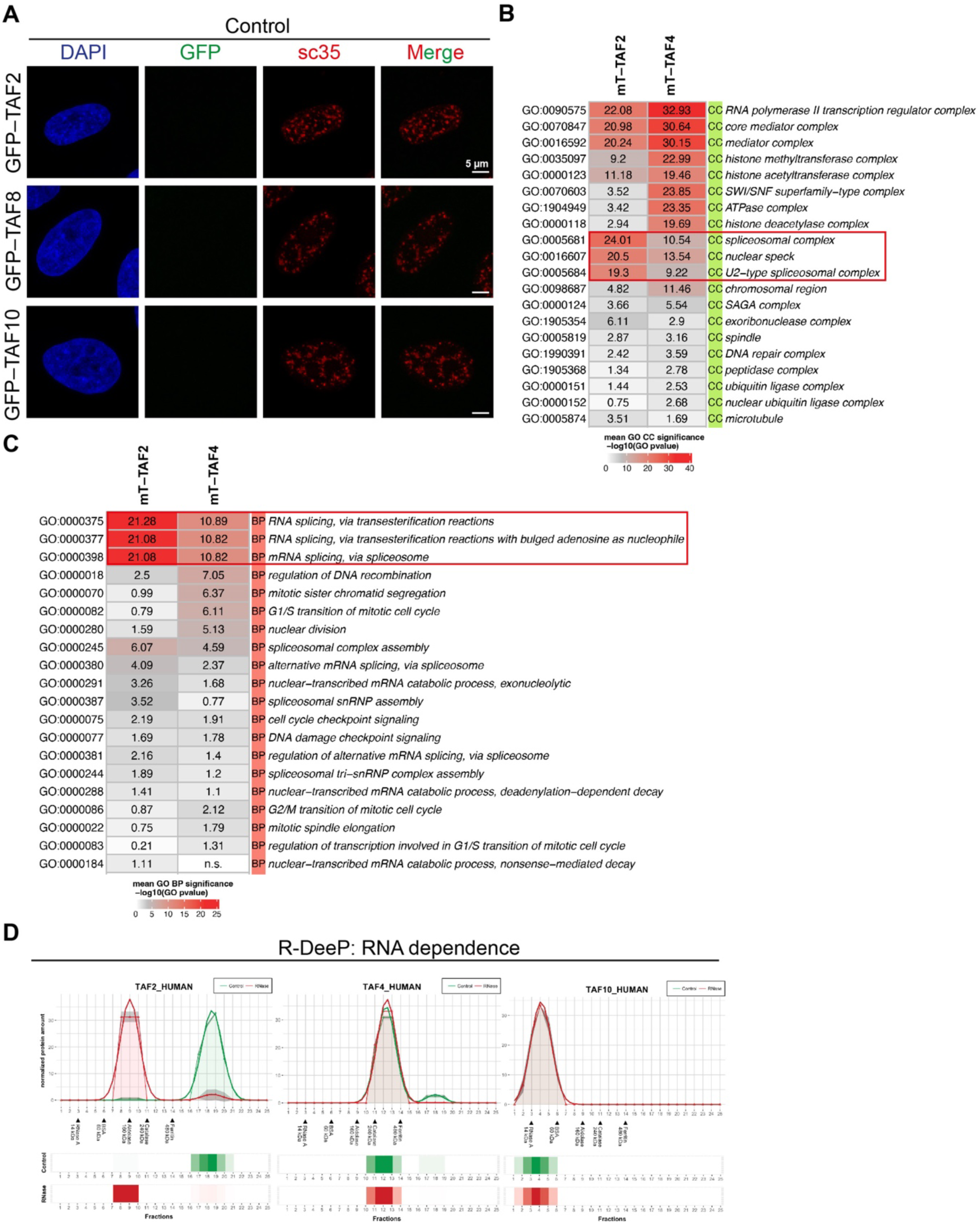
GFP–TAF co-staining experiments and RNA dependence. **(A)** Fixed-cell confocal microscopy (AiryScan) images of uninduced (control) HeLa cell lines harboring the transgene for expression of the GFP-fusion proteins of the 3TAF complex (TAF2, TAF8, and TAF10) as in Fig. 2A. Maximum intensity projections from Z-stacks are shown. **(B)** Gene ontology (GO) analyses of specific interactors identified by quantitative mass spectrometry in miniTurbo biotinylation experiments for TAF2 and TAF4. Categories of cellular components **(B)** and biological processes **(C)** are shown. mT, miniTurbo. **(D)** R-DeeP profiles for the TFIID subunits TAF2, TAF4, and TAF10. RNA-dependent proteins interact with RNA-binding proteins and/or RNA. The profiles of proteins eluted from sucrose density gradients with and without RNase treatment were obtained from the website https://r-deep.dkfz.de [1].

**Figure S3:**
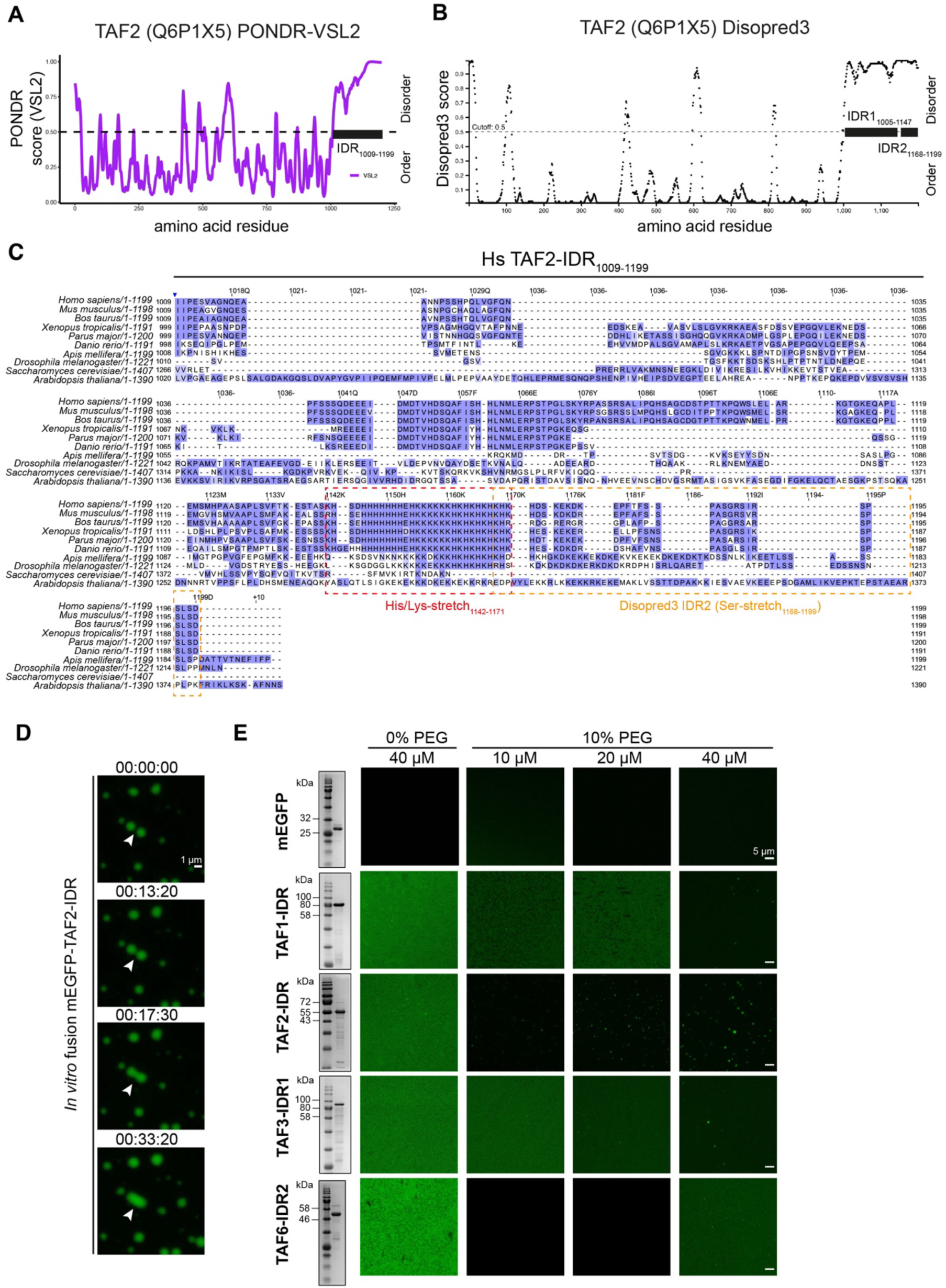
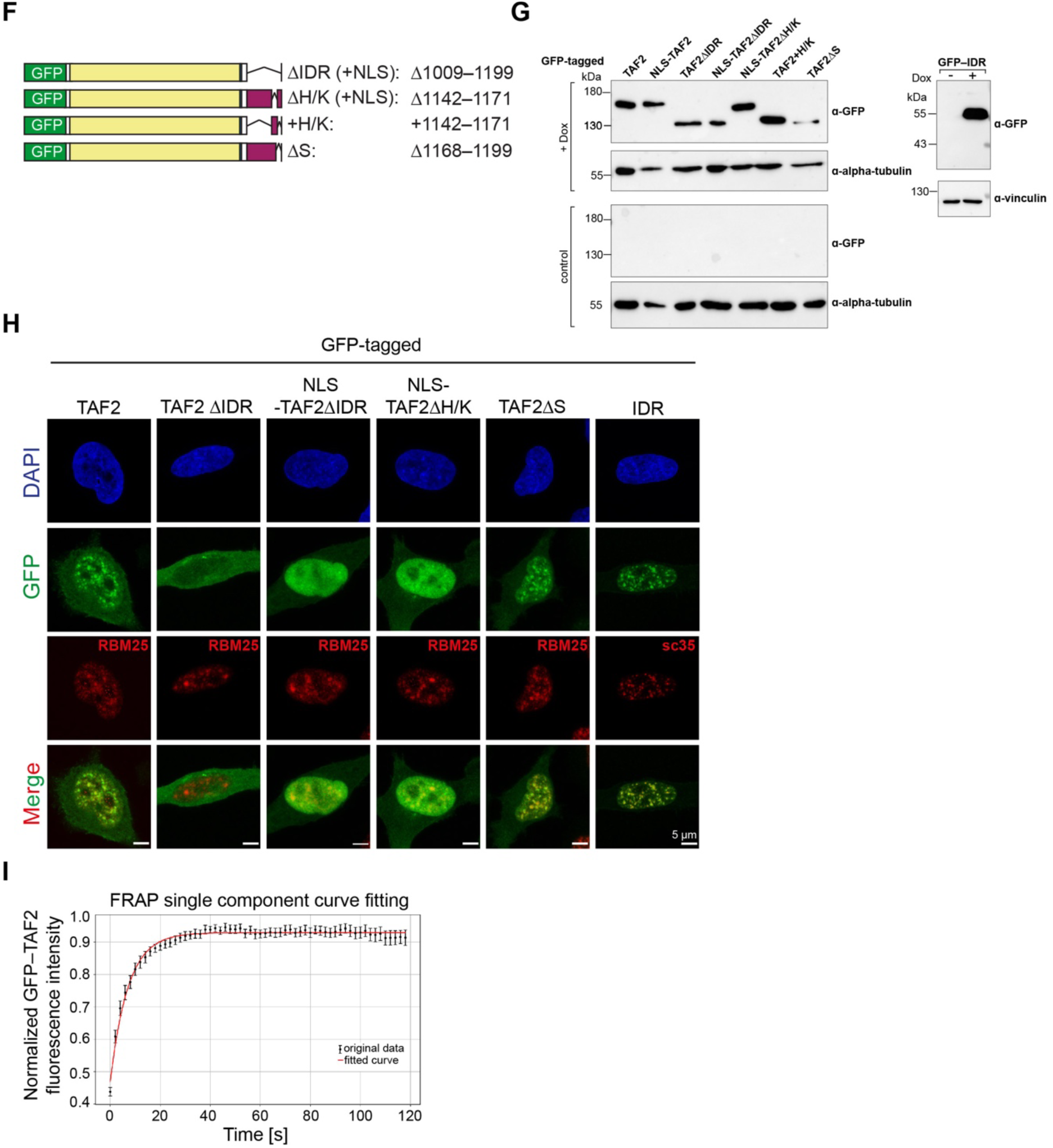
TAF2 protein sequence analyses, TFIID *in vitro* droplet assays, GFP–TAF2 deletion construct imaging and FRAP analyses. **(A)** Intrinsically disordered domain (IDR) prediction using PONDR [2] (http://www.pondr.com) or **(B)** DISOPRED3 [3] (http://bioinf.cs.ucl.ac.uk/psipred/) **(C)** Multiple protein sequence alignment created with Jalview [4]. Alignment of 10 full-length protein sequences of TAF2 from different species. *Homo sapiens* (Q6P1X5), *Mus musculus* (B9EJX5), *Bos taurus* (A0A3Q1MUH2), *Xenopus tropicalis* (ENSXETP00000027252), *Parus major* (ENSPMJP00000014835), *Danio rerio* (Q32PW3), *Apis mellifera* (A0A088AQA5), *Drosophila melanogaster* (Q24325), *Saccharomyces cerevisiae* (P23255), *Arabidopsis thaliana* (Q8LPF0). Residues were colored according to conservation following the Blosum62 score. The numbering corresponds to the human protein sequence. **(D)** Confocal images from an *in vitro* droplet assay observed for 33 min and 20 sec. The white arrowhead points to a droplet fusion event. The droplets imaged have sedimented on the coverslip (focal plane). **(E)** Confocal images from *in vitro* droplet assays using different TAF-IDR fusions with monomeric EGFP (mEGFP). Scale bar, 5 µm. Coomassie gels for each mEGFP-TAF-IDR fusion protein purification are shown on the left. **(F)** Schematic of the GFP–TAF2 deletion constructs used for live-cell imaging in Fig. 3F. **(G)** Immunoblot of whole-cell lysates from HeLa cells shown in Fig. 3F. Protein expression was induced with doxycycline (Dox). Uncropped images can be found in the source data related to Fig. S3G. **(H)** Fixed-cell confocal microscopy (AiryScan) images of HeLa cell lines expressing GFP–TAF2 deletion constructs and co-stained with the nuclear speckle markers RBM25 or sc35. Maximum intensity projections from Z-stacks are shown. Scale bar, 5 µm. **(I)** Curve fitting of FRAP data shown in Fig. 3G.

**Figure S4:**
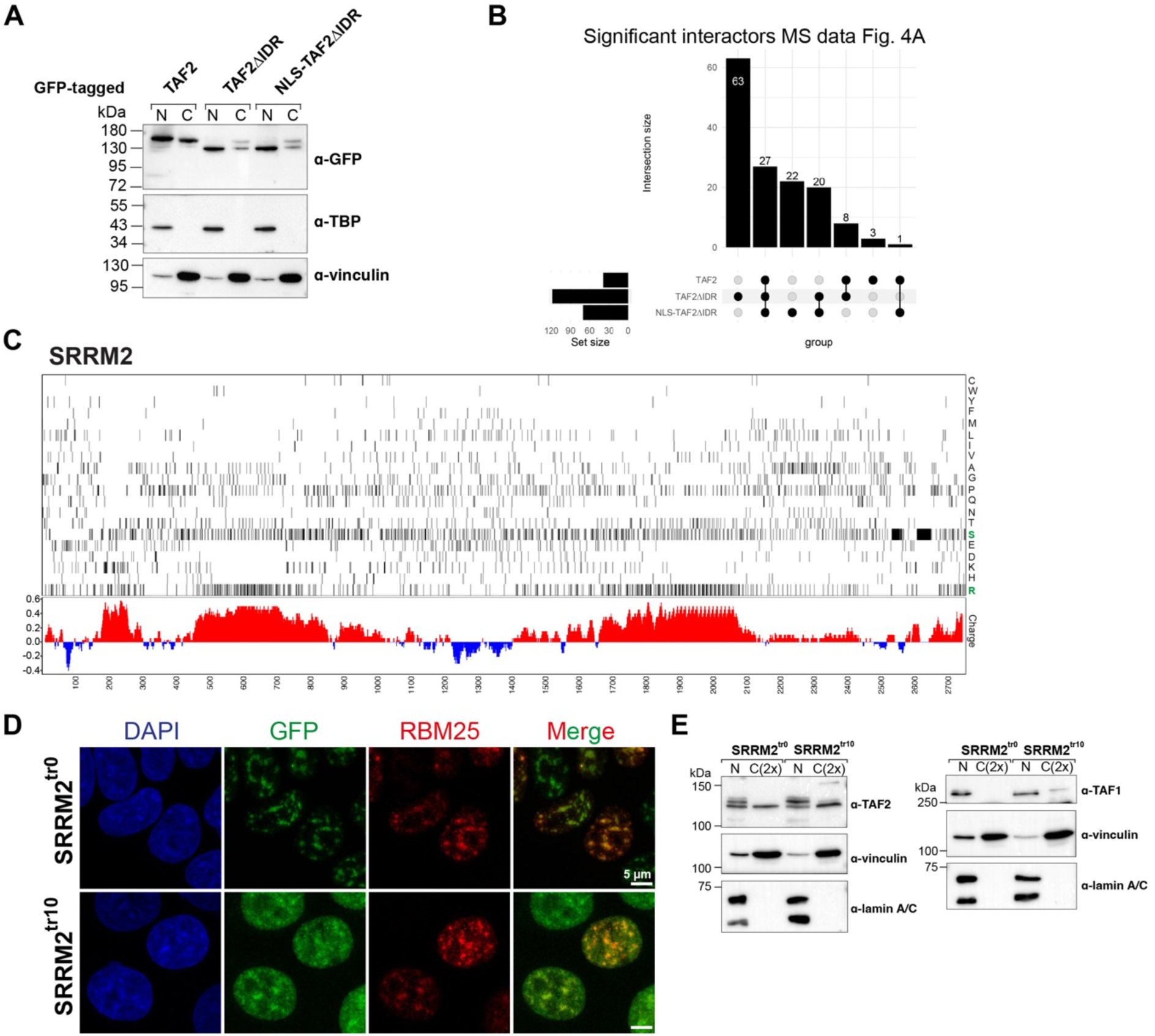
HeLa and HAP1 cell fractionations, MS data analysis and SRRM2 sequence analysis. **(A)** Subcellular fractionation of HeLa cells expressing GFP-tagged TAF2, TAF2ΔIDR and NLS-TAF2ΔIDR followed by immunoblot analysis. TBP and vinculin were used as loading and fractionation controls. The amount of loaded cytoplasmic (C) extract was equal to the amount of the nuclear (N) extract counterpart. Uncropped images can be found in the source data related to Fig. S4A. **(B)** Upset plot showing the intersections of common and unique significant interactors from the Volcano plots in Fig. 4A. (**C**) Human SRRM2 (Q9UQ35) amino acid composition plot. The heatmap represents the position of SRRM2 residues (rows: amino acids; columns: position) together with the related average electrostatic charge (red: positive charge; blue: negative charge) calculated with a sliding window of 20 amino acids. (**D**) Fixed-cell confocal microscopy (AiryScan) images of HAP1 cell lines expressing SRRM2^tr0^ or SRRM2^tr10^ co-stained with nuclear speckle marker RBM25. Z-stacks are shown as maximum intensity projections. (**E**) Subcellular fractionation of HAP1 cells [5] expressing SRRM2^tr0^ or SRRM2^tr10^ run on a 6% polyacrylamide gel followed by immunoblot analysis of endogenous TAF1 (left panel) or TAF2 (right panel). Lamin A/C and vinculin were used as loading and fractionation controls. The amount of loaded cytoplasmic (C) extract was double the amount of the nuclear (N) extract counterpart. Uncropped images can be found in the source data related to Fig. S4A.

**Figure S5:**
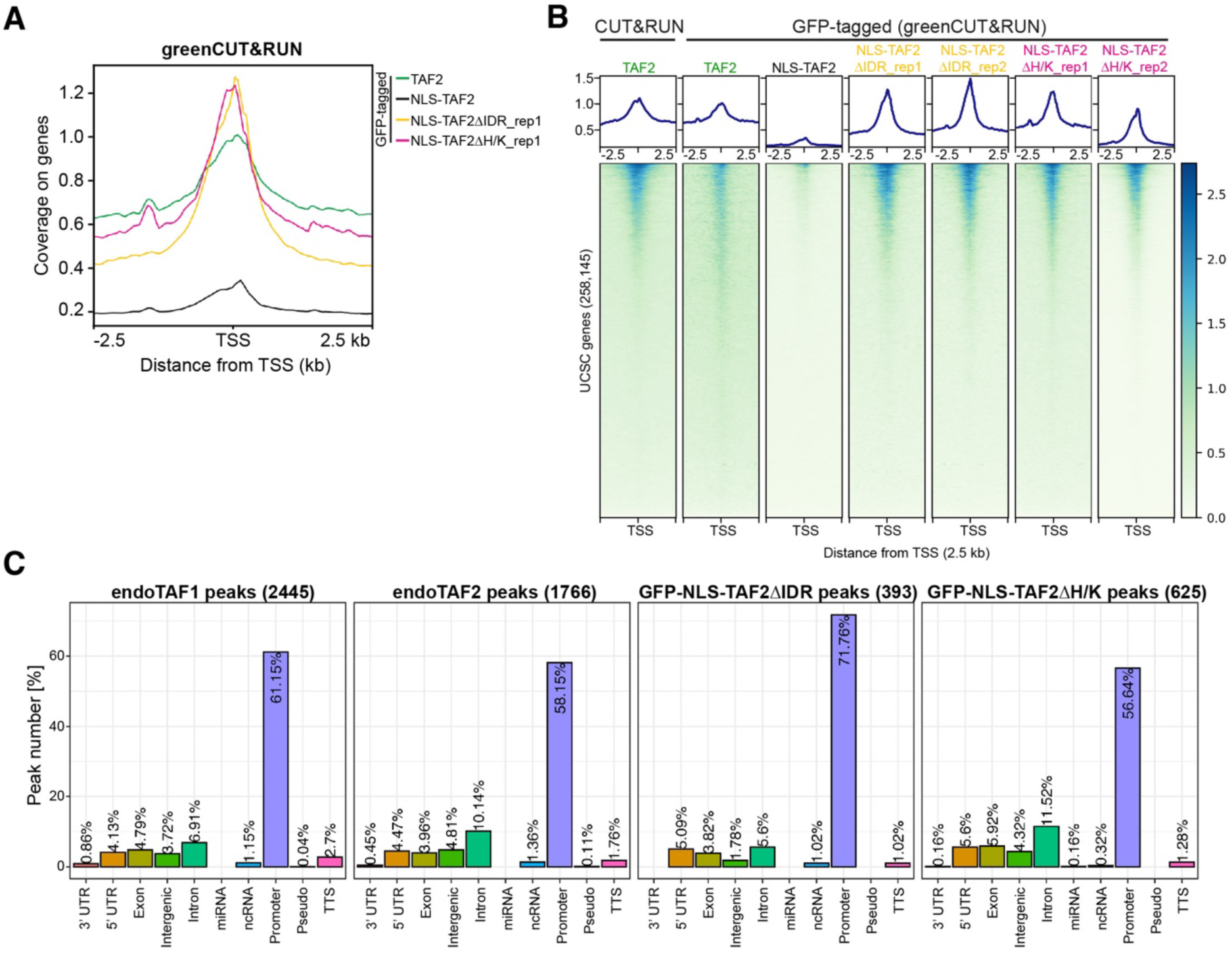
CUT&RUN coverage over transcription start sites and CUT&RUN peak analyses. **(A)** Profile plot displaying greenCUT&RUN coverage of GFP-tagged TAF2 full-length or deletion constructs inducibly expressed in HeLa cells. Spike-in normalized coverage coverage (1x normalization, RPGC) is centered on transcription start sites (TSS) of human coding and non-coding genes and the coverage 2.5 kilobases (kb) up- and downstream is shown. **(B)** Heatmaps displaying CUT&RUN coverage of endogenous TAF2 in control HeLa cells and greenCUT&RUN coverages of GFP-tagged full-length TAF2 and deletion constructs. The spike-in normalized coverage is centered on the TSS of coding and non-coding genes and the coverage 2.5 kb up- and downstream is shown. rep1 and rep2 denote biological replicates 1 and 2, respectively. **(C)** TAF2 peaks mostly fall into promoter regions. Comparison of peak annotations from peaks of endogenous TAF1 and TAF2 CUT&RUN and greenCUT&RUN of GFP-NLS-TAF2ΔIDR and GFP-NLS-TAF2ΔH/K (common peaks from biological duplicates in **(B)**.)

**Figure S6:**
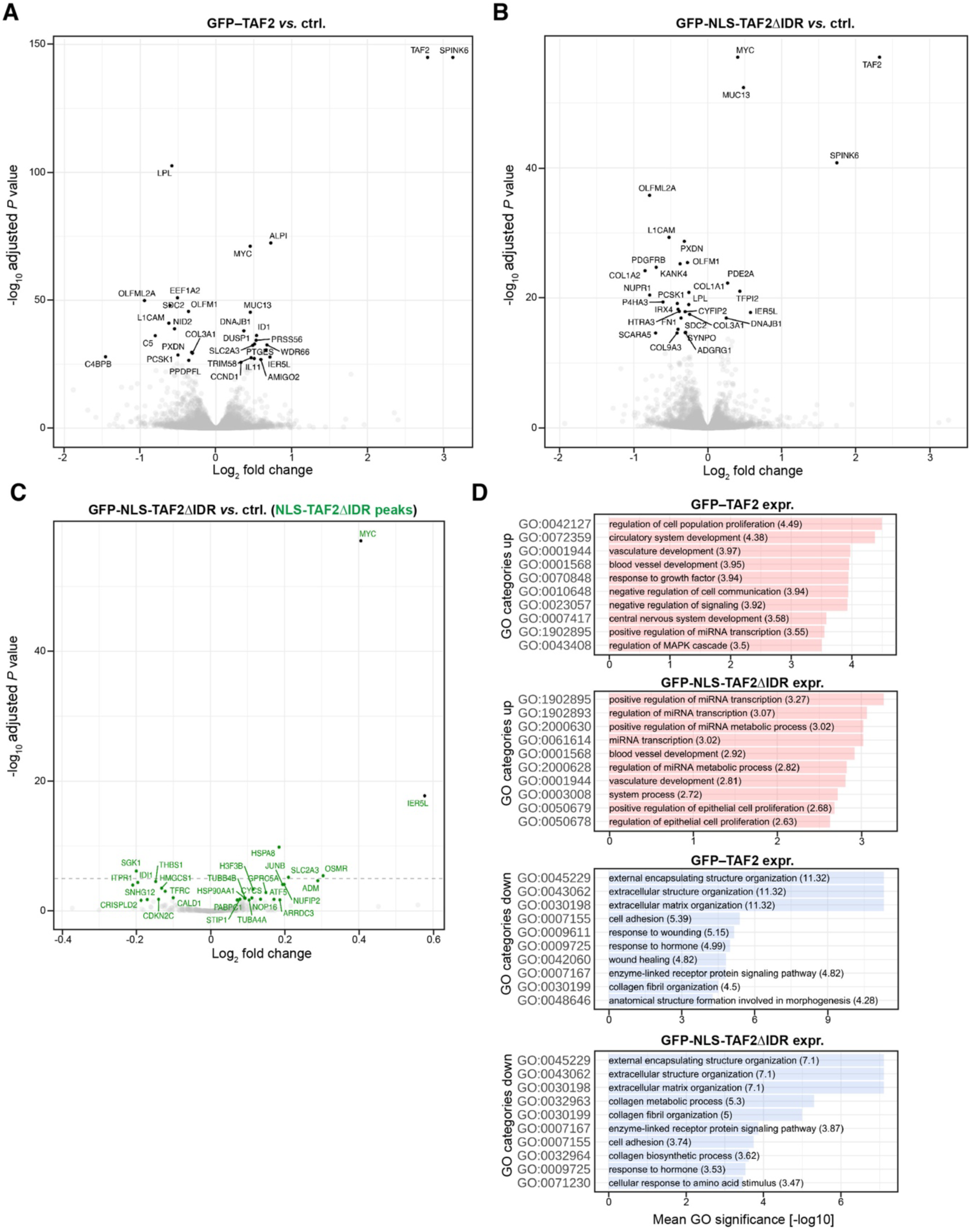
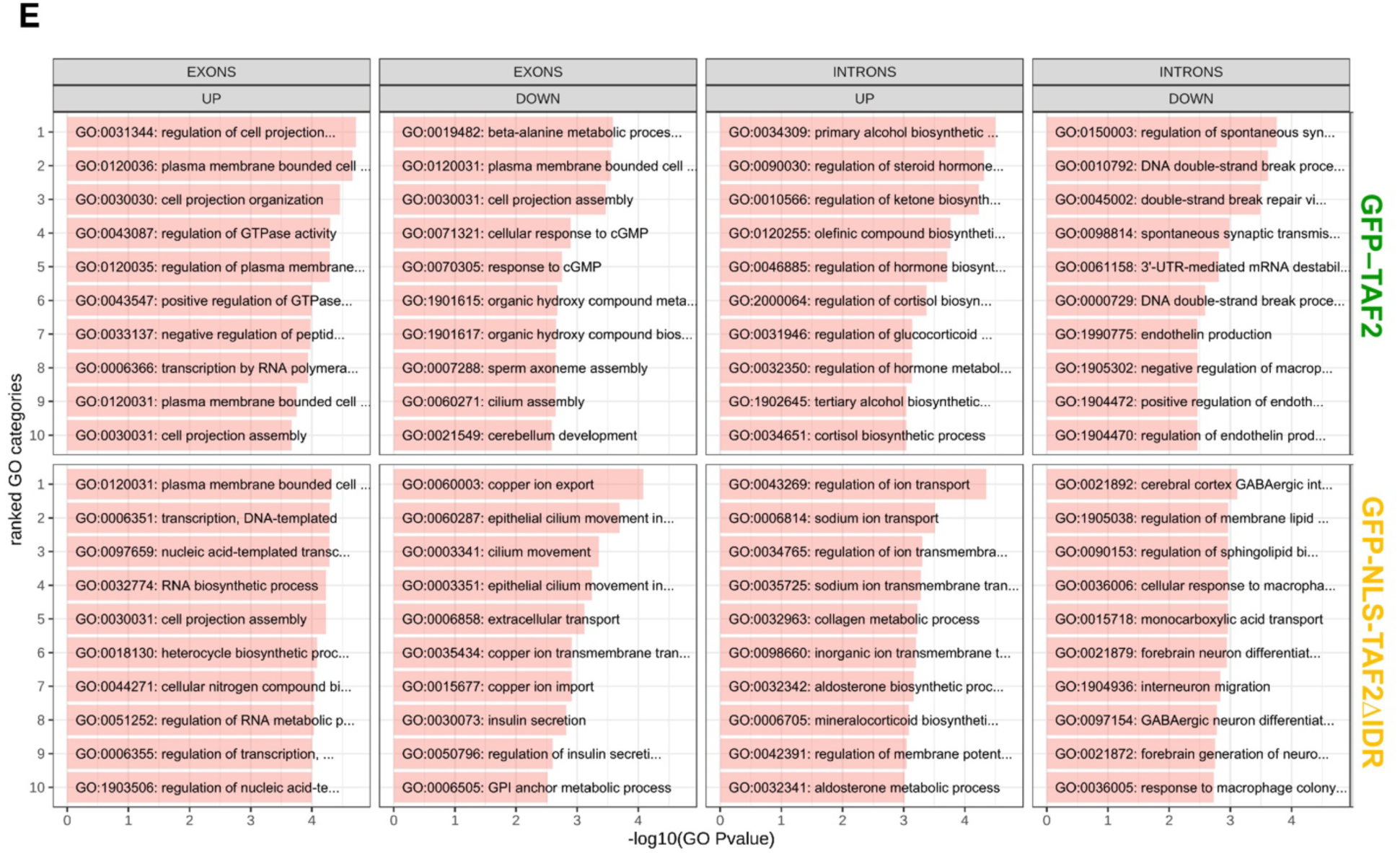
RNA-seq differential expression and gene ontology analyses. **(A)** and **(B)**, Volcano plots showing differential expression (DE) analysis results from RNA-seq data comparisons GFP–TAF2 *vs.* control (ctrl.) and GFP-NLS-TAF2ΔIDR *vs*. control (ctrl.). Positive fold changes denote up-regulated genes in GFP–TAF2 and GFP-NLS-TAF2ΔIDR expressing HeLa cells, respectively. The top 30 DE genes according to the adjusted *P* value (*P*-valadj) are labeled for each comparison. RNA-seq experiments were performed as technical triplicates. **(C)** Volcano plot showing DE results from RNA-seq data comparison GFP-NLS-TAF2ΔIDR *vs*. control (ctrl.). Displayed are 393 genes with increased GFP-NLS-TAF2ΔIDR promoter enrichment (green labels). Black dots denote genes that are among the top 30 DE genes according to *P*-valadj. **(D)** Gene ontology (GO) analysis of RNA-seq data showing the top 10 categories of biological processes of up- and downregulated genes. Mean GO *P* value represents the average of obtained *P* values after several runs of ClusterProfiler. **(E)** GO analysis of alternative splicing data performed with GOfuncR [6] for each gene group of differentially spliced exons and introns, ranked by minus log10 of the GO *P* value association. Categories are shown if the associated *P* value is below 0.01.

**Source data related to Fig. S1B.**
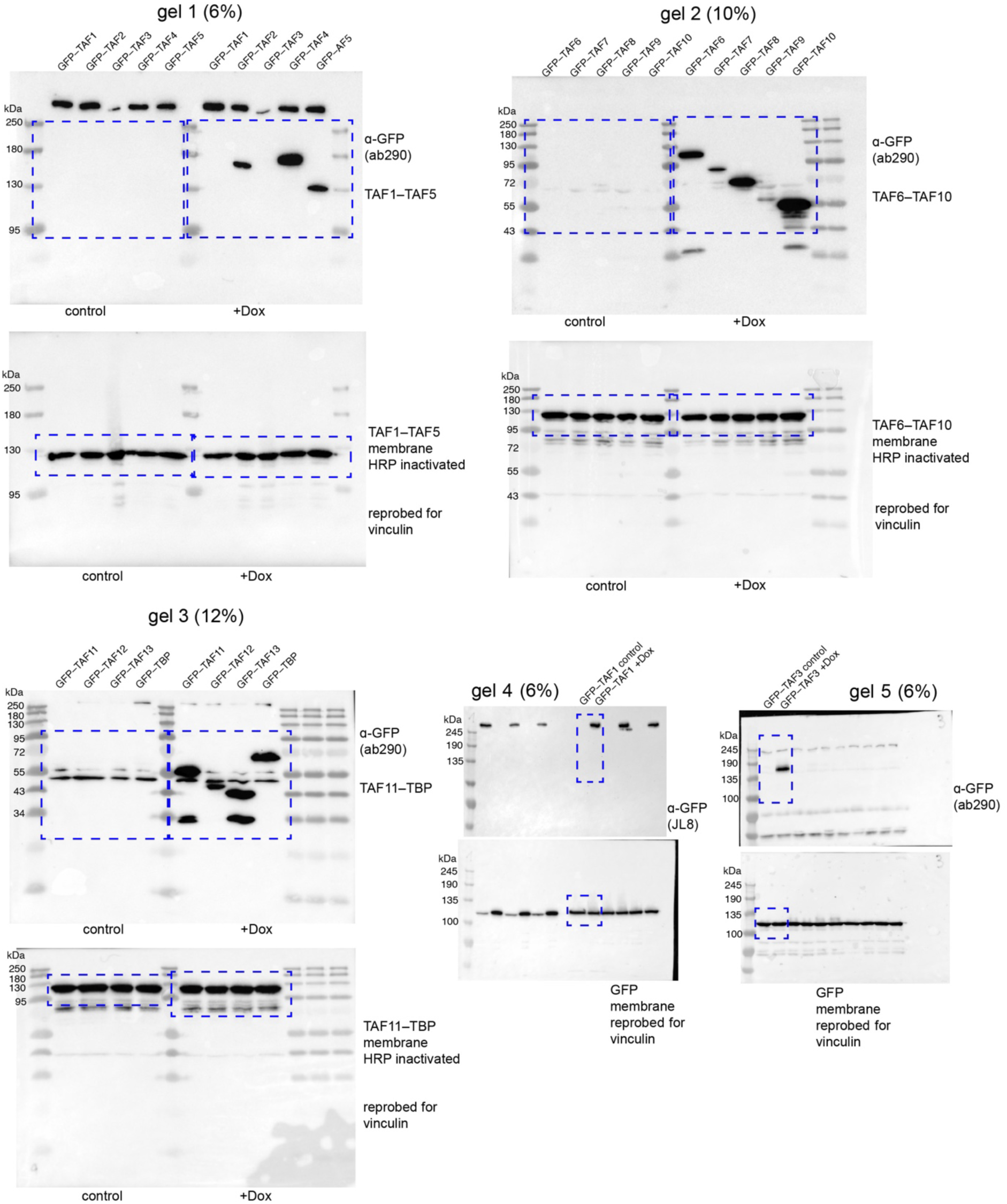

**Source data related to Fig. S3G.**
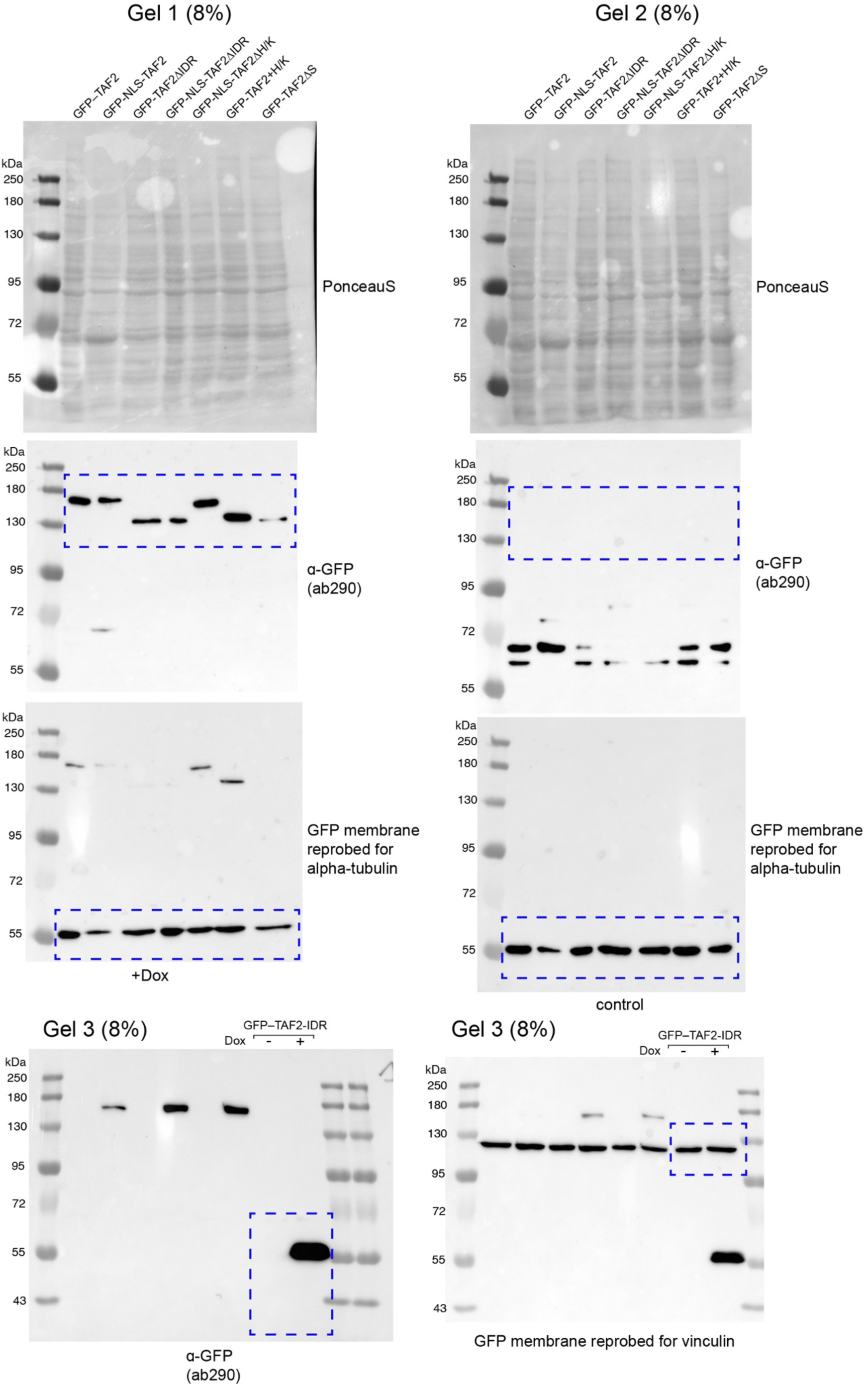

**Source data related to Fig. S4A.**
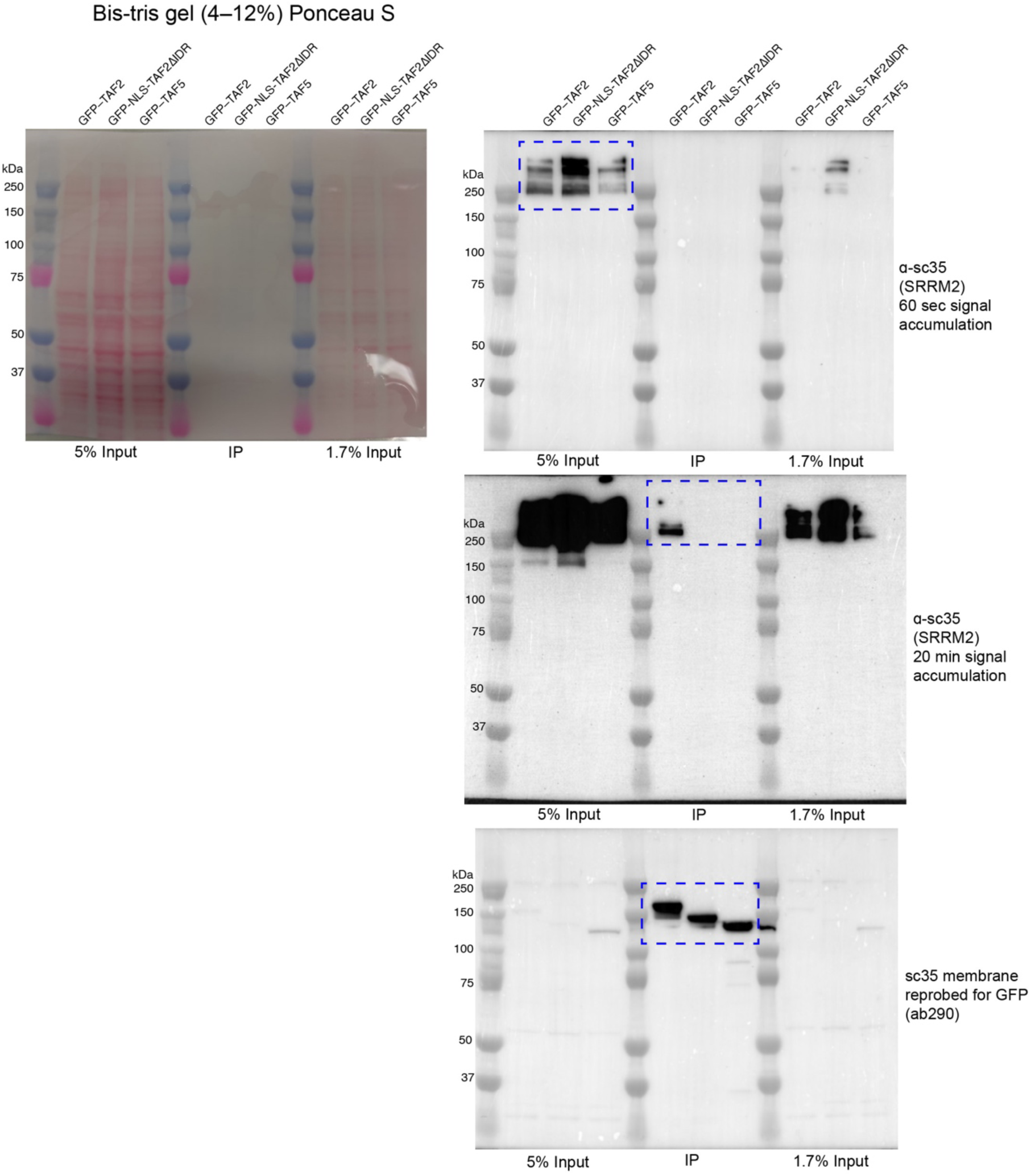

**Source data related to Fig. 4C.**
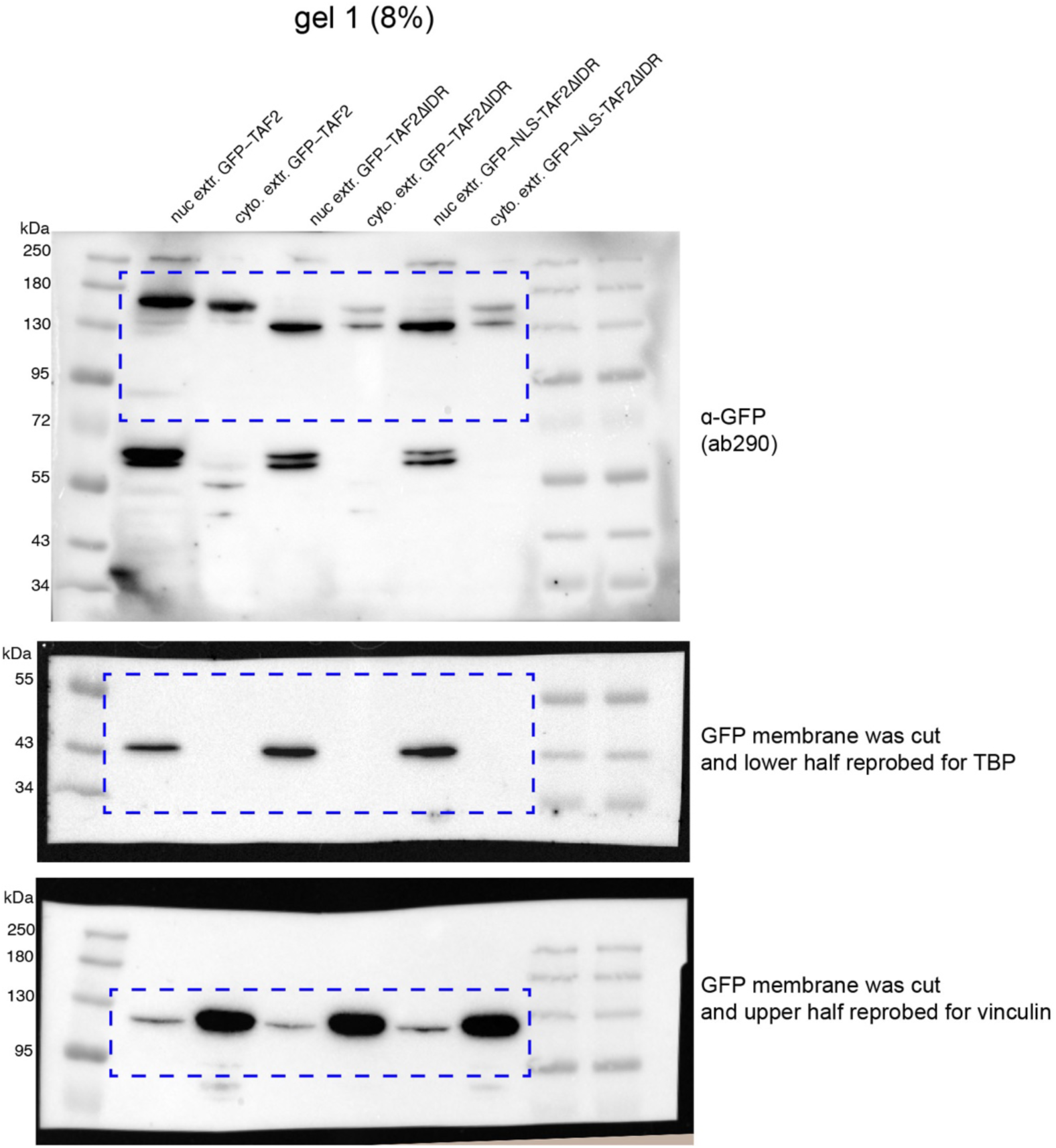

**Source data related to Fig. S4A.**
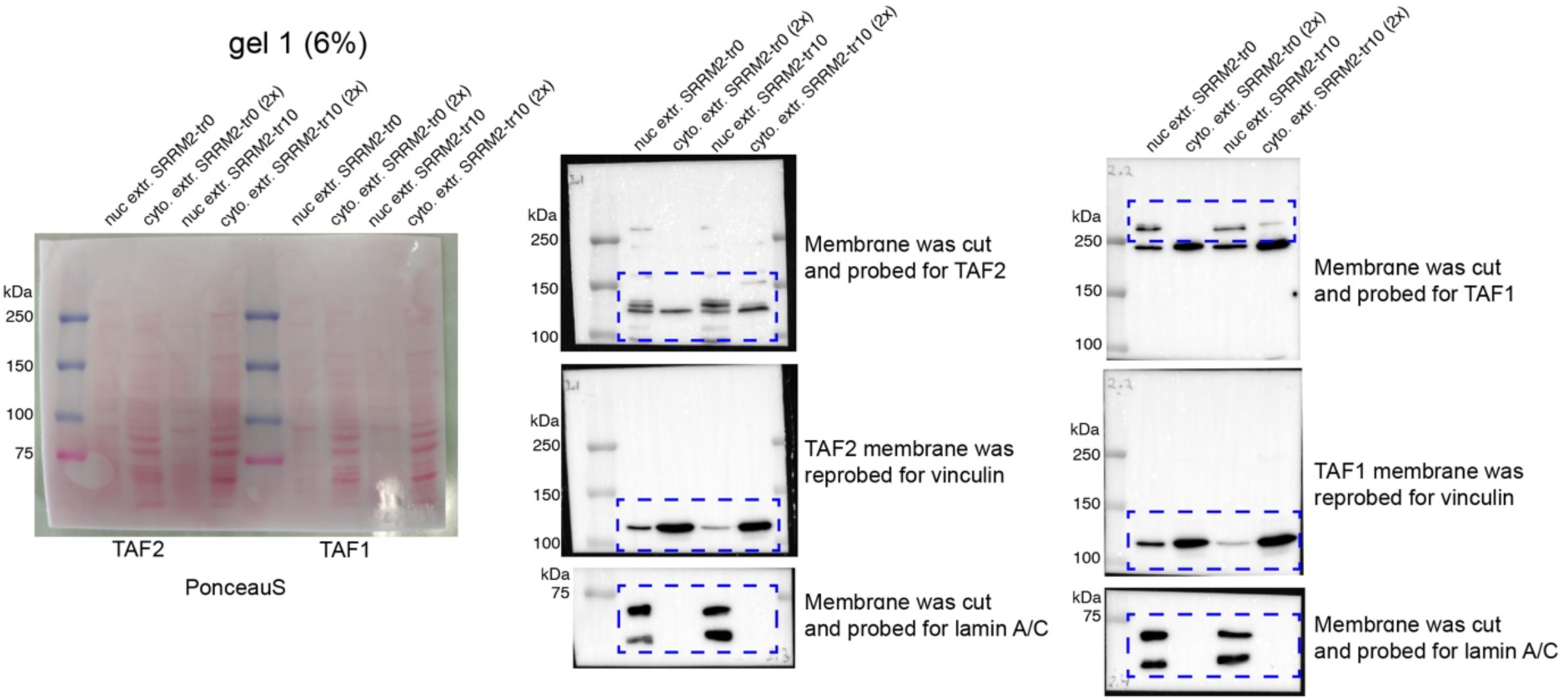

## References

1. Roeder, R.G. (2019). 50+ years of eukaryotic transcription: an expanding universe of factors and mechanisms. Nat Struct Mol Biol 26, 783–791. 10.1038/s41594-019-0287-x.

2. Bhuiyan, T., and Timmers, H.T.M. (2019). Promoter Recognition: Putting TFIID on the Spot. Trends Cell Biol 29, 752–763. 10.1016/j.tcb.2019.06.004.

3. Malik, S., and Roeder, R.G. (2023). Regulation of the RNA polymerase II pre-initiation complex by its associated coactivators. Nat Rev Genet. 10.1038/s41576-023-00630-9.

4. Orphanides, G., Lagrange, T., and Reinberg, D. (1996). The general transcription factors of RNA polymerase II. Genes Dev 10, 2657–2683.

5. Roeder, R.G. (1996). The role of general initiation factors in transcription by RNA polymerase II. Trends Biochem Sci 21, 327–335.

6. Thomas, M.C., and Chiang, C.M. (2006). The general transcription machinery and general cofactors. Crit Rev Biochem Mol Biol 41, 105–178. 10.1080/10409230600648736.

7. Allen, B.L., and Taatjes, D.J. (2015). The Mediator complex: a central integrator of transcription. Nat Rev Mol Cell Biol 16, 155–166. 10.1038/nrm3951.

8. Burley, S.K., and Roeder, R.G. (1996). Biochemistry and structural biology of transcription factor IID (TFIID). Annu Rev Biochem 65, 769–799. 10.1146/annurev.bi.65.070196.004005.

9. Cianfrocco, M.A., Kassavetis, G.A., Grob, P., Fang, J., Juven-Gershon, T., Kadonaga, J.T., and Nogales, E. (2013). Human TFIID binds to core promoter DNA in a reorganized structural state. Cell 152, 120–131. 10.1016/j.cell.2012.12.005.

10. Patel, A.B., Louder, R.K., Greber, B.J., Grunberg, S., Luo, J., Fang, J., Liu, Y., Ranish, J., Hahn, S., and Nogales, E. (2018). Structure of human TFIID and mechanism of TBP loading onto promoter DNA. Science 362. 10.1126/science.aau8872.

11. Chen, X., Qi, Y., Wu, Z., Wang, X., Li, J., Zhao, D., Hou, H., Li, Y., Yu, Z., Liu, W., et al. (2021). Structural insights into preinitiation complex assembly on core promoters. Science 372. 10.1126/science.aba8490.

12. Bernardini, A., Mukherjee, P., Scheer, E., Kamenova, I., Antonova, S., Mendoza Sanchez, P.K., Yayli, G., Morlet, B., Timmers, H.T.M., and Tora, L. (2023). Hierarchical TAF1-dependent co-translational assembly of the basal transcription factor TFIID. Nat Struct Mol Biol 30, 1141–1152. 10.1038/s41594-023-01026-3.

13. Antonova, S.V., Haffke, M., Corradini, E., Mikuciunas, M., Low, T.Y., Signor, L., van Es, R.M., Gupta, K., Scheer, E., Vos, H.R., et al. (2018). Chaperonin CCT checkpoint function in basal transcription factor TFIID assembly. Nat Struct Mol Biol 25, 1119–1127. 10.1038/s41594-018-0156-z.

14. Louder, R.K., He, Y., Lopez-Blanco, J.R., Fang, J., Chacon, P., and Nogales, E. (2016). Structure of promoter-bound TFIID and model of human pre-initiation complex assembly. Nature 531, 604–609. 10.1038/nature17394.

15. Mohan, W.S., Jr., Scheer, E., Wendling, O., Metzger, D., and Tora, L. (2003). TAF10 (TAF(II)30) is necessary for TFIID stability and early embryogenesis in mice. Mol Cell Biol 23, 4307–4318. 10.1128/MCB.23.12.4307-4318.2003.

16. Gudmundsson, S., Wilbe, M., Filipek-Gorniok, B., Molin, A.M., Ekvall, S., Johansson, J., Allalou, A., Gylje, H., Kalscheuer, V.M., Ledin, J., et al. (2019). TAF1, associated with intellectual disability in humans, is essential for embryogenesis and regulates neurodevelopmental processes in zebrafish. Sci Rep 9, 10730. 10.1038/s41598-019-46632-8.

17. Gegonne, A., Tai, X., Zhang, J., Wu, G., Zhu, J., Yoshimoto, A., Hanson, J., Cultraro, C., Chen, Q.R., Guinter, T., et al. (2012). The general transcription factor TAF7 is essential for embryonic development but not essential for the survival or differentiation of mature T cells. Mol Cell Biol 32, 1984–1997. 10.1128/MCB.06305-11.

18. Verrijzer, C.P., Yokomori, K., Chen, J.L., and Tjian, R. (1994). Drosophila TAFII150: similarity to yeast gene TSM-1 and specific binding to core promoter DNA. Science 264, 933–941. 10.1126/science.8178153.

19. Kaufmann, J., Ahrens, K., Koop, R., Smale, S.T., and Muller, R. (1998). CIF150, a human cofactor for transcription factor IID-dependent initiator function. Mol Cell Biol 18, 233–239. 10.1128/MCB.18.1.233.

20. Martinez, E., Ge, H., Tao, Y., Yuan, C.X., Palhan, V., and Roeder, R.G. (1998). Novel cofactors and TFIIA mediate functional core promoter selectivity by the human TAFII150-containing TFIID complex. Mol Cell Biol 18, 6571–6583. 10.1128/MCB.18.11.6571.

21. Kolesnikova, O., Ben-Shem, A., Luo, J., Ranish, J., Schultz, P., and Papai, G. (2018). Molecular structure of promoter-bound yeast TFIID. Nat Commun 9, 4666. 10.1038/s41467-018-07096-y.

22. Antonova, S.V., Boeren, J., Timmers, H.T.M., and Snel, B. (2019). Epigenetics and transcription regulation during eukaryotic diversification: the saga of TFIID. Genes Dev 33, 888–902. 10.1101/gad.300475.117.

23. Scheer, E., Luo, J., Bernardini, A., Ruffenach, F., Garnier, J.M., Kolb-Cheynel, I., Gupta, K., Berger, I., Ranish, J., and Tora, L. (2021). TAF8 regions important for TFIID lobe B assembly or for TAF2 interactions are required for embryonic stem cell survival. J Biol Chem 297, 101288. 10.1016/j.jbc.2021.101288.

24. Najmabadi, H., Hu, H., Garshasbi, M., Zemojtel, T., Abedini, S.S., Chen, W., Hosseini, M., Behjati, F., Haas, S., Jamali, P., et al. (2011). Deep sequencing reveals 50 novel genes for recessive cognitive disorders. Nature 478, 57–63. 10.1038/nature10423.

25. Hellman-Aharony, S., Smirin-Yosef, P., Halevy, A., Pasmanik-Chor, M., Yeheskel, A., Har-Zahav, A., Maya, I., Straussberg, R., Dahary, D., Haviv, A., et al. (2013). Microcephaly thin corpus callosum intellectual disability syndrome caused by mutated TAF2. Pediatr Neurol 49, 411–416 e411. 10.1016/j.pediatrneurol.2013.07.017.

26. Lesieur-Sebellin, M., Capri, Y., Grisval, M., Courtin, T., Burtz, A., Thevenon, J., Buratti, J., Lejeune, E., Faivre, L., and Keren, B. (2021). Phenotype associated with TAF2 biallelic mutations: A clinical description of four individuals and review of the literature. Eur J Med Genet 64, 104323. 10.1016/j.ejmg.2021.104323.

27. Parris, T.Z., Kovacs, A., Hajizadeh, S., Nemes, S., Semaan, M., Levin, M., Karlsson, P., and Helou, K. (2014). Frequent MYC coamplification and DNA hypomethylation of multiple genes on 8q in 8p11-p12-amplified breast carcinomas. Oncogenesis 3, e95. 10.1038/oncsis.2014.8.

28. Martin, J., Halenbeck, R., and Kaufmann, J. (1999). Human transcription factor hTAF(II)150 (CIF150) is involved in transcriptional regulation of cell cycle progression. Mol Cell Biol 19, 5548–5556. 10.1128/MCB.19.8.5548.

29. Zhang, Z., Boskovic, Z., Hussain, M.M., Hu, W., Inouye, C., Kim, H.J., Abole, A.K., Doud, M.K., Lewis, T.A., Koehler, A.N., et al. (2015). Chemical perturbation of an intrinsically disordered region of TFIID distinguishes two modes of transcription initiation. Elife 4. 10.7554/eLife.07777.

30. Cermakova, K., and Hodges, H.C. (2023). Interaction modules that impart specificity to disordered protein. Trends Biochem Sci 48, 477–490. 10.1016/j.tibs.2023.01.004.

31. Cumberworth, A., Lamour, G., Babu, M.M., and Gsponer, J. (2013). Promiscuity as a functional trait: intrinsically disordered regions as central players of interactomes. Biochem J 454, 361–369. 10.1042/BJ20130545.

32. Zarin, T., Strome, B., Nguyen Ba, A.N., Alberti, S., Forman-Kay, J.D., and Moses, A.M. (2019). Proteome-wide signatures of function in highly diverged intrinsically disordered regions. Elife 8. 10.7554/eLife.46883.

33. Pappu, R.V. (2020). Phase Separation-A Physical Mechanism for Organizing Information and Biochemical Reactions. Dev Cell 55, 1–3. 10.1016/j.devcel.2020.09.023.

34. Sabari, B.R., Dall’Agnese, A., and Young, R.A. (2020). Biomolecular Condensates in the Nucleus. Trends Biochem Sci 45, 961–977. 10.1016/j.tibs.2020.06.007.

35. Sabari, B.R., Dall’Agnese, A., Boija, A., Klein, I.A., Coffey, E.L., Shrinivas, K., Abraham, B.J., Hannett, N.M., Zamudio, A.V., Manteiga, J.C., et al. (2018). Coactivator condensation at super-enhancers links phase separation and gene control. Science 361. 10.1126/science.aar3958.

36. Cho, W.K., Spille, J.H., Hecht, M., Lee, C., Li, C., Grube, V., and Cisse, II (2018). Mediator and RNA polymerase II clusters associate in transcription-dependent condensates. Science 361, 412–415. 10.1126/science.aar4199.

37. Rawat, P., Boehning, M., Hummel, B., Aprile-Garcia, F., Pandit, A.S., Eisenhardt, N., Khavaran, A., Niskanen, E., Vos, S.M., Palvimo, J.J., et al. (2021). Stress-induced nuclear condensation of NELF drives transcriptional downregulation. Mol Cell 81, 1013–1026 e1011. 10.1016/j.molcel.2021.01.016.

38. Guo, C., Che, Z., Yue, J., Xie, P., Hao, S., Xie, W., Luo, Z., and Lin, C. (2020). ENL initiates multivalent phase separation of the super elongation complex (SEC) in controlling rapid transcriptional activation. Sci Adv 6, eaay4858. 10.1126/sciadv.aay4858.

39. Wan, L., Chong, S., Xuan, F., Liang, A., Cui, X., Gates, L., Carroll, T.S., Li, Y., Feng, L., Chen, G., et al. (2020). Impaired cell fate through gain-of-function mutations in a chromatin reader. Nature 577, 121–126. 10.1038/s41586-019-1842-7.

40. Palacio, M., and Taatjes, D.J. (2022). Merging Established Mechanisms with New Insights: Condensates, Hubs, and the Regulation of RNA Polymerase II Transcription. J Mol Biol 434, 167216. 10.1016/j.jmb.2021.167216.

41. Henninger, J.E., Oksuz, O., Shrinivas, K., Sagi, I., LeRoy, G., Zheng, M.M., Andrews, J.O., Zamudio, A.V., Lazaris, C., Hannett, N.M., et al. (2021). RNA-Mediated Feedback Control of Transcriptional Condensates. Cell 184, 207–225 e224. 10.1016/j.cell.2020.11.030.

42. Lewis, B.A., Das, S.K., Jha, R.K., and Levens, D. (2023). Self-assembly of promoter DNA and RNA Pol II machinery into transcriptionally active biomolecular condensates. Sci Adv 9, eadi4565. 10.1126/sciadv.adi4565.

43. Spector, D.L., and Lamond, A.I. (2011). Nuclear speckles. Cold Spring Harb Perspect Biol 3. 10.1101/cshperspect.a000646.

44. Ilik, I.A., and Aktas, T. (2021). Nuclear speckles: dynamic hubs of gene expression regulation. FEBS J. 10.1111/febs.16117.

45. Belmont, A.S. (2022). Nuclear Compartments: An Incomplete Primer to Nuclear Compartments, Bodies, and Genome Organization Relative to Nuclear Architecture. Cold Spring Harb Perspect Biol 14. 10.1101/cshperspect.a041268.

46. Faber, G.P., Nadav-Eliyahu, S., and Shav-Tal, Y. (2022). Nuclear speckles - a driving force in gene expression. J Cell Sci 135. 10.1242/jcs.259594.

47. Galganski, L., Urbanek, M.O., and Krzyzosiak, W.J. (2017). Nuclear speckles: molecular organization, biological function and role in disease. Nucleic Acids Res 45, 10350–10368. 10.1093/nar/gkx759.

48. Zhang, Q., Kota, K.P., Alam, S.G., Nickerson, J.A., Dickinson, R.B., and Lele, T.P. (2016). Coordinated Dynamics of RNA Splicing Speckles in the Nucleus. J Cell Physiol 231, 1269–1275. 10.1002/jcp.25224.

49. Marzahn, M.R., Marada, S., Lee, J., Nourse, A., Kenrick, S., Zhao, H., Ben-Nissan, G., Kolaitis, R.M., Peters, J.L., Pounds, S., et al. (2016). Higher-order oligomerization promotes localization of SPOP to liquid nuclear speckles. EMBO J 35, 1254–1275. 10.15252/embj.201593169.

50. Kim, J., Han, K.Y., Khanna, N., Ha, T., and Belmont, A.S. (2019). Nuclear speckle fusion via long-range directional motion regulates speckle morphology after transcriptional inhibition. J Cell Sci 132. 10.1242/jcs.226563.

51. Dopie, J., Sweredoski, M.J., Moradian, A., and Belmont, A.S. (2020). Tyramide signal amplification mass spectrometry (TSA-MS) ratio identifies nuclear speckle proteins. J Cell Biol 219. 10.1083/jcb.201910207.

52. Ilik, I.A., Malszycki, M., Lubke, A.K., Schade, C., Meierhofer, D., and Aktas, T. (2020). SON and SRRM2 are essential for nuclear speckle formation. Elife 9. 10.7554/eLife.60579.

53. Trowitzsch, S., Viola, C., Scheer, E., Conic, S., Chavant, V., Fournier, M., Papai, G., Ebong, I.O., Schaffitzel, C., Zou, J., et al. (2015). Cytoplasmic TAF2-TAF8-TAF10 complex provides evidence for nuclear holo-TFIID assembly from preformed submodules. Nat Commun 6, 6011. 10.1038/ncomms7011.

54. Branon, T.C., Bosch, J.A., Sanchez, A.D., Udeshi, N.D., Svinkina, T., Carr, S.A., Feldman, J.L., Perrimon, N., and Ting, A.Y. (2018). Efficient proximity labeling in living cells and organisms with TurboID. Nat Biotechnol 36, 880–887. 10.1038/nbt.4201.

55. Caudron-Herger, M., Rusin, S.F., Adamo, M.E., Seiler, J., Schmid, V.K., Barreau, E., Kettenbach, A.N., and Diederichs, S. (2019). R-DeeP: Proteome-wide and Quantitative Identification of RNA-Dependent Proteins by Density Gradient Ultracentrifugation. Mol Cell 75, 184–199 e110. 10.1016/j.molcel.2019.04.018.

56. Peng, K., Radivojac, P., Vucetic, S., Dunker, A.K., and Obradovic, Z. (2006). Length-dependent prediction of protein intrinsic disorder. BMC Bioinformatics 7, 208. 10.1186/1471-2105-7-208.

57. Jones, D.T., and Cozzetto, D. (2015). DISOPRED3: precise disordered region predictions with annotated protein-binding activity. Bioinformatics 31, 857–863. 10.1093/bioinformatics/btu744.

58. Alvarez, M., Estivill, X., and de la Luna, S. (2003). DYRK1A accumulates in splicing speckles through a novel targeting signal and induces speckle disassembly. J Cell Sci 116, 3099–3107. 10.1242/jcs.00618.

59. Salichs, E., Ledda, A., Mularoni, L., Alba, M.M., and de la Luna, S. (2009). Genome-wide analysis of histidine repeats reveals their role in the localization of human proteins to the nuclear speckles compartment. PLoS Genet 5, e1000397. 10.1371/journal.pgen.1000397.

60. Lyons, H., Veettil, R.T., Pradhan, P., Fornero, C., De La Cruz, N., Ito, K., Eppert, M., Roeder, R.G., and Sabari, B.R. (2023). Functional partitioning of transcriptional regulators by patterned charge blocks. Cell 186, 327–345 e328. 10.1016/j.cell.2022.12.013.

61. Kulak, N.A., Pichler, G., Paron, I., Nagaraj, N., and Mann, M. (2014). Minimal, encapsulated proteomic-sample processing applied to copy-number estimation in eukaryotic cells. Nat Methods 11, 319–324. 10.1038/nmeth.2834.

62. Axelrod, D., Koppel, D.E., Schlessinger, J., Elson, E., and Webb, W.W. (1976). Mobility measurement by analysis of fluorescence photobleaching recovery kinetics. Biophys J 16, 1055–1069. 10.1016/S0006-3495(76)85755-4.

63. Rai, A.K., Chen, J.X., Selbach, M., and Pelkmans, L. (2018). Kinase-controlled phase transition of membraneless organelles in mitosis. Nature 559, 211–216. 10.1038/s41586-018-0279-8.

64. Xu, S., Lai, S.K., Sim, D.Y., Ang, W.S.L., Li, H.Y., and Roca, X. (2022). SRRM2 organizes splicing condensates to regulate alternative splicing. Nucleic Acids Res 50, 8599–8614. 10.1093/nar/gkac669.

65. Jia, X., and Sun, C. (2018). Structural dynamics of the N-terminal domain and the Switch loop of Prp8 during spliceosome assembly and activation. Nucleic Acids Res 46, 3833–3840. 10.1093/nar/gky242.

66. Blencowe, B.J., Bauren, G., Eldridge, A.G., Issner, R., Nickerson, J.A., Rosonina, E., and Sharp, P.A. (2000). The SRm160/300 splicing coactivator subunits. RNA 6, 111–120. 10.1017/s1355838200991982.

67. Eldridge, A.G., Li, Y., Sharp, P.A., and Blencowe, B.J. (1999). The SRm160/300 splicing coactivator is required for exon-enhancer function. Proc Natl Acad Sci U S A 96, 6125–6130. 10.1073/pnas.96.11.6125.

68. Nizamuddin, S., Koidl, S., Bhuiyan, T., Werner, T.V., Biniossek, M.L., Bonvin, A., Lassmann, S., and Timmers, H. (2021). Integrating quantitative proteomics with accurate genome profiling of transcription factors by greenCUT&RUN. Nucleic Acids Res 49, e49. 10.1093/nar/gkab038.

69. Koidl, S., and Timmers, H.T.M. (2021). greenCUT&RUN: Efficient Genomic Profiling of GFP-Tagged Transcription Factors and Chromatin Regulators. Curr Protoc 1, e266. 10.1002/cpz1.266.

70. Chen, X., Yin, X., Li, J., Wu, Z., Qi, Y., Wang, X., Liu, W., and Xu, Y. (2021). Structures of the human Mediator and Mediator-bound preinitiation complex. Science 372. 10.1126/science.abg0635.

71. Papai, G., Tripathi, M.K., Ruhlmann, C., Werten, S., Crucifix, C., Weil, P.A., and Schultz, P. (2009). Mapping the initiator binding Taf2 subunit in the structure of hydrated yeast TFIID. Structure 17, 363–373. 10.1016/j.str.2009.01.006.

72. Portz, B., Lu, F., Gibbs, E.B., Mayfield, J.E., Rachel Mehaffey, M., Zhang, Y.J., Brodbelt, J.S., Showalter, S.A., and Gilmour, D.S. (2017). Structural heterogeneity in the intrinsically disordered RNA polymerase II C-terminal domain. Nat Commun 8, 15231. 10.1038/ncomms15231.

73. Boehning, M., Dugast-Darzacq, C., Rankovic, M., Hansen, A.S., Yu, T., Marie-Nelly, H., McSwiggen, D.T., Kokic, G., Dailey, G.M., Cramer, P., et al. (2018). RNA polymerase II clustering through carboxy-terminal domain phase separation. Nat Struct Mol Biol 25, 833–840. 10.1038/s41594-018-0112-y.

74. Guo, Y.E., Manteiga, J.C., Henninger, J.E., Sabari, B.R., Dall’Agnese, A., Hannett, N.M., Spille, J.H., Afeyan, L.K., Zamudio, A.V., Shrinivas, K., et al. (2019). Pol II phosphorylation regulates a switch between transcriptional and splicing condensates. Nature 572, 543–548. 10.1038/s41586-019-1464-0.

75. Hornbeck, P.V., Zhang, B., Murray, B., Kornhauser, J.M., Latham, V., and Skrzypek, E. (2015). PhosphoSitePlus, 2014: mutations, PTMs and recalibrations. Nucleic Acids Res 43, D512–520. 10.1093/nar/gku1267.

76. Yu, R., Roseman, S., Siegenfeld, A.P., Nguyen, S.C., Joyce, E.F., Liau, B.B., Krantz, I.D., Alexander, K.A., and Berger, S.L. (2023). CTCF/cohesin organize the ground state of chromatin-nuclear speckle association. bioRxiv. 10.1101/2023.07.22.550178.

77. Alexander, K.A., Cote, A., Nguyen, S.C., Zhang, L., Gholamalamdari, O., Agudelo-Garcia, P., Lin-Shiao, E., Tanim, K.M.A., Lim, J., Biddle, N., et al. (2021). p53 mediates target gene association with nuclear speckles for amplified RNA expression. Mol Cell 81, 1666–1681 e1666. 10.1016/j.molcel.2021.03.006.

78. Greig, J.A., Nguyen, T.A., Lee, M., Holehouse, A.S., Posey, A.E., Pappu, R.V., and Jedd, G. (2020). Arginine-Enriched Mixed-Charge Domains Provide Cohesion for Nuclear Speckle Condensation. Mol Cell 77, 1237–1250 e1234. 10.1016/j.molcel.2020.01.025.

79. Oksuz, O., Henninger, J.E., Warneford-Thomson, R., Zheng, M.M., Erb, H., Vancura, A., Overholt, K.J., Hawken, S.W., Banani, S.F., Lauman, R., et al. (2023). Transcription factors interact with RNA to regulate genes. Mol Cell 83, 2449–2463 e2413. 10.1016/j.molcel.2023.06.012.

80. Herrmann, C.H., and Mancini, M.A. (2001). The Cdk9 and cyclin T subunits of TAK/P-TEFb localize to splicing factor-rich nuclear speckle regions. J Cell Sci 114, 1491–1503. 10.1242/jcs.114.8.1491.

81. Lu, H., Yu, D., Hansen, A.S., Ganguly, S., Liu, R., Heckert, A., Darzacq, X., and Zhou, Q. (2018). Phase-separation mechanism for C-terminal hyperphosphorylation of RNA polymerase II. Nature 558, 318–323. 10.1038/s41586-018-0174-3.

82. El-Saafin, F., Curry, C., Ye, T., Garnier, J.M., Kolb-Cheynel, I., Stierle, M., Downer, N.L., Dixon, M.P., Negroni, L., Berger, I., et al. (2018). Homozygous TAF8 mutation in a patient with intellectual disability results in undetectable TAF8 protein, but preserved RNA polymerase II transcription. Hum Mol Genet 27, 2171–2186. 10.1093/hmg/ddy126.

83. Jacq, X., Brou, C., Lutz, Y., Davidson, I., Chambon, P., and Tora, L. (1994). Human TAFII30 is present in a distinct TFIID complex and is required for transcriptional activation by the estrogen receptor. Cell 79, 107–117. 10.1016/0092-8674(94)90404-9.

84. Hisler, V., Bardot, P., Detilleux, D., Stierle, M., Sanchez, E.G., Richard, C., Arab, L.H., Ehrhard, C., Morlet, B., Hadzhiev, Y., et al. (2023). RNA polymerase II transcription with partially assembled TFIID complexes. bioRxiv. 10.1101/2023.11.27.567046.

85. Damgaard, C.K., Kahns, S., Lykke-Andersen, S., Nielsen, A.L., Jensen, T.H., and Kjems, J. (2008). A 5’ splice site enhances the recruitment of basal transcription initiation factors in vivo. Mol Cell 29, 271–278. 10.1016/j.molcel.2007.11.035.

86. Dantonel, J.C., Murthy, K.G., Manley, J.L., and Tora, L. (1997). Transcription factor TFIID recruits factor CPSF for formation of 3’ end of mRNA. Nature 389, 399–402. 10.1038/38763.

87. Cui, H., Diedrich, J.K., Wu, D.C., Lim, J.J., Nottingham, R.M., Moresco, J.J., Yates, J.R., 3rd, Blencowe, B.J., Lambowitz, A.M., and Schimmel, P. (2023). Arg-tRNA synthetase links inflammatory metabolism to RNA splicing and nuclear trafficking via SRRM2. Nat Cell Biol 25, 592–603. 10.1038/s41556-023-01118-8.

88. Cuinat, S., Nizon, M., Isidor, B., Stegmann, A., van Jaarsveld, R.H., van Gassen, K.L., van der Smagt, J.J., Volker-Touw, C.M.L., Holwerda, S.J.B., Terhal, P.A., et al. (2022). Loss-of-function variants in SRRM2 cause a neurodevelopmental disorder. Genet Med 24, 1774–1780. 10.1016/j.gim.2022.04.011.

89. Lester, E., Ooi, F.K., Bakkar, N., Ayers, J., Woerman, A.L., Wheeler, J., Bowser, R., Carlson, G.A., Prusiner, S.B., and Parker, R. (2021). Tau aggregates are RNA-protein assemblies that mislocalize multiple nuclear speckle components. Neuron 109, 1675–1691 e1679. 10.1016/j.neuron.2021.03.026.

90. Shehadeh, L.A., Yu, K., Wang, L., Guevara, A., Singer, C., Vance, J., and Papapetropoulos, S. (2010). SRRM2, a potential blood biomarker revealing high alternative splicing in Parkinson’s disease. PLoS One 5, e9104. 10.1371/journal.pone.0009104.

91. van Nuland, R., Schram, A.W., van Schaik, F.M., Jansen, P.W., Vermeulen, M., and Marc Timmers, H.T. (2013). Multivalent engagement of TFIID to nucleosomes. PLoS One 8, e73495. 10.1371/journal.pone.0073495.

92. van Nuland, R., Smits, A.H., Pallaki, P., Jansen, P.W., Vermeulen, M., and Timmers, H.T. (2013). Quantitative dissection and stoichiometry determination of the human SET1/MLL histone methyltransferase complexes. Mol Cell Biol 33, 2067–2077. 10.1128/MCB.01742-12.

93. Capponi, S., Stoffler, N., Irimia, M., Van Schaik, F.M.A., Ondik, M.M., Biniossek, M.L., Lehmann, L., Mitschke, J., Vermunt, M.W., Creyghton, M.P., et al. (2020). Neuronal-specific microexon splicing of TAF1 mRNA is directly regulated by SRRM4/nSR100. RNA Biol 17, 62–74. 10.1080/15476286.2019.1667214.

94. Phair, R.D., Gorski, S.A., and Misteli, T. (2004). Measurement of dynamic protein binding to chromatin in vivo, using photobleaching microscopy. Methods Enzymol 375, 393–414. 10.1016/s0076-6879(03)75025-3.

95. Koulouras, G., Panagopoulos, A., Rapsomaniki, M.A., Giakoumakis, N.N., Taraviras, S., and Lygerou, Z. (2018). EasyFRAP-web: a web-based tool for the analysis of fluorescence recovery after photobleaching data. Nucleic Acids Res 46, W467–W472. 10.1093/nar/gky508.

96. Carey, M.F., Peterson, C.L., and Smale, S.T. (2009). Dignam and Roeder nuclear extract preparation. Cold Spring Harb Protoc 2009, pdb prot5330. 10.1101/pdb.prot5330.

97. Spruijt, C.G., Baymaz, H.I., and Vermeulen, M. (2013). Identifying specific protein-DNA interactions using SILAC-based quantitative proteomics. Methods Mol Biol 977, 137–157. 10.1007/978-1-62703-284-1_11.

98. Tyanova, S., Temu, T., and Cox, J. (2016). The MaxQuant computational platform for mass spectrometry-based shotgun proteomics. Nat Protoc 11, 2301–2319. 10.1038/nprot.2016.136.

99. Schwanhausser, B., Busse, D., Li, N., Dittmar, G., Schuchhardt, J., Wolf, J., Chen, W., and Selbach, M. (2011). Global quantification of mammalian gene expression control. Nature 473, 337–342. 10.1038/nature10098.

100. Perez-Riverol, Y., Bai, J., Bandla, C., Garcia-Seisdedos, D., Hewapathirana, S., Kamatchinathan, S., Kundu, D.J., Prakash, A., Frericks-Zipper, A., Eisenacher, M., et al. (2022). The PRIDE database resources in 2022: a hub for mass spectrometry-based proteomics evidences. Nucleic Acids Res 50, D543–D552. 10.1093/nar/gkab1038.

101. Tyanova, S., Temu, T., Sinitcyn, P., Carlson, A., Hein, M.Y., Geiger, T., Mann, M., and Cox, J. (2016). The Perseus computational platform for comprehensive analysis of (prote)omics data. Nat Methods 13, 731–740. 10.1038/nmeth.3901.

102. Skene, P.J., Henikoff, J.G., and Henikoff, S. (2018). Targeted in situ genome-wide profiling with high efficiency for low cell numbers. Nat Protoc 13, 1006–1019. 10.1038/nprot.2018.015.

103. Bardot, P., Vincent, S.D., Fournier, M., Hubaud, A., Joint, M., Tora, L., and Pourquie, O. (2017). The TAF10-containing TFIID and SAGA transcriptional complexes are dispensable for early somitogenesis in the mouse embryo. Development 144, 3808–3818. 10.1242/dev.146902.

104. Nguyen Ba, A.N., Pogoutse, A., Provart, N., and Moses, A.M. (2009). NLStradamus: a simple Hidden Markov Model for nuclear localization signal prediction. BMC Bioinformatics 10, 202. 10.1186/1471-2105-10-202.

105. Heinz, S., Benner, C., Spann, N., Bertolino, E., Lin, Y.C., Laslo, P., Cheng, J.X., Murre, C., Singh, H., and Glass, C.K. (2010). Simple combinations of lineage-determining transcription factors prime cis-regulatory elements required for macrophage and B cell identities. Mol Cell 38, 576–589. 10.1016/j.molcel.2010.05.004.

106. Afgan, E., Baker, D., van den Beek, M., Blankenberg, D., Bouvier, D., Cech, M., Chilton, J., Clements, D., Coraor, N., Eberhard, C., et al. (2016). The Galaxy platform for accessible, reproducible and collaborative biomedical analyses: 2016 update. Nucleic Acids Res 44, W3–W10. 10.1093/nar/gkw343.

107. Ramirez, F., Dundar, F., Diehl, S., Gruning, B.A., and Manke, T. (2014). deepTools: a flexible platform for exploring deep-sequencing data. Nucleic Acids Res 42, W187–191. 10.1093/nar/gku365.

108. Liao, Y., Smyth, G.K., and Shi, W. (2019). The R package Rsubread is easier, faster, cheaper and better for alignment and quantification of RNA sequencing reads. Nucleic Acids Res 47, e47. 10.1093/nar/gkz114.

109. Mehmet Tekman, G.-J.K. (2023). GitHub - mtekman/rnaseqhelper: RNAseq Helper.

110. Love, M.I., Huber, W., and Anders, S. (2014). Moderated estimation of fold change and dispersion for RNA-seq data with DESeq2. Genome Biol 15, 550. 10.1186/s13059-014-0550-8.

111. Prunotto, A., Stevenson, B.J., Berthonneche, C., Schupfer, F., Beckmann, J.S., Maurer, F., and Bergmann, S. (2016). RNAseq analysis of heart tissue from mice treated with atenolol and isoproterenol reveals a reciprocal transcriptional response. BMC Genomics 17, 717. 10.1186/s12864-016-3059-6.

112. Tapial, J., Ha, K.C.H., Sterne-Weiler, T., Gohr, A., Braunschweig, U., Hermoso-Pulido, A., Quesnel-Vallieres, M., Permanyer, J., Sodaei, R., Marquez, Y., et al. (2017). An atlas of alternative splicing profiles and functional associations reveals new regulatory programs and genes that simultaneously express multiple major isoforms. Genome Res 27, 1759–1768. 10.1101/gr.220962.117.

113. Zhao, M., Mantel, I., Gelize, E., Li, X., Xie, X., Arboleda, A., Seminel, M., Levy-Boukris, R., Dernigoghossian, M., Prunotto, A., et al. (2019). Mineralocorticoid receptor antagonism limits experimental choroidal neovascularization and structural changes associated with neovascular age-related macular degeneration. Nat Commun 10, 369. 10.1038/s41467-018-08125-6.

114. Wu, T., Hu, E., Xu, S., Chen, M., Guo, P., Dai, Z., Feng, T., Zhou, L., Tang, W., Zhan, L., et al. (2021). clusterProfiler 4.0: A universal enrichment tool for interpreting omics data. Innovation (N Y) 2, 100141. 10.1016/j.xinn.2021.100141.

115. Grote, S. (2023). GOfuncR: Gene ontology enrichment using FUNC. R package version 1.22.0. doi:10.18129/B9.bioc.GOfuncR.

116. Hansen, K.D., Langmead, B., and Irizarry, R.A. (2012). BSmooth: from whole genome bisulfite sequencing reads to differentially methylated regions. Genome Biol 13, R83. 10.1186/gb-2012-13-10-r83.

117. Korthauer, K., Chakraborty, S., Benjamini, Y., and Irizarry, R.A. (2019). Detection and accurate false discovery rate control of differentially methylated regions from whole genome bisulfite sequencing. Biostatistics 20, 367–383. 10.1093/biostatistics/kxy007.

118. Laufer, B.I., Hwang, H., Jianu, J.M., Mordaunt, C.E., Korf, I.F., Hertz-Picciotto, I., and LaSalle, J.M. (2021). Low-pass whole genome bisulfite sequencing of neonatal dried blood spots identifies a role for RUNX1 in Down syndrome DNA methylation profiles. Hum Mol Genet 29, 3465–3476. 10.1093/hmg/ddaa218.

119. Waterhouse, A.M., Procter, J.B., Martin, D.M., Clamp, M., and Barton, G.J. (2009). Jalview Version 2--a multiple sequence alignment editor and analysis workbench. Bioinformatics 25, 1189–1191. 10.1093/bioinformatics/btp033.

120. Madeira, F., Park, Y.M., Lee, J., Buso, N., Gur, T., Madhusoodanan, N., Basutkar, P., Tivey, A.R.N., Potter, S.C., Finn, R.D., and Lopez, R. (2019). The EMBL-EBI search and sequence analysis tools APIs in 2019. Nucleic Acids Res 47, W636–W641. 10.1093/nar/gkz268.

## Supplemental references

[1.] Caudron-Herger, M., Rusin, S.F., Adamo, M.E., Seiler, J., Schmid, V.K., Barreau, E., Kettenbach, A.N., and Diederichs, S. (2019). R-DeeP: Proteome-wide and Quantitative Identification of RNA-Dependent Proteins by Density Gradient Ultracentrifugation. Mol Cell 75, 184–199 e110. 10.1016/j.molcel.2019.04.018.

[2.] Peng, K., Radivojac, P., Vucetic, S., Dunker, A.K., and Obradovic, Z. (2006). Length-dependent prediction of protein intrinsic disorder. BMC Bioinformatics 7, 208. 10.1186/1471-2105-7-208.

[3.] Jones, D.T., and Cozzetto, D. (2015). DISOPRED3: precise disordered region predictions with annotated protein-binding activity. Bioinformatics 31, 857–863. 10.1093/bioinformatics/btu744.

[4.] Waterhouse, A.M., Procter, J.B., Martin, D.M., Clamp, M., and Barton, G.J. (2009). Jalview Version 2--a multiple sequence alignment editor and analysis workbench. Bioinformatics 25, 1189–1191. 10.1093/bioinformatics/btp033.

[5.] Ilik, I.A., Malszycki, M., Lubke, A.K., Schade, C., Meierhofer, D., and Aktas, T. (2020). SON and SRRM2 are essential for nuclear speckle formation. Elife 9. 10.7554/eLife.60579.

[6.] Grote, S. (2023). GOfuncR: Gene ontology enrichment using FUNC. R package version 1.22.0. doi:10.18129/B9.bioc.GOfuncR.

